# diagFDR: Assessing False Discovery Rate Claims in Proteomics with Scope, Calibration and Stability Diagnostics

**DOI:** 10.64898/2026.04.16.718468

**Authors:** Marie Chion, Alexandre Godmer, Thibaut Douché, Mariette Matondo, Quentin Giai Gianetto

**Affiliations:** Medical Research Council Biostatistics Unit, University of Cambridge, Cambridge, United Kingdom; Sorbonne Université, CNRS, Inserm, Centre d’Immunologie et des Maladies Infectieuses, CIMI, Paris, F-75013, France; AP-HP, Sorbonne Université, Hôpital Pitié-Salpêtrière, Centre National de Référence des Mycobactéries et de la Résistance des Mycobactéries aux Antituberculeux, Paris, France; Institut Pasteur, Université Paris Cité, Proteomics Platform, Mass Spectrometry for Biology Unit, UAR CNRS 2024, Paris, France; Institut Pasteur, Université Paris Cité, Bioinformatics and Biostatistics HUB, Paris, France

**Keywords:** false discovery rate, target–decoy competition, proteomics, data-independent acquisition, entrapment, reproducibility

## Abstract

In mass spectrometry (MS)-based proteomics, false discovery rates (FDRs) for peptide and protein identifications are estimated by comparing target and decoy matches. Yet published studies rarely give enough information to audit a reported “1% FDR”. In a convenience sample of recent data-independent acquisition (DIA) studies, none reported the number of accepted decoys or how target/decoy labels were assigned to protein groups, and few specified whether thresholds were applied per run or across the experiment. We propose a framework that separates three questions: is the claim fully specified; is it consistent with the exported decoys; and is it supported by independent evidence, such as an entrapment search? We define scope, calibration and stability diagnostics computable from decoy-inclusive exports. In a DIA-NN dataset, filtering on run-wise q-values alone gave a list with a recomputed decoy ratio of 2.2% at a nominal 1% and a Jaccard overlap of 0.93 with the experiment-wide list; the added identifications had a twelvefold higher raw entrapment-match rate. Four protein-group labelling rules changed the protein-level decoy ratio by a factor of 1.4. We derive and verify by simulation a benchmark for the dispersion of the realised false discovery proportion (FDP) around the FDR, close to 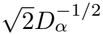 with *D*_*α*_ the number of accepted decoys. Internal diagnostics did not detect simulated decoy misspecification that raised the realised FDP up to sevenfold. We propose a five-point reporting standard and provide the R package diagFDR and its Python port diagfdr, which implement the diagnostics, label rules, entrapment estimators and a reviewer-facing report.

## 1 Introduction

Almost every proteomics study reports its identifications “at 1% FDR”. The number is produced by the target–decoy strategy [1, 2, 3]: the search is run against the real (target) sequences and against decoy sequences that cannot be in the sample, and the proportion of decoys among the accepted matches is used to estimate the proportion of false target matches. The idea is simple, but the resulting number has become hard to interpret. Early recommendations asked authors to say which list and which unit an error rate applies to [4]; the terms are still used loosely [5]. The false discovery proportion (FDP) is the fraction of false identifications actually present in a list, while the FDR is its expectation over repeated experiments [6]; q-values are meant to control the FDR of a list, under the assumptions of the procedure that produced them, whereas posterior error probabilities (PEPs) are statements about single matches [2]. In multi-run data-independent acquisition (DIA) even the “match” is ambiguous: co-fragmentation makes the correspondence between spectra and peptides many-to-many, so software scores chromatographic features against decoy features [7, 8, 9], errors propagate from precursors to proteins [10], and entrapment experiments show that DIA tools often exceed their nominal error rate [11]. We separate three questions that a reader may ask about an FDR claim:

(A1) **Is the claim fully specified?** The unit controlled (precursor, peptide, protein group), the scope of the threshold (one run or the whole experiment), the decoy construction, the rule that aggregates exported rows into one reported unit, the rule that gives that unit a target or decoy label, and the quantity thresholded should all be stated, so that the calculation can be reconstructed.

(A2) **Is the claim consistent with the exported decoys?** Given the decoys the software produced, the reported confidence values should reproduce the stated definitions, PEPs should agree with the error frequencies implied by the decoys, and the number of accepted decoys should be reported to assess the variability of the realised FDP under the stated assumptions.

(A3) **Is the claim supported by independent evidence?** Internal consistency does not show that the decoys behave like incorrect target matches; assessing this requires evidence independent of the decoys, in practice an entrapment analysis, which rests on assumptions of its own (§3.5).

A1 and A2 separate a value a reader can verify from one accepted on trust. Section 2 shows that the studies we audited rarely allow either, although the software computes everything needed. Section 3 defines diagnostics for scope, calibration and stability (Figure 1) and the entrapment estimators that address A3; Section 4 evaluates them on simulated data with known truth, then on DIA-NN and MS^2^Rescore outputs. The diagnostics build on scope ambiguity [4], decoy sparsity after strict filtering [12], target/decoy score quality control [13] and entrapment validation [11], and are implemented in the R package diagFDR on CRAN (Figure 2).

**Figure 1:**
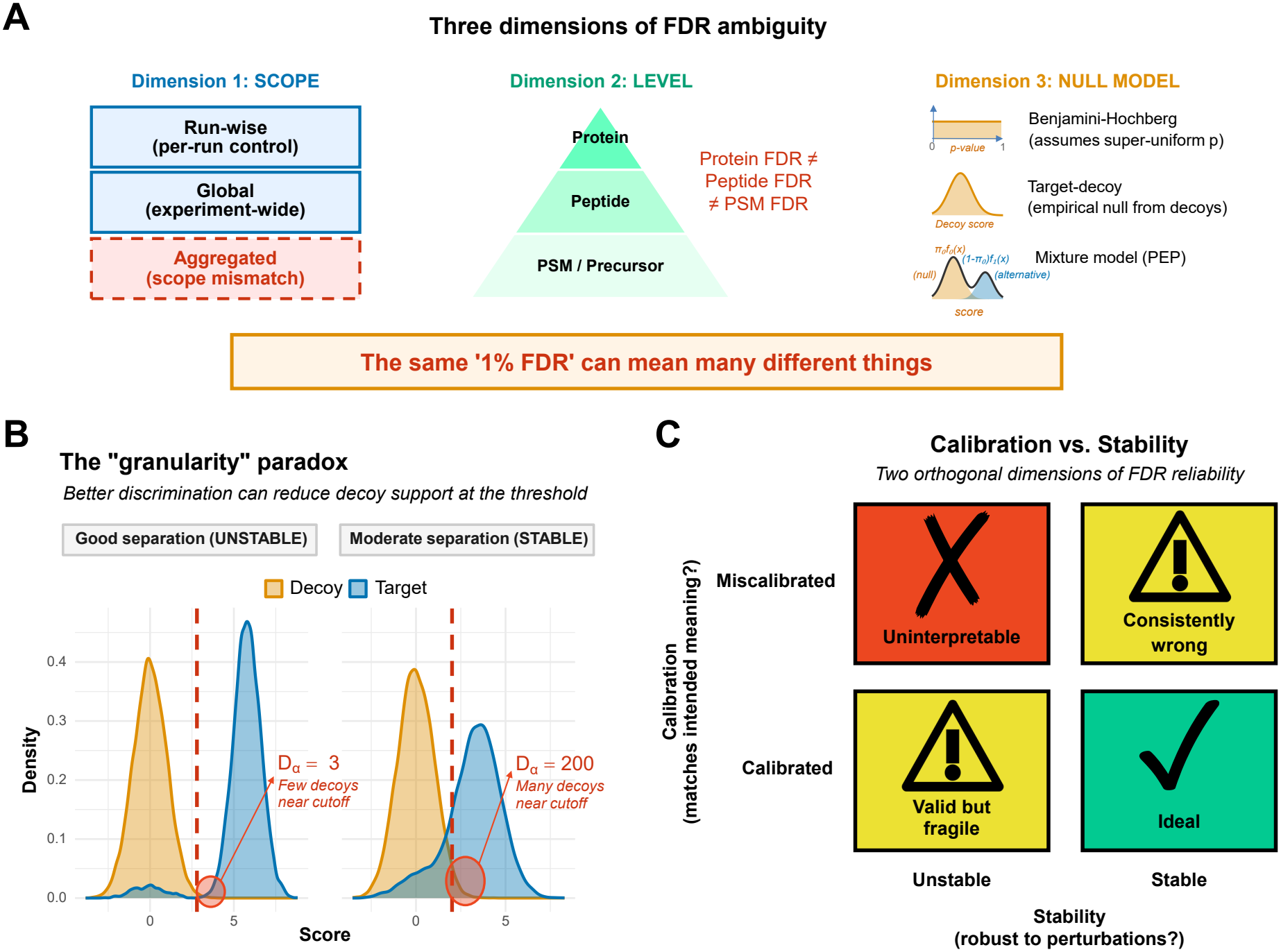
Three properties of an auditable FDR claim. (A) Dimensions along which nominally identical FDR claims diverge: the scope of the threshold, the level at which it is applied, and the null model. (B) Decoy tail granularity: at stringent cutoffs so few decoys remain that the realised false discovery proportion becomes more variable, with the variability benchmarked by Proposition 1; better scoring thins the decoy tail at a fixed sensitivity (the “granularity paradox”, which applies in that comparison only) but thickens it at a fixed nominal level, under the conditions given in Propositions S1 and S2 (§3.4). (C) FDR reliability as a joint function of calibration and stability. *Alt text:* Three-panel schematic. Panel A lists the dimensions along which identical FDR claims diverge. Panel B shows decoy counts thinning as the threshold becomes stringent, with the spread of realised error widening. Panel C plots reliability as a joint function of calibration and stability.

**Figure 2:**
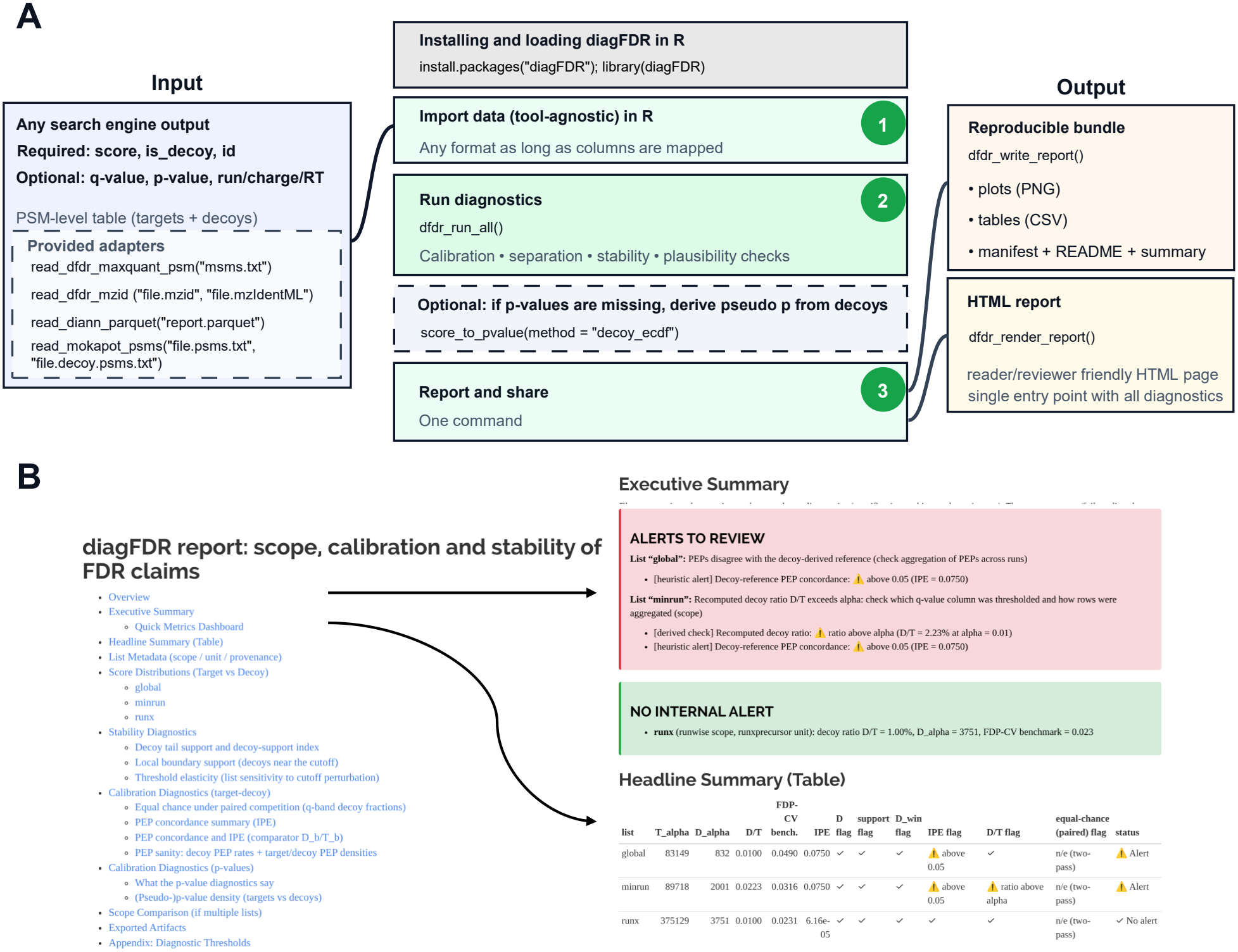
The diagFDR workflow. (A) A search output exported as a target/decoy table (`score, is_decoy`, `id` required; q-values, PEPs and run-level covariates each required by the diagnostics that use them; external validation requires entrapment or ground-truth labels) is imported through adapters or column mapping, and a standardised suite generates shareable artefacts. (B) HTML report from `dfdr_render_report()` for three lists of the basic DIA-NN search: the summary panel lists alerts tagged with their evidential basis rather than a verdict, marks the equal-chance check as not evaluable on these two-pass exports and reports the recomputed decoy ratio under that name. *Alt text:* Flow diagram of the diagFDR workflow from search output through adapters to a standardised target/decoy table, then to scope, calibration and stability diagnostics, with a screenshot of the rendered HTML report.

## 2 Why “1% FDR” is ambiguous

Three sources of ambiguity are worth distinguishing.

First, “FDR = 1%” is usually the value typed into the software; the FDP actually present in the list fluctuates around it, by an amount that depends mainly on how many decoys were accepted (§3.3).

Second, the scope is often undefined. DIA-NN [14] exports a run-wise `Q.Value` next to an experiment-wide `Global.Q.Value`, a `Lib.Q.Value`, and the protein-group equivalents `PG.Q.Valu` and `Global.PG.Q.Value;` Spectronaut distinguishes `EG.Qvalue`, `PG.Qvalue` and `EG.PEP`. Each can be thresholded at 1%, but they control different things on different lists, so the sentence “FDR was set to 1% at the precursor and protein levels” identifies neither. Identifications transferred between runs add to this: the error rate of the assembled matrix need not equal that of any single run [9, 15]. Whenever several exported rows are collapsed into one reported unit, for example precursor rows into a protein group, a rule is needed to decide whether that unit counts as a target or a decoy. This rule is almost never stated (§3.1).

Third, the decoys may not represent incorrect matches. Target–decoy estimation assumes that an incorrect match is as likely to land on a decoy as on a target. Model-derived information can break this symmetry: decoys generated from predicted or measured spectra need not behave like incorrect targets [16]; cross-validated rescoring [17, 18, 19, 20] can leak near-identical matches between folds [21]; and a confidence value computed on a different set of candidates, or from a fitted model, is no longer tied to the decoy count of the reported list (§3.4). When the decoy model is wrong, the counts and the decoy ratio look exactly as expected while the true error is not controlled, which is why independent evidence (A3) is a separate requirement. Neither the number of accepted decoys nor the label rule is currently asked for by the HPP guidelines [22].

To see whether these omissions are occasional or systematic, we scored a convenience sample of 63 recent DIA articles from 34 journals on ten criteria (Note S7, Data S1). The audit records what the main text states, and authors may have done more than they report. None of the 63 articles reported the number of accepted decoys and none stated a label-propagation rule. Eight stated the scope of their threshold and six named the column they thresholded; most gave a count of identifications and a threshold, but only one in five gave the two together (Table S3). Among the 36 studies using DIA-NN, 26 state no scope, so a reader cannot tell whether their counts come from the main report, which DIA-NN filters at `–qvalue` on the run-wise `Q.Value` and on the library q-value `Lib.Q.Value`, not on `Global.Q.Value` (the construction examined in §4.2), or from the matrices, which it filters on both columns. Of the ten DIA-NN studies that state the scope, two filter the main report on `Q.Value` and the library-level `Lib.Q.Value` without the experiment-wide column. On our data this strategy happens to be conservative (Note S7), but neither report gives enough information to establish whether this is also the case for its own data. One study [23] meets eight of the ten criteria, so complete reporting is achievable. The stakes are real: benchmarks on samples of known composition have repeatedly measured DIA error rates above the nominal level [24, 25]; for a nominal 1%, Gu et al. [26] measured protein-level FDRs of 1.81–2.85% with DIA-NN, and Konno et al. [27] 0.76–8.41% across three tools on the same data.

## 3 Scope, calibration and stability

diagFDR works on the table a search engine exports, provided the decoys are included: one row per studied entity (PSM, precursor, peak group, peptide or protein group), a target/decoy label and at least one confidence value. No new search is needed.

Throughout, *α* is the FDR threshold applied to the reported confidence column (usually 1%), *A*(*α*) the set of rows it accepts, and *T*_*α*_ and *D*_*α*_ the numbers of targets and decoys in *A*(*α*) (glossary in Table S7).

### 3.1 Scope: which list, and which labels

Software exports several rows per entity (one per run and precursor; many precursor rows per protein group), so building the list a paper reports involves two rules that are rarely written down. The *aggregation rule* decides which row represents the entity, for example the row with the smallest q-value across runs, or a dedicated experiment-wide column such as `Global.Q.Value.` The *label-propagation rule* decides which target/decoy label the entity inherits when its rows disagree: a protein group with three precursor rows of mixed labels must be counted as one target, one decoy, or (as DIA-NN does for its protein-group q-values) as two competing hypotheses. Four rules are compared in §4.2. A different solution is to pit the target and decoy versions of each protein against each other and keep the better one, so that no label has to be propagated [28, 29].

The scope diagnostic recomputes the decoy ratio *D*_*α*_*/T*_*α*_ on each candidate list. On the list a q-value column was defined for, the ratio is close to *α* by construction, so the check is only informative when it fails: a ratio well above *α* means that the column was thresholded on a list it does not describe, and a ratio well below *α* means that the q-values were computed on another candidate set (mokapot’s exported values in §4.3) or that a further filter was applied, and is read as conservative only once that set or filter is identified. Because control at precursor level does not imply control at protein level, where inference aggregates evidence across peptides and changes the set of hypotheses [30, 31, 32, 33], and because DIA-NN infers proteins only from precursors that already pass a q-value filter (Note S8), the check must be run at the level that is reported.

### 3.2 Calibration: do the confidence values mean what they say

Here “calibration” means agreement with the reference implied by the exported decoys, not agreement with the truth; the latter is question A3.

#### From scores to p-values

Most search engines report a score rather than a p-value, and a score has no fixed meaning: a score of 30 is good or bad depending on the engine, the sample and the database. A p-value fixes the meaning: it is the probability that an incorrect match would score at least as high. If the null distribution is right, the p-values of incorrect matches are uniform between 0 and 1, which a histogram or a quantile–quantile plot can check [34]. In a target–decoy search the decoys provide an empirical null: the p-value of a target match can be defined as the fraction of decoy scores that exceed its score [12]. These *pseudo-p-values* are, up to ties, uniform on the decoys by construction, so the informative check is their distribution on the targets, a mixture of a flat component (incorrect targets, which behave like decoys) and a peak near zero (correct targets). The height of the flat part estimates the proportion *π*_0_ of incorrect identifications among targets [35, 34], and a histogram that is not flat on its right-hand side shows that the decoys describe the incorrect targets poorly in that score range. None of this tests whether decoys are a faithful model of incorrect targets in the first place: pseudo-p-values assume it, and only external evidence can test it (Note S2).

#### PEPs against the decoy reference

The PEP of a target match is the probability that it is incorrect given its score. Under 1:1 competition with equal chance, the number of incorrect targets in a score bin *b* is estimated by the decoys in that bin, *D*_*b*_, so the average target PEP in bin *b* should be close to 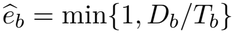. We summarise the disagreement by the size-weighted mean absolute difference, which we call the decoy-reference PEP concordance error (IPE) to make clear that it is an internal check, not a calibration against the truth,

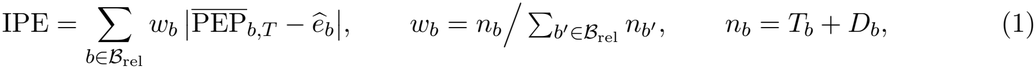

over bins with mean target PEP at most 0.5 and at least 200 entries; across bin counts of 5 to 20 and occupancies of 50 to 500 the IPE of the lists studied below moves by less than 3% (Note S5). An IPE near zero means that the reported PEPs and the exported decoys tell the same story. Two companions do not depend on binning: the sum of PEPs over accepted targets, which should be close to the number of accepted decoys, and the requirement that decoy PEPs be stochastically larger than target PEPs.

#### Equal chance in the low-confidence region

Under 1:1 competition, an incorrect match is a target or a decoy with probability one half, so among matches that are mostly incorrect, those with q-values between 0.2 and 0.5, the decoy fraction should be close to 0.5. We compare the observed fraction with 0.5 using an exact binomial test, a Wilson interval and an effect size. The check only applies to single-pass exports, which excludes DIA-NN’s default two-pass output: in a two-pass pipeline the second search is restricted to a library enriched in first-pass identifications, so the low-confidence region is itself the product of a selection and an excess of decoys there does not indicate a fault (§4.2).

### 3.3 Stability: how much the realised FDP can vary

The decoy ratio stays near *α* by definition, so it says nothing about how far the realised FDP of this particular list may be from *α*. That distance is governed by the number of accepted decoys. When *D*_*α*_ is small, each decoy stands for a large share of the estimated error, and the realised FDP can swing widely from one experiment to the next (Figure 1B). Let *V*_*α*_ be the number of accepted targets that are in fact incorrect, so that FDP_*α*_ = *V*_*α*_*/T*_*α*_.

**Proposition 1** (Approximate variability benchmark for the realised FDP). Assume (i) each incorrect match is a target or a decoy with probability 1*/*2; (ii) these labels are independent across incorrect matches; (iii) every accepted decoy displaced an incorrect match. Ranked by score, the incorrect matches then behave like a sequence of fair coin tosses. If the threshold is idealised as stopping this sequence at the *D*_*α*_-th decoy, the number of incorrect targets accepted before it is approximately negative binomial, 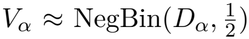, with mean *D*_*α*_ and variance 2*D*_*α*_, and by the law of total variance

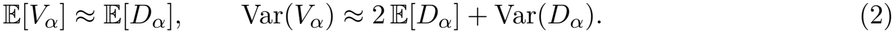

If, in addition, (iv) *D*_*α*_ and *T*_*α*_ vary little between experiments (Var(*D*_*α*_) ≪ E[*D*_*α*_] and CV(*T*_*α*_) ≪ CV(*V*_*α*_)), then

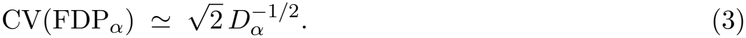

The derivation is in Note S1. It is a heuristic: it describes how *V*_*α*_ varies across experiments with a given decoy support, not its distribution given the observed *D*_*α*_. Unlike the finite-sample prediction bounds of Ebadi et al. [36], it is not a bound but a plug-in value read from a single export: it is neither a confidence interval nor an upper bound for the FDP of the observed list, and the observed *D*_*α*_ is not a fixed stopping index, since the rule that selects it also depends on the target count (Note S1). Simulation confirms it under its assumptions: with independent Gaussian nulls the observed coefficient of variation of the FDP was 1.41 to 1.48 times 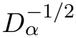 at four thresholds, against the predicted 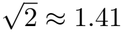 (§4.1). Two departures matter in practice. Shared peptides, match-between-runs and protein grouping correlate labels within blocks of units, violating (ii). We found that simulated block dependence raised the coefficient to 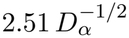. Heavy-tailed null scores let some decoys beat the correct target of their own unit, violating (iii), and raised it by up to 28%. The benchmark is therefore an independence reference, not an exact value; the block dependence simulated here inflated it, and other dependence structures were not examined.

We call 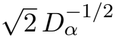 the *FDP-CV benchmark* and *D*_*α*_^*−*1*/*2^ the *decoy-support index*. Because a large *D*_*α*_ can still hide a sparse boundary, `diagFDR` also reports the number of decoys just beyond the cutoff, *D*_*α*,win_ = #{decoys : *q* ∈ (*α*, (1 + *w*)*α*]} with *w* = 0.2, the *threshold elasticity S*_*α*_(*ε*) = *J* (*A*(*α*), *A*((1 + *ε*)*α*)), the Jaccard overlap between the lists at *α* and at a slightly looser threshold, and the change in the decoy ratio when *k* decoys are added or removed.

### 3.4 Better discrimination does not always mean a more stable claim

A rescoring method that separates target and decoy scores better does not necessarily make the FDR claim more reliable; it depends on what is held fixed (Propositions S1 and S2, Note S1). At a fixed nominal level, counting on the reported list ties the decoy count to its size, *D*_*α*_ = *αT*_*α*_ + *O*(1) whenever the step-up threshold is reached inside that list (Proposition S2 states the conditions; q-values taken from another universe or a list truncated before the crossing need not obey it), so admitting more correct identifications also accepts more decoys and the benchmark tightens. At a fixed sensitivity the relationship reverses: raising the threshold empties the decoy tail while barely touching well-separated targets, so E[*D*_*α*_] falls towards zero as the discrimination improves. We call this the “granularity paradox”: at fixed sensitivity, better discrimination empties the decoy tail and the estimate becomes fragile. Which case applies depends on how the lists are compared: a benchmark that matches methods at equal sensitivity is in the second, whereas §4.3, which compares lists at a fixed nominal level, is in the first. Benchmark studies that compare rescoring methods by AUROC should therefore report *D*_*α*_ next to it.

### 3.5 External validation by entrapment

Entrapment adds to the target database a proteome known to be absent from the sample, searched as ordinary targets. Accepted matches that map only to it are known to be false, and they are counted separately from the decoys. With *E*_*α*_ such matches among the *T*_*α*_ accepted targets (*T*_*α*_ counts every accepted non-decoy match, entrapment included), the raw entrapment FDP is 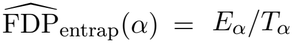. It underestimates the error because an incorrect match can also land on a real target sequence. The correction of Wen et al. [11] accounts for the entrapment-to-target size ratio *r* = *N*_*ET*_ */N*_*T*_,

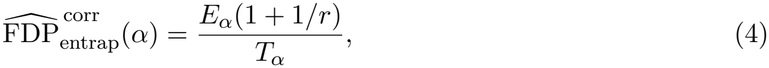

which estimates the FDP of the combined target-plus-entrapment list. Wen et al. show that the raw value is a lower bound on this FDP and that Equation (4) overestimates it on average, so we read them as lower and upper bounds: a corrected value below *α* is consistent with control, a raw value above *α* is evidence against it, and anything in between is unresolved. The “sample” estimate for the original targets only, (*E*_*α*_*/r*)*/*(*T*_*α*_ − *E*_*α*_), is neither bound and typically underestimates [11], so we do not use it for verdicts (Note S4). Homology between the proteomes and differences in detectability can bias all of them, so we use them mainly to compare lists. The size ratio depends on the level at which it is measured and must be reported: the *Arabidopsis thaliana* proteome used below gives *r* = 0.155 at protein level and *r* = 0.432 at tryptic-peptide level after removing the 10,037 peptides (1.04% of the entrapment digest) shared with the target proteomes (Note S4). Raw estimates carry 95% Wilson intervals, which treat accepted units as independent and are therefore too narrow where shared peptides, runs or protein groups cluster the errors. Paired entrapment designs, in which each target is matched with a shuffled entrapment sequence, are an alternative we did not evaluate.

### 3.6 Implementation

`diagFDR` reads DIA-NN, Spectronaut, MaxQuant, mokapot and mzIdentML outputs and any table with the required columns (Figure 2; Table S1 maps every diagnostic to a function). Every number in this article derived from the deposited exports can be recomputed with version 0.2.1 of the package; the simulations use its generative model. A Python port, `diagfdr` (PyPI), implements the same diagnostics for Python-based pipelines and is tested against the R package and against the numbers reported here (Note S8). A single HTML report containing all diagnostics and a summary from a search can be obtained with the `dfdr_render_report` function (https://github.com/Jacky11/diagFDR). Table 2 states what each diagnostic can and cannot detect and what it assumes, while Note S3 gives the corrective action when a check fires.

**Table 1:**
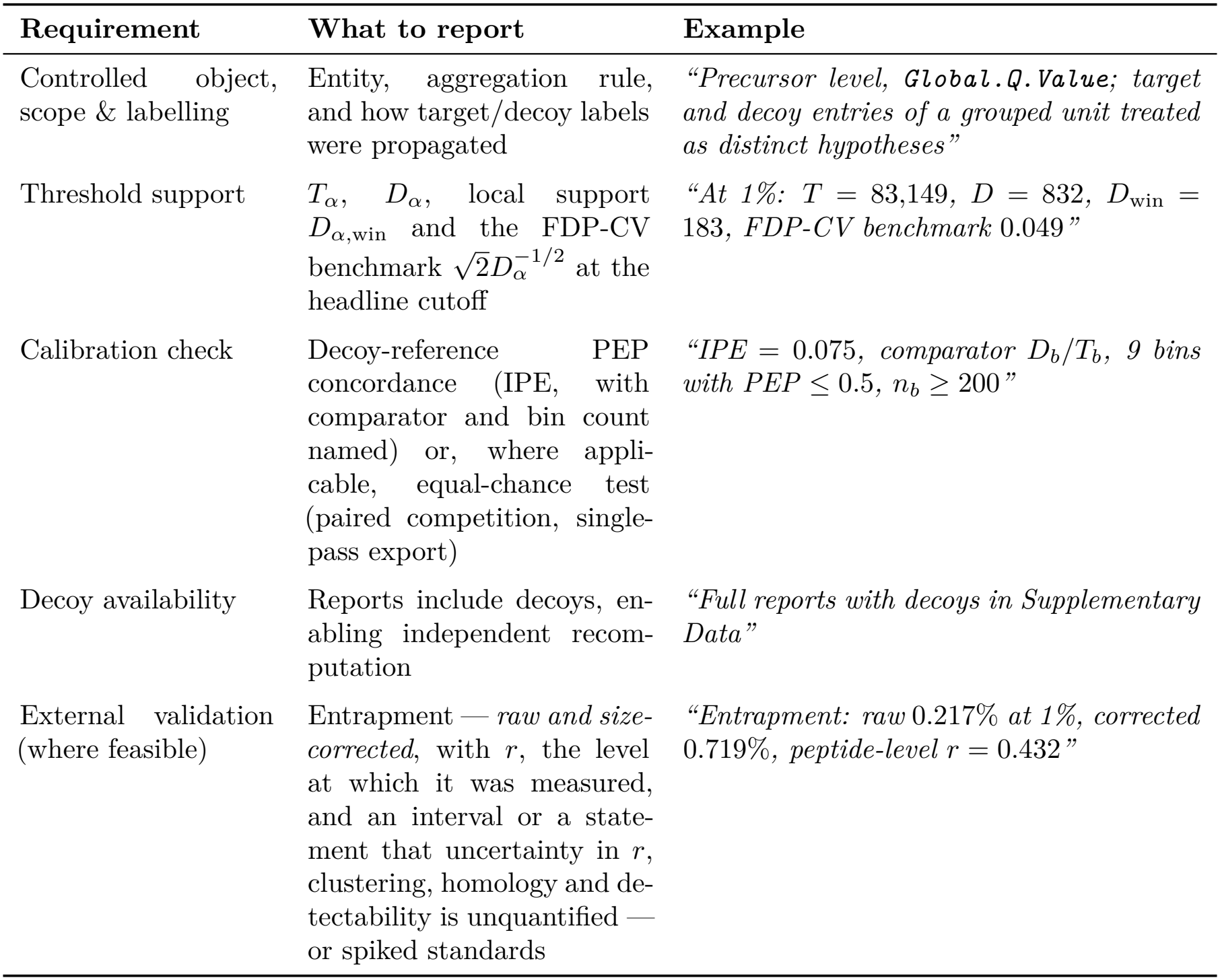
Proposed minimal reporting standard for an auditable “FDR = *α*%” claim. Items 1–4 can be produced from any decoy-inclusive export and are the minimum for every claim; item 5 is the only one bearing on external validity (A3) and is required where an entrapment or known-composition experiment is feasible. The comprehensive checklist is Supplementary Table S2.

| **Requirement** | **What to report** | **Example** |
|---|---|---|
| Controlled object, scope & labelling | Entity, aggregation rule, and how target/decoy labels were propagated | *“Precursor level, **Global.Q.Value**; target and decoy entries of a grouped unit treated as distinct hypotheses”* |
| Threshold support | *T*<sub>α</sub>, *D*<sub>α</sub>, local support *D*<sub>α,win</sub> and the FDP-CV benchmark √2*D*<sub>α</sub><sup>−1/2</sup> at the headline cutoff | *“At 1%: T* = 83,149, *D* = 832, *D*<sub>win</sub> = 183, *FDP-CV benchmark* 0.049*”* |
| Calibration check | Decoy-reference PEP concordance (IPE, with comparator and bin count named) or, where appli-cable, equal-chance test (paired competition, single-pass export) | *“IPE* = 0.075, *comparator D*<sub>*b*</sub>/*T*<sub>*b*</sub>, *9 bins with PEP* ≤ 0.5, *n*<sub>*b*</sub> ≥ 200*”* |
| Decoy availability | Reports include decoys, en-abling independent recom-putation | *“Full reports with decoys in Supplementary Data”* |
| External validation (where feasible) | Entrapment — *raw and size-corrected*, with *r*, the level at which it was measured, and an interval or a state-ment that uncertainty in *r*, clustering, homology and de-tectability is unquantified — or spiked standards | *“Entrapment: raw* 0.217% *at 1%, corrected* 0.719%, *peptide-level r* = 0.432*”* |

**Table 2:**
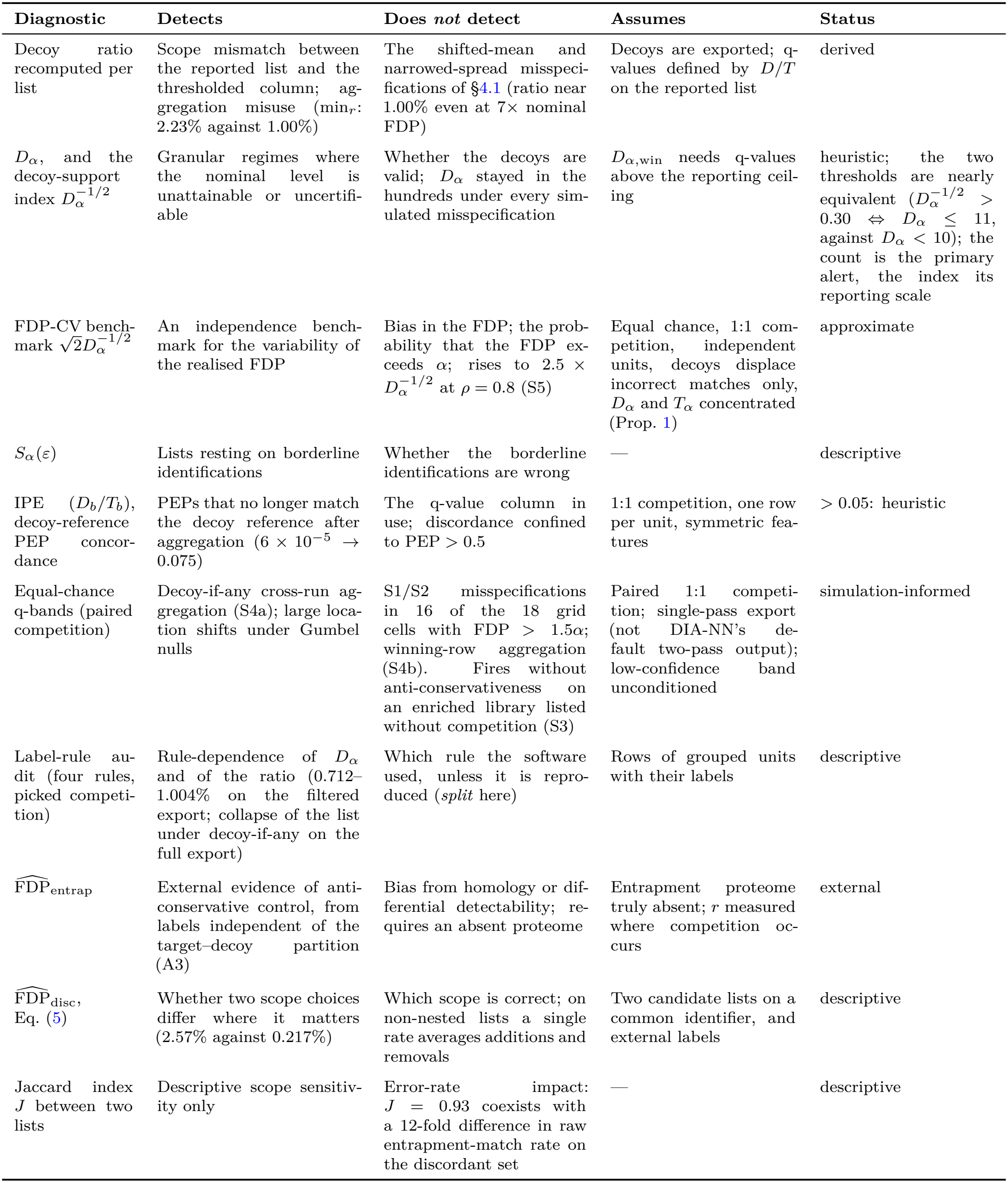
What each diagnostic can and cannot establish. *Status*, the evidential basis of each alert: **derived** check, following from a stated result; **simulation-informed** threshold, calibrated in §4.1; **heuristic** threshold, from the two case studies, to be re-derived locally; **approximate** benchmark; **descriptive**, no threshold; **external**, comparative evidence from independent labels. Alerts motivate inspection, never a pass/fail verdict.

| **Diagnostic** | **Detects** | **Does *not* detect** | **Assumes** | **Status** |
|---|---|---|---|---|
| Decoy ratio recomputed per list | Scope mismatch between the reported list and the thresholded column; ag-gregation misuse (min<sub>r</sub>: 2.23% against 1.00%) | The shifted-mean and narrowed-spread misspeci-fications of §4.1 (ratio near 1.00% even at 7× nominal FDP) | Decoys are exported; q-values defined by D/T on the reported list | derived |
| D<sub>α</sub>, and the decoy-support index D<sub>α</sub><sup>−1/2</sup> | Granular regimes where the nominal level is unattainable or uncertifi-able | Whether the decoys are valid; D<sub>α</sub> stayed in the hundreds under every sim-ulated misspecification | D<sub>α,win</sub> needs q-values above the reporting ceil-ing | heuristic; the two thresholds are nearly equivalent (D<sub>α</sub><sup>−1/2</sup> > 0.30 ⇔ D<sub>α</sub> ≤ 11, against D<sub>α</sub> < 10); the count is the primary alert, the index its reporting scale |
| FDP-CV bench-mark √2D<sub>α</sub><sup>−1/2</sup> | An independence bench-mark for the variability of the realised FDP | Bias in the FDP; the prob-ability that the FDP ex-ceeds α; rises to 2.5 × D<sub>α</sub><sup>−1/2</sup> at ρ = 0.8 (S5) | Equal chance, 1:1 com-petition, independent units, decoys displace incorrect matches only, D<sub>α</sub> and T<sub>α</sub> concentrated (Prop. 1) | approximate |
| S<sub>α</sub>(ε) | Lists resting on borderline identifications | Whether the borderline identifications are wrong | — | descriptive |
| IPE (D<sub>b</sub>/T<sub>b</sub>), decoy-reference PEP concor-dance | PEPs that no longer match the decoy reference after aggregation (6 × 10<sup>−5</sup> → 0.075) | The q-value column in use; discordance confined to PEP > 0.5 | 1:1 competition, one row per unit, symmetric fea-tures | > 0.05: heuristic |
| Equal-chance q-bands (paired competition) | Decoy-if-any cross-run ag-gregation (S4a); large loca-tion shifts under Gumbel nulls | S1/S2 misspecifications in 16 of the 18 grid cells with FDP > 1.5α; winning-row aggregation (S4b). Fires without anti-conservativeness on an enriched library listed without competition (S3) | Paired 1:1 competi-tion; single-pass export (not DIA-NN’s de-fault two-pass output); low-confidence band unconditioned | simulation-informed |
| Label-rule au-dit (four rules, picked competi-tion) | Rule-dependence of D<sub>α</sub> and of the ratio (0.712–1.004% on the filtered export; collapse of the list under decoy-if-any on the full export) | Which rule the software used, unless it is repro-duced (*split* here) | Rows of grouped units with their labels | descriptive |
| $\widehat{\mathrm{FDP}}_{\mathrm{entrap}}$ | External evidence of anti-conservative control, from labels independent of the target–decoy partition (A3) | Bias from homology or dif-ferential detectability; re-quires an absent proteome | Entrapment proteome truly absent; r measured where competition oc-curs | external |
| $\widehat{\mathrm{FDP}}_{\mathrm{disc}}$, Eq. (5) | Whether two scope choices differ where it matters (2.57% against 0.217%) | Which scope is correct; on non-nested lists a single rate averages additions and removals | Two candidate lists on a common identifier, and external labels | descriptive |
| Jaccard index J between two lists | Descriptive scope sensitiv-ity only | Error-rate impact: J = 0.93 coexists with a 12-fold difference in raw entrapment-match rate on the discordant set | — | descriptive |

## 4 Validation and applications

### 4.1 Simulations with known ground truth

We simulated lists of 2 × 10^5^ units in which a target and a decoy score compete. In null units (a fraction *π*_0_ = 0.667) both scores come from the same standardised distribution (Gaussian, Gumbel or Student-*t*_4_), so equal chance holds by construction. In the others, the target score is shifted by *δ* = 4.65, giving accepted decoy counts of the order observed in the DIA-NN data (Note S6). Six scenarios were run with 1000 replicates each: correct specification under the simulated model (S0); decoys too easy to beat (S1, shifted down by 0.35) or under-dispersed (S2, scaled by 0.85); an enriched library listed without competition, as in a second pass (S3); the smallest run-level q-value over six runs, labelled *decoy-if-any* (S4a) or *winning row* (S4b); and block-correlated scores (S5). Four review alerts were defined 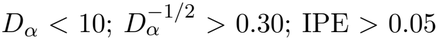 decoy fraction among q-values in [0.2, 0.5] more than 0.15 from 0.5), and a replicate is considered *bad* when its realised FDP exceeds 1.5*α* (Table S4).

First, under correct specification no replicate raised any alert at *α* = 1% or 5% in any null family, and where almost no decoys remained (*α* ≤ 0.002 with heavy-tailed nulls) the support alerts fired in every replicate, refusing a level the list cannot support. The ratio of the observed CV of the FDP to 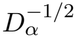 was 1.41 to 1.48 at four thresholds, against the 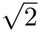 of Proposition 1, and rose to 2.51 under a block correlation of 0.8. The mean FDP stayed at nominal but exceeded 1% in 32% of replicates, because FDR control bounds an expectation, not each experiment. Second, the scope diagnostic detects the aggregation error: under S4b the realised FDP is 5.36% for a nominal 1% and the recomputed decoy ratio reads 5.35%, whereas S4a, the same aggregation with the other label rule, gives 0.18%. Third, the internal diagnostics are blind to a wrong decoy model: under S1 and S2 the mean FDP rose to 2.59% and 2.83% with every replicate bad and no alert, because the decoy ratio stayed at 1% and the equal-chance fraction within tolerance. Across a grid of five shifts and five scalings the FDP reached seven times nominal with no alert in 16 of the 18 cells where it exceeded 1.5*α* (Table S9). With an equal-chance tolerance of 0.05 instead of 0.15, eight cells stay undetected, the largest undetected inflation is 3.9 times, and four cells within 1.5*α* raise an alert (Note S6). Scenario S3 shows the converse: valid decoys and a conservative list, yet an equal-chance fraction of 0.69, the value seen in real two-pass exports. The sign flip of §3.4 is reproduced in Figure S1.

### 4.2 DIA-NN case study

Six DIA runs of a public benchmark were searched with DIA-NN three times: using tryptic (“basic”), semi-tryptic, and basic with the *A. thaliana* proteome appended as entrapment (Materials and methods). Each exported row (one per run and precursor) carries a run-wise `Q.Value` and an experiment-wide `Global.Q.Value`, and protein groups the analogous `PG.Q.Value` and `Global.PG.Q.Value`.

#### Scope

An experiment-wide precursor list at *α* = 1% can be built in three ways: on `Global.Q.Value;` on `Q.Value` alone, keeping each precursor’s best run, which we write min_*r*_(Q.Value); or on both columns at once, which is how DIA-NN filters its precursor matrices. The first and third give the same list on every search (*T* = 83,149, *D* = 832 on the basic search), so the matrices are safe. The main report is filtered on `Q.Value` and on the library q-value `Lib.Q.Value`, not on `Global.Q.Value` (Note S8). In a single-pass search against a predicted library `Lib.Q.Value` is zero, and in an export written at a liberal ceiling, as recommended for the decoys, the library constraint is not binding at 1%. In both cases, keeping the unique precursors of the main report with `Q.Value` ≤ 0.01 in any run gives the second list, whose decoy ratio is 2.23% (basic) and 2.08% (semi-tryptic), because keeping the best run gives an incorrect precursor several chances to pass. In a two-pass search at the default `--qvalue 0.01, Lib.Q.Value` is the first-pass experiment-wide q-value, so the main report carries an implicit global filter and the same count gives a conservative list (*T* = 85,197, *D* = 712, 0.84% on the basic export; Note S8). The inflation grows with the number of runs, from 1% with one run to 2.23% with six (Note S4), so larger cohorts will see more. Protein groups behave the same way (1% against 2.39%) (Table 3, Figure 3A).

**Table 3:**
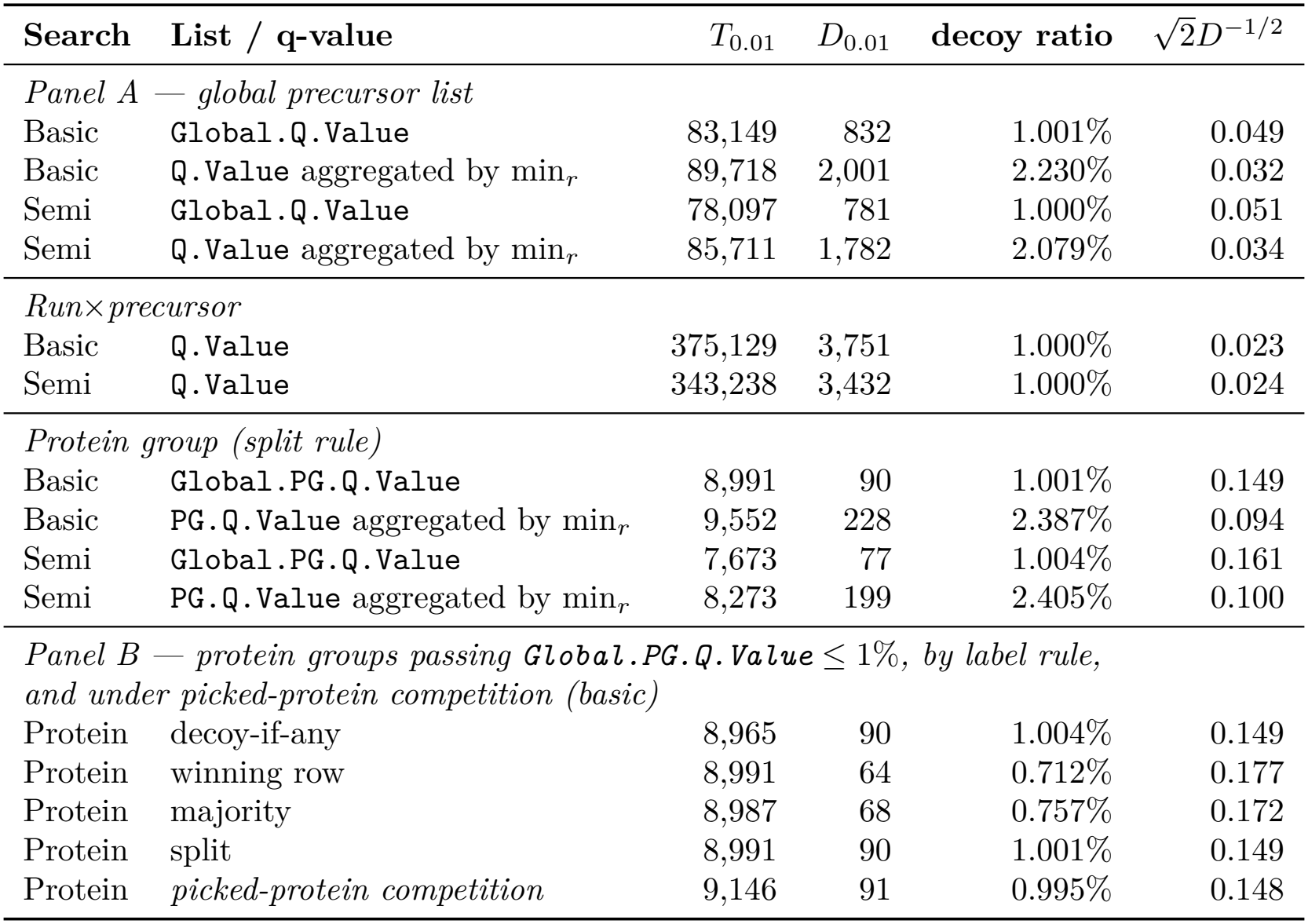
Scope audit of the DIA-NN analysis at *α* = 1%. The decoy ratio *D*_*α*_*/T*_*α*_ recomputes the exported confidence column on the reported list, a consistency check (A2); entrapment (§4.2) supplies the external comparison. For a fixed export the ratio is deterministic and reported without interval; its variability across experiments is that of the realised FDP, for which the last column gives the benchmark of Proposition 1. Precursor counts are after deduplication; the double filter on `Q.Value` and `Global.Q.Value` that DIA-NN applies to its matrices returns exactly the `Global.Q.Value` list and is not shown separately; filtering the main report on `Q.Value` alone, with-out the library or experiment-wide constraint, yields the min_*r*_ rows. Panel B: protein groups with `Global.PG.Q.Value` ≤ 0.01 under the four label-propagation rules of §4.2, which differ only for the 26 groups retaining both a target and a decoy row, and under picked-protein competition, which changes the accepted list. Definitions, a worked example and the values on the unfiltered export are in Note S9.

| **Search** | **List / q-value** | *T*<sub>0.01</sub> | *D*<sub>0.01</sub> | **decoy ratio** | √2*D*<sup>−1/2</sup> |
|---|---|---|---|---|---|
| *Panel A — global precursor list* |  |  |  |  |  |
| Basic | Global.Q.Value | 83,149 | 832 | 1.001% | 0.049 |
| Basic | Q.Value aggregated by min<sub>*r*</sub> | 89,718 | 2,001 | 2.230% | 0.032 |
| Semi | Global.Q.Value | 78,097 | 781 | 1.000% | 0.051 |
| Semi | Q.Value aggregated by min<sub>*r*</sub> | 85,711 | 1,782 | 2.079% | 0.034 |
| *Run×precursor* |  |  |  |  |  |
| Basic | Q.Value | 375,129 | 3,751 | 1.000% | 0.023 |
| Semi | Q.Value | 343,238 | 3,432 | 1.000% | 0.024 |
| *Protein group (split rule)* |  |  |  |  |  |
| Basic | Global.PG.Q.Value | 8,991 | 90 | 1.001% | 0.149 |
| Basic | PG.Q.Value aggregated by min<sub>*r*</sub> | 9,552 | 228 | 2.387% | 0.094 |
| Semi | Global.PG.Q.Value | 7,673 | 77 | 1.004% | 0.161 |
| Semi | PG.Q.Value aggregated by min<sub>*r*</sub> | 8,273 | 199 | 2.405% | 0.100 |
| *Panel B — protein groups passing* ***Global.PG.Q.Value*** ≤ 1%*, by label rule, and under picked-protein competition (basic)* |  |  |  |  |  |
| Protein | decoy-if-any | 8,965 | 90 | 1.004% | 0.149 |
| Protein | winning row | 8,991 | 64 | 0.712% | 0.177 |
| Protein | majority | 8,987 | 68 | 0.757% | 0.172 |
| Protein | split | 8,991 | 90 | 1.001% | 0.149 |
| Protein | *picked-protein competition* | 9,146 | 91 | 0.995% | 0.148 |

**Figure 3:**
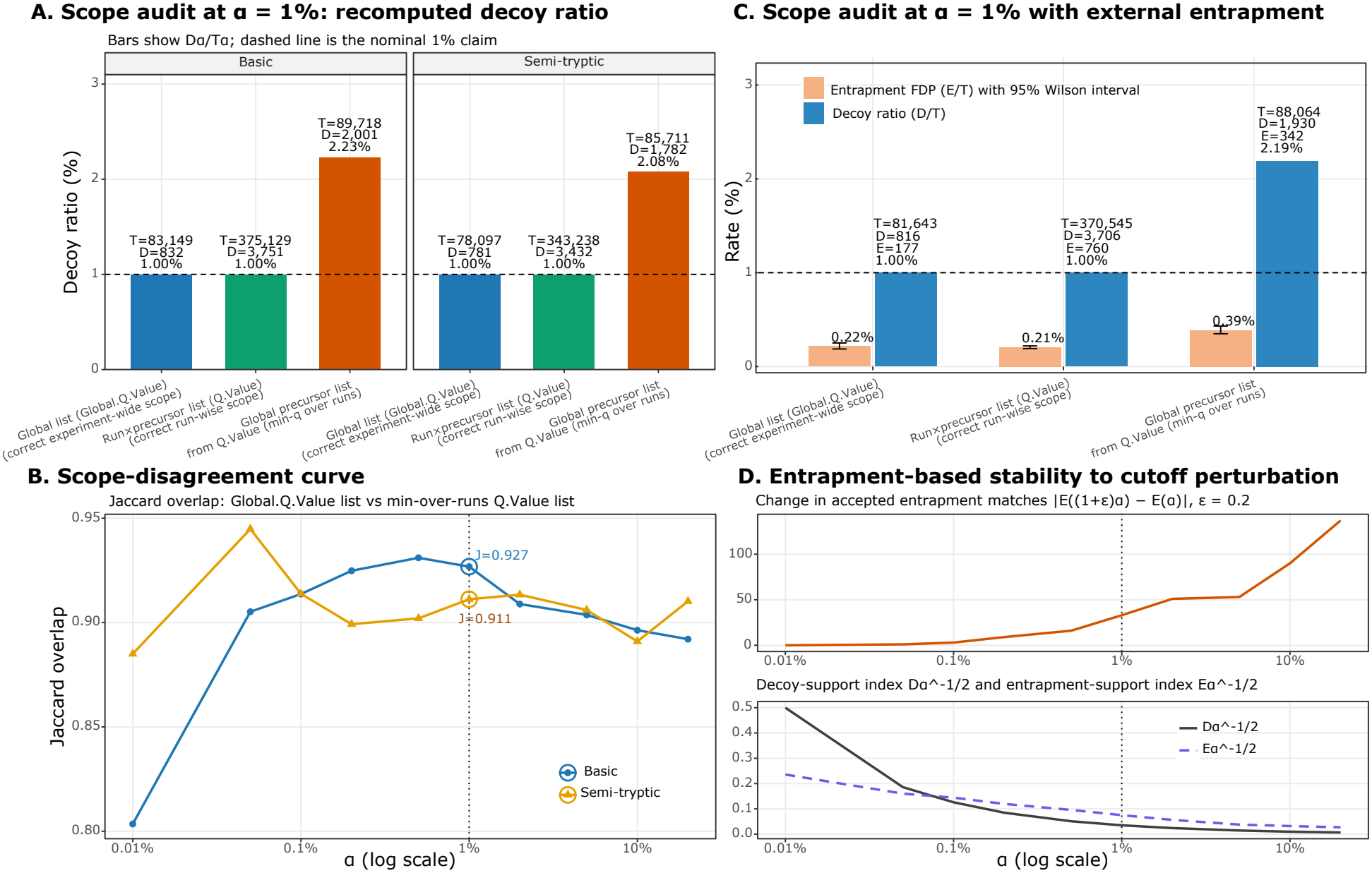
Scope audit and external validation of the DIA-NN analysis. (A) Recomputed decoy ratio on the reported list by q-value construction, for the basic and semi-tryptic searches; the axis is labelled *decoy ratio* because it recomputes the exported column, a consistency check. (B) Jaccard overlap between the `Global.Q.Value` list and the min_*r*_(`Q.Value`) list against *α*; the overlap stays high while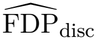 is up to twelve times 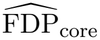 (§4.2, Equation (5)). (C) Recomputed decoy ratio and raw entrapment FDP for the basic search including the *A. thaliana* proteome (counts therefore differ from Table 3, which reports the search without it); entrapment estimates carry Wilson intervals, which are descriptive and ignore the clustering of errors within protein groups (a cluster bootstrap is in Note S4), and decoy ratios do not, the ratio being deterministic for a fixed export. (D) Stability to cutoff perturbation under `Global.Q.Value:` change in the number of accepted entrapment matches when the threshold moves from *α* to 1.2*α* (top), and the decoysupport index 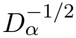next to the analogous entrapment-support index 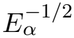(bottom). *Alt text:* Four panels comparing scope choices in the DIA-NN analysis: recomputed decoy ratio by q-value construction, the Jaccard overlap between the global and min-over-runs lists against the threshold, decoy ratio against entrapment-based false discovery proportion, and the stability of both support indices to perturbation of the cutoff.

The entrapment search provides an external check. The global list holds *T* = 81,643 targets and *E* = 177 matches mapping only to *A. thaliana*, a raw entrapment FDP of 0.217% (95% CI [0.187, 0.251]%). The min_*r*_ list holds *T* = 88,064 and *E* = 342, or 0.388% ([0.349, 0.432]%). Corrected for database size (*r* = 0.432) with Equation (4), these become 0.719% and 1.29%: the global list is not contradicted by the entrapment analysis and the min_*r*_ list is not confirmed, although its raw value does not refute it either; both readings assume a valid entrapment proteome and the size correction of Wen et al. (Note S4). Yet the two lists overlap almost entirely (Jaccard index 0.93). The disagreement is confined to their margin, so we compute the entrapment rate separately on the identifications the two lists share and on those on which they disagree,

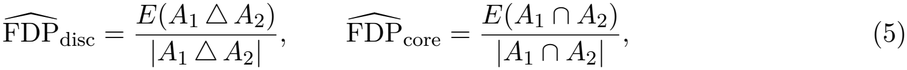

where *A*_1_*ΔA*_2_ is the symmetric difference of the two accepted lists and *E*(·) counts entrapment matches (Note S4). The 6,421 precursors that min_*r*_ adds have 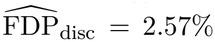 ([2.21, 2.99]%) against 0.217% on the 81,643 shared ones, 11.9 times more (a bootstrap over protein groups, which treats clustered errors as one unit, gives [9.3, 14.9]; Note S4). These added precursors are simply the margin of a looser list: the two columns rank precursors almost identically (Spearman correlation 0.99), and the min_*r*_ list at 1% is exactly the `Global.Q.Value` list at 2.19%. The misuse relabels a 2.2% list as a 1% list, and 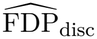 measures what that margin costs. Such a ratio is expected whenever a calibrated list is loosened; the overlap of 0.93 alone would never show how much. The ratio stays between 3.9 and 11.9 from *α* = 10^*−*3^ to 5 × 10^*−*2^ while the Jaccard index stays between 0.91 and 0.94 (Figure 3B).

#### Label propagation

In DIA-NN the target and decoy precursors of a protein share one `Protein.Group` identifier, and 10,261 of the 13,750 groups of the basic search contain both. Collapsed to one entry, a group keeps its smallest q-value but needs a label. We compare four rules (Note S9): *decoy-if-any*; *winning row*, the label of the row that supplied the q-value; *majority*; and *split*, in which the target rows and the decoy rows of a group are two competing hypotheses, as in DIA-NN’s own protein-group q-values. On the groups passing `Global.PG.Q.Value` ≤ 1%, the export a user would normally keep, the rules give decoy ratios of 1.004%, 0.712%, 0.757% and 1.001%, with 90, 64, 68 and 90 accepted decoys for 8,965 to 8,991 targets (Table 3, Panel B). That is a factor of 1.41 in the reported ratio and a 41% change in the decoy count that sets the stability benchmark, all due to 26 groups that keep both a target and a decoy row. On the unfiltered export, decoy-if-any labels 8,092 of the 9,055 accepted groups as decoys, leaving 963 targets, because almost every group carries some low-confidence decoy row (Note S9). The rule to report is the one that reproduces the software’s own construction, here *split*. On the entrapment search the four rules accept nearly the same targets (8,733 to 8,751) with the same raw entrapment FDP (0.65–0.66%) while their decoy ratios range from 0.81% to 1.01%: the list is nearly the same, but the rule changes the reported number. Picked-protein competition [28], which keeps the better of each group’s target and decoy entries and recomputes the q-values, gives the same ratio (*T* = 9,146, *D* = 91, 0.995%) but a longer list: the 75 groups it adds on the entrapment search contain 10 entrapment matches (13%), so the two procedures are not interchangeable. Whatever the rule, the protein-level list carries more error than the precursor-level list: its raw entrapment FDP is 0.663% ([0.513, 0.856]%; *T* = 8751, *E* = 58), three times the precursor-level value, and 2.20% or 4.94% after correction with the peptide-level or the protein-level *r*. Both exceed 1% while the raw value does not, so protein-level control is neither confirmed nor refuted on these data. The range overlaps that of Gu et al. [26], a descriptive comparison only, since instruments, databases and estimands differ (Note S4).

#### Stability and calibration

At *α* = 1% the `Global.Q.Value` list holds 832 decoys (basic; 781 semi-tryptic), a benchmark of 0.049, and 183 decoys in the window just beyond the cutoff. Tighten-ing the threshold erodes this support: at *α* = 10^*−*4^ three decoys remain (benchmark 0.82), too few to support a claim at that level (Figure 4A–C); separately, the raw entrapment FDP there, 5.4 × 10^*−*4^ with an interval excluding 10^*−*4^, is evidence inconsistent with nominal control at that stringency, subject to the same assumption and to only 18 entrapment matches in 14 protein groups; the list is not only unstable there. PEP concordance depends on the unit for which the PEPs were computed: on DIA-NN’s native rows they agree with the decoy reference (IPE = 6 × 10^*−*5^; accepted-target PEPs sum to 3,690 against 3,751 accepted decoys), but collapsed to one row per precursor they fall below it (IPE 0.075; sum 320 against 832; Figure 4D,F). The IPE detects this aggregation, not the scope error: it is identical on the global and min_*r*_ lists. The decoy fraction among q-values in [0.2, 0.5] is 0.66–0.71 on every second-pass list but 0.55 after the first pass (Figure 4E), the pattern simulation S3 reproduces with valid decoys, so the equal-chance check is not evaluable on two-pass exports. Used as intended, on its native rows, `Q.Value` raises no alert.

**Figure 4:**
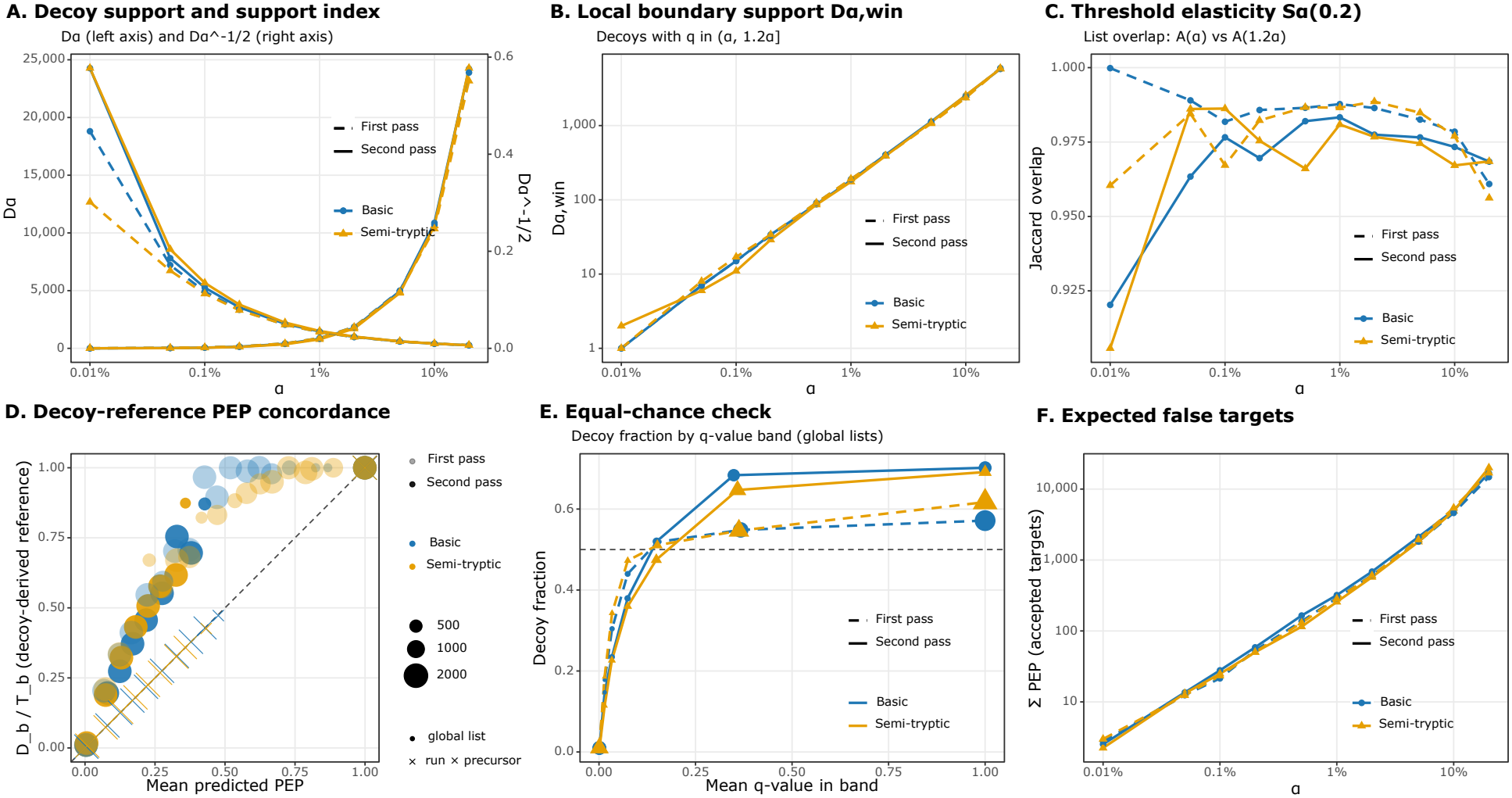
Calibration and stability diagnostics from DIA-NN outputs, comparing first-pass and second-pass (`--reanalyse`) exports under basic and semi-tryptic searches. (A) *D*_*α*_ against *α* (left axis) with the decoy-support index 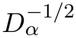(right axis); the FDP-CV benchmark is 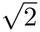 times the index. (B) Local boundary support *D*_*α*,win_ against *α* (*w* = 0.2). (C) Threshold elasticity *S*_*α*_(0.2) against *α*. (D) Decoy-reference PEP concordance: observed 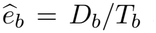 against mean predicted PEP among targets; the run × precursor list lies on the diagonal, the aggregated lists above it. (E) Decoy fraction against mean q-value in band; first-pass exports move towards 0.5 (0.55), second-pass exports do not. (F) PEP among accepted targets against *α. Alt text:* Six line plots of DIA-NN diagnostics against the FDR threshold, comparing first-pass and second-pass exports for basic and semi-tryptic searches: decoy support, local boundary support, threshold elasticity, a PEP concordance curve, decoy fraction by q-value band, and summed PEP among accepted targets.

### 4.3 MS^2^Rescore case study

The second case study illustrates, on a single HeLa run searched with MaxQuant and rescored with MS^2^Rescore and mokapot (Note S8), what rescoring does to the decoy support of a list. Only A1 and A2 can be assessed. After competition, the run has 10,568 spectra with one winning match each. A q-value can be attached to these winners in three ways: by target–decoy counting on the Andromeda score (before rescoring); by taking mokapot’s exported q-values (after rescoring); or by counting again on the mokapot score (after rescoring). At *α* = 1% the three lists hold 5610, 7070 and 7164 targets; at 5 × 10^*−*4^ they hold 2138, 5958 and 6013 (Figure 5D).

**Figure 5:**
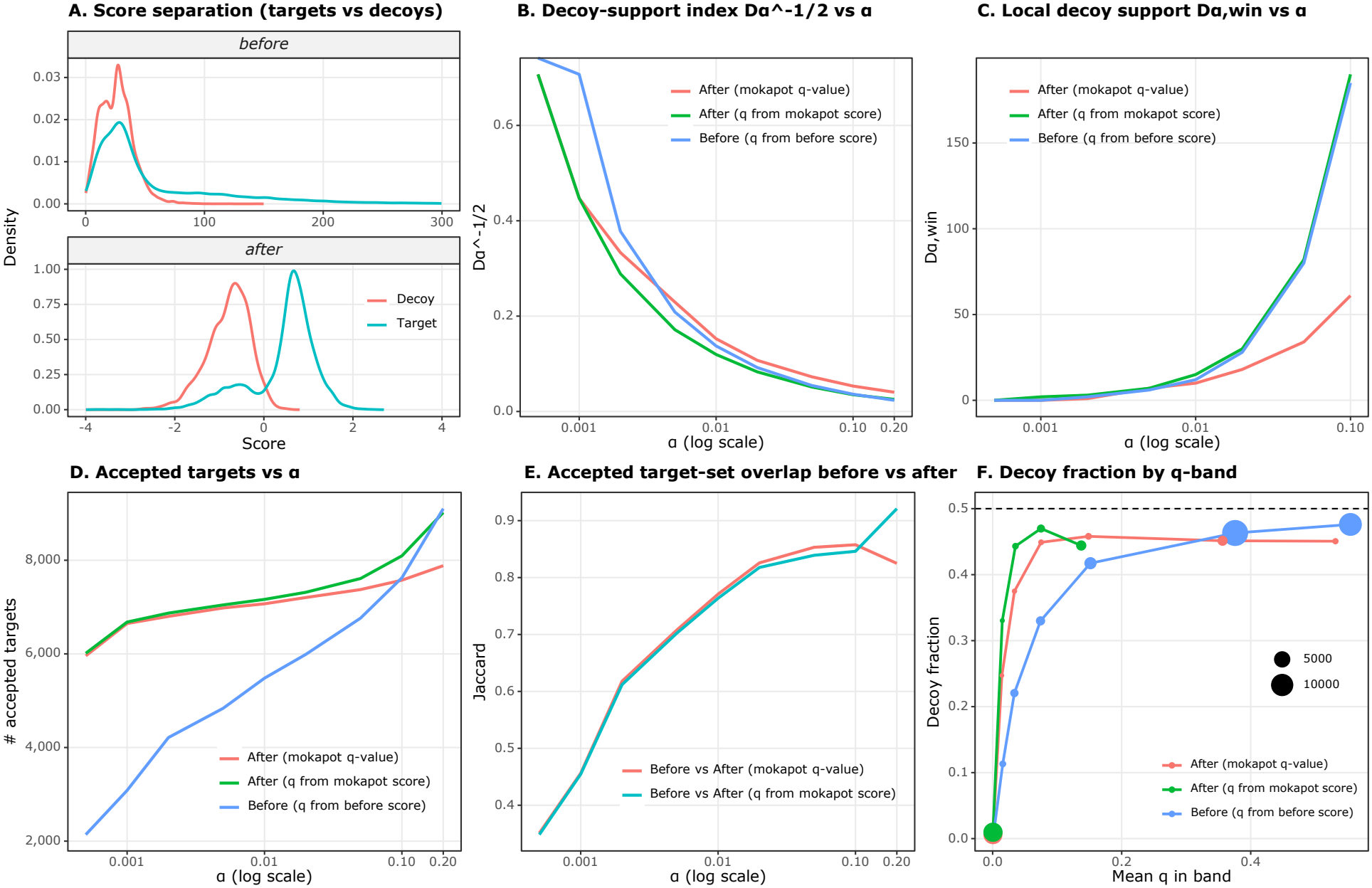
Diagnostics from an MS^2^Rescore analysis under different q-value constructions. This case study has no entrapment arm, so every panel is an internal (A1/A2) diagnostic and none evidences a change in true error control. (A) Score densities for targets and decoys before (Andromeda) and after (mokapot) rescoring. (B) The decoy-support index 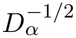 against *α*. (C) *D*_*α*,win_ against *α*. (D) Accepted targets against *α*: before rescoring on all PSMs (5481 at *α* = 1%; 5610 on the spectrum-level winners quoted in the text, Note S8), after rescoring on the spectrum-level winners. (E) Jaccard overlap of the accepted spectra before and after rescoring against *α*; both lists are competed to one winner per run and scan (10,568 spectra each) so that they index the same identifiers. (F) Decoy fraction against mean q-value in band. *Alt text:* Six panels from an MS2Rescore analysis: target and decoy score densities, and curves against the FDR threshold showing the decoy-support index, local boundary support, accepted target counts, the Jaccard overlap between the pre- and post-rescoring accepted lists, and decoy fraction by q-value band.

When q-values are counted on the reported list, more accepted targets bring more accepted decoys, from 55 to 70 at 1%, and the FDP-CV benchmark falls from 0.19 to 0.17, as Proposition S2 predicts at a fixed nominal level. What changes the support much more is which q-value is used. Mokapot’s exported q-values are also target–decoy counts, but over all 31,314 candidate matches, up to ten per spectrum; applied to the 10,568 winners they accept only 43 decoys (0.61%, benchmark 0.22), so the list reported at “1%” holds the decoys of a 0.6% list.

Mokapot’s PEPs agree with the decoy reference (IPE = 0.002) and are slightly conservative: they sum to 59 over the 7070 accepted targets, against 43 accepted decoys. This export is single-pass, so the equal-chance check is evaluable: among the 1624 matches with q-values in [0.2, 0.5] the decoy fraction is 0.451, within the review tolerance although a binomial test rejects exact equality (*p* = 10^*−*4^). The local support just beyond the cutoff is thin for both lists (*D*_0.01,win_ = 10 and 15), so small changes of threshold move the boundary.

At stringent thresholds there is nothing left to measure. Before rescoring, the list at 5 × 10^*−*4^ holds 2138 targets and no decoy (the count uses (*D* + 1)*/T*, so a list reaches 5 × 10^*−*4^ with no accepted decoy once it holds 2,000 targets); after rescoring it holds about 6000 targets and two decoys. The two lists overlap less and less as the threshold tightens, from a Jaccard index of 0.85 at 5% to 0.35 at 5 × 10^*−*4^ (Figure 5E), and no decoy evidence can say which is right. For a reader told only “1% FDR”, the list could be any of three, with 70, 43 or no decoys behind it, and the conservative-looking one has the least support. What is compared here is the universe on which the q-values are counted, not the rescoring algorithm, and the numbers describe this run and this configuration rather than MS^2^Rescore in general.

## 5 Discussion

An FDR claim depends on the scope of the threshold, on the aggregation rule and on the label-propagation rule as much as on the nominal level. On the DIA-NN data, filtering the run-wise Q.Value at 1% gave a 2.2% list whose added precursors carried twelve times the raw entrapment-match rate of the shared ones; the label rule changed the protein-level ratio by a factor of 1.4 on a filtered export, and *decoy-if-any* on the unfiltered export labelled almost every group a decoy; PEPs averaged across runs no longer matched the decoy reference. The diagnostics reveal these classes of error from the export alone, under the conditions examined here. What they are not sufficient to detect is a decoy model that does not represent incorrect matches: in the simulated shift and scaling scenarios, such a model raised the realised FDP to seven times the nominal level without triggering any alert at the review tolerances. Only external evidence can reveal this, in practice an entrapment search analysed with the estimators of §3.5.

The diagnostics have other limits (Table 2): a wrongly scoped list can look more stable than the right one, because it accepts more decoys; a high Jaccard index between two lists can hide a large difference in error on the part where they disagree; and the equal-chance check cannot be read on two-pass exports. The case studies also rest on one DIA dataset and one search engine; the diagnostics read Spectronaut exports too, but their behaviour there, in particular on decoy sets that are not 1:1, was not examined. In return they cost nothing beyond an export that includes the decoys and a liberal q-value ceiling.

The stability benchmark (Proposition 1) could be extended in three directions. It describes how much the realised FDP spreads around its mean, not how likely it is to exceed a given value; prediction bounds [36] and conformal risk control [37, 38] answer that question, the latter without assumptions on the score distribution. It assumes independent units, and conformal methods exchangeable ones, whereas a peptide can map to several proteins and a protein to several peptides. Accounting for this dependence remains open. On real data, it could be checked by generating several decoy sets for one search, for example by shuffling and re-predicting the library [16], and measuring how much the accepted list changes between them.

Learned rescoring adds complexity to an already assumption-dependent procedure. Better separation of targets and decoys does not establish that the decoys remain representative of incorrect target matches, nor that the reported list has adequate support at the chosen threshold. These questions are distinct from discrimination itself, and a nominal FDR level answers neither of them. They also arise when FDR claims are checked automatically: a script, or a language model reviewing a paper, cannot reconstruct the claim without the target–decoy export and the metadata that define its scope.

Existing tools address parts of this problem. `TargetDecoy` [13] provides graphical checks of target–decoy assumptions, while `FDRBench` [11], available through ProteoBench [39], and known-composition samples [26] provide external assessments of FDP. None, however, determines whether the reported “1% FDR” was computed on the reported list, at the stated level, with the relevant label rule and decoy support. `diagFDR` addresses this gap from a decoy-inclusive export that standard search software can produce (Table 4, step 1). It reconstructs the relevant lists, reports target and decoy counts and stability diagnostics, audits grouped-unit labels, and computes entrapment estimates when external validation is available. Its R and Python interfaces and reviewer-facing HTML report turn these checks into a reproducible workflow.

**Table 4:**
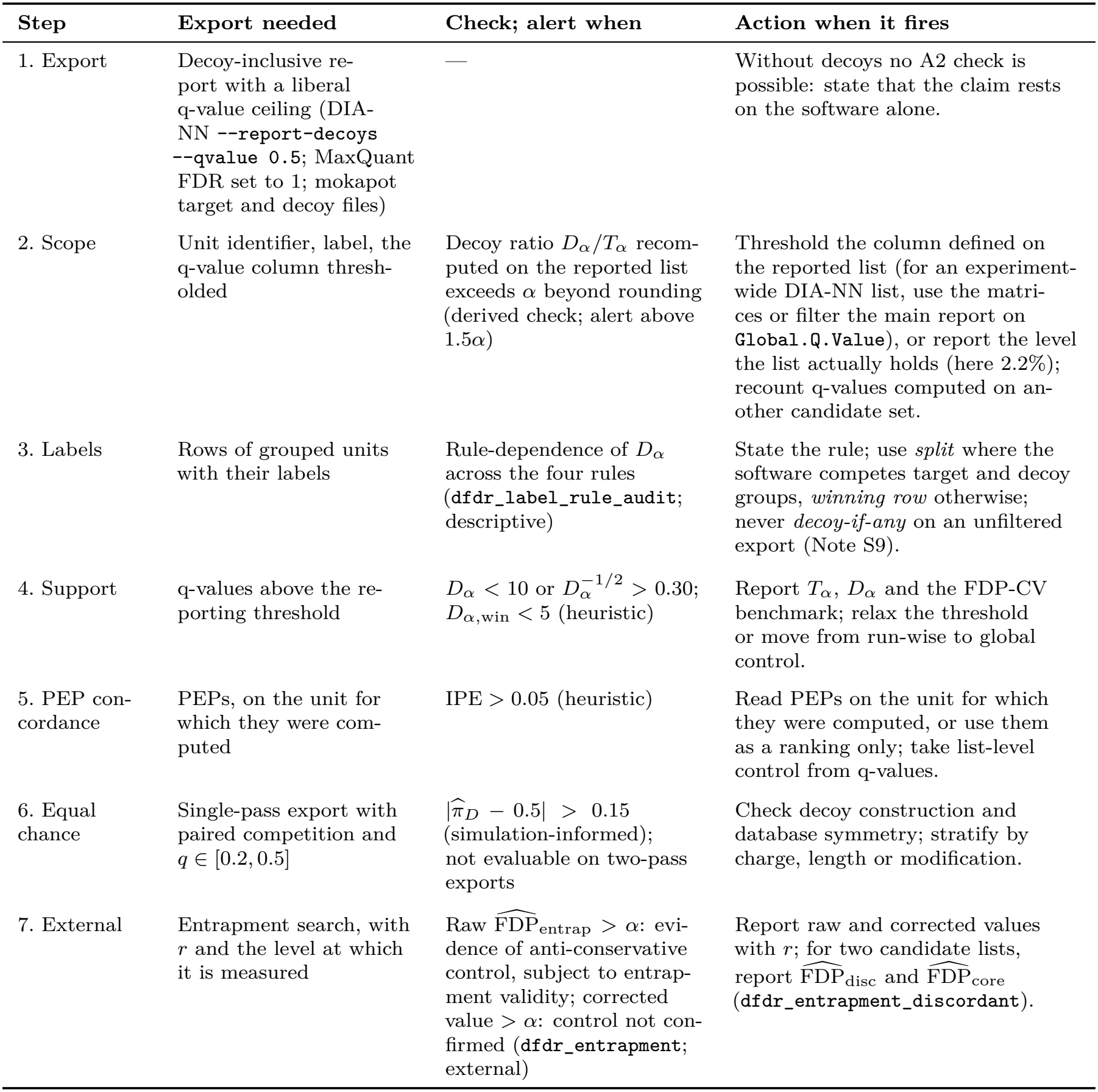
Protocol for an auditable FDR report. Each step lists what the export must contain, the check computed by `diagFDR` (function names in Table S1), the condition that raises a review alert (evidential basis as in Table 2), and the action to take. Alerts motivate inspection; they are not pass/fail verdicts. Corrective strategies are detailed in Note S3.

| Step | Export needed | Check; alert when | Action when it fires |
|---|---|---|---|
| 1. Export | Decoy-inclusive re-port with a liberal q-value ceiling (DIA-NN --report-decoys --qvalue 0.5; MaxQuant FDR set to 1; mokapot target and decoy files) | — | Without decoys no A2 check is possible: state that the claim rests on the software alone. |
| 2. Scope | Unit identifier, label, the q-value column thresh-olded | Decoy ratio D<sub>α</sub>/T<sub>α</sub> recom-puted on the reported list exceeds α beyond rounding (derived check; alert above 1.5α) | Threshold the column defined on the reported list (for an experiment-wide DIA-NN list, use the matri-ces or filter the main report on Global.Q.Value), or report the level the list actually holds (here 2.2%); recount q-values computed on an-other candidate set. |
| 3. Labels | Rows of grouped units with their labels | Rule-dependence of D<sub>α</sub> across the four rules (dfdr_label_rule_audit; descriptive) | State the rule; use *split* where the software competes target and decoy groups, *winning row* otherwise; never *decoy-if-any* on an unfiltered export (Note S9). |
| 4. Support | q-values above the re-porting threshold | D<sub>α</sub> < 10 or D<sub>α</sub><sup>−1/2</sup> > 0.30; D<sub>α,win</sub> < 5 (heuristic) | Report T<sub>α</sub>, D<sub>α</sub> and the FDP-CV benchmark; relax the threshold or move from run-wise to global control. |
| 5. PEP con-cordance | PEPs, on the unit for which they were com-puted | IPE > 0.05 (heuristic) | Read PEPs on the unit for which they were computed, or use them as a ranking only; take list-level control from q-values. |
| 6. Equal chance | Single-pass export with paired competition and *q* ∈ [0.2, 0.5] | \|$\widehat{\pi}_D$ − 0.5\| > 0.15 (simulation-informed); not evaluable on two-pass exports | Check decoy construction and database symmetry; stratify by charge, length or modification. |
| 7. External | Entrapment search, with *r* and the level at which it is measured | Raw $\widehat{\text{FDP}}_{\text{entrap}}$ > α: evi-dence of anti-conservative control, subject to entrap-ment validity; corrected value > α: control not con-firmed (dfdr_entrapment; external) | Report raw and corrected values with *r*; for two candidate lists, report $\widehat{\text{FDP}}_{\text{disc}}$ and $\widehat{\text{FDP}}_{\text{core}}$ (dfdr_entrapment_discordant). |

The five reporting items in Table 1 make the resulting claim auditable. None of the studies in our audit reported them all, although the software used in most of those studies can provide the necessary export. `diagFDR` does not replace external validation, and its internal diagnostics cannot establish that a decoy model is valid. It does, however, make the specification and internal consistency of an FDR claim visible and verifiable. We therefore recommend that decoy-inclusive exports and a `diagFDR` report accompany published FDR claims whenever the underlying software permits them.

## 6 Materials and methods

### Reporting audit

A convenience sample of 63 DIA articles published 2024–2026 was scored from the main text on ten criteria (Note S7). A language model proposed each score with the sentence supporting it; one author checked every score against the article, and a script verified that each quoted sentence occurs verbatim in its source and that the scores are mutually consistent (see the statement on artificial intelligence). The scores and supporting sentences are in Data S1, so any judgement can be re-examined. The two criteria on which the argument rests (accepted decoy count reported; label rule stated) are presence-or-absence checks.

### Simulation

*B* = 1000 replicates of *n* = 2 × 10^5^ units with design values *π*_0_ = 0.667 and *δ* = 4.65 (Note S6; *B* = 500 for the (*π*_0_, *δ*) robustness grid (Table S8) and 200 for the misspecification grid; seed 20260101); q-values by target–decoy counting; the true number of false targets recorded. The generative model is `dfdr_simulate_tdc()` in `diagFDR;` the scenario scripts are in the code repository.

### Datasets

Six-run DIA from the Q Exactive HF-X Orbitrap AIF benchmark [40] (PXD028735), processed with DIA-NN v2.3.1 [14] in tryptic and semi-tryptic two-pass searches (`--report-decoys --qvalue 0.5`) and in a tryptic search with the *A. thaliana* proteome appended; an identification counts towards *E*_*α*_ only when every accession of its protein group belongs to it. Protein-group lists use the *split* rule unless stated (Note S9). The second case study is a MaxQuant search [41] processed with MS^2^Rescore v3.2.0 [42, 43] and mokapot [44] with MS^2^PIP [45] and DeepLC [46] features (Note S8). All numbers derived from these exports were recomputed with `diagFDR` 0.2.1 (Note S8).

## Key Points

- “FDR = 1%” describes a whole procedure: the unit, the scope of the threshold, the confidence column, the aggregation rule and the label-propagation rule must be stated, and none of 63 audited DIA studies stated them all.
- On a DIA-NN dataset, filtering on the run-wise q-value relabelled a 2.2% list as a 1% list, and the identifications it added carried a twelve times higher raw entrapment-match rate behind a Jaccard overlap of 0.93.
- The number of accepted decoys *D*_*α*_ should accompany every threshold: the realised FDP varies between experiments with a coefficient of variation of about 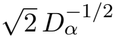 under independence, and more under the block dependence simulated here.
- Internal diagnostics detect scope, aggregation and label errors but are not sufficient against an invalid decoy model, which in simulation raised the realised FDP to seven times nominal undetected; entrapment is a separate requirement.
- `diagFDR` (CRAN; Python port `diagfdr,` PyPI) computes the diagnostics, the label rules and the entrapment estimators and produces a reviewer-facing report; a five-item reporting standard makes its specification and internal consistency auditable.

## Author biographies

**Marie Chion** is a statistician at the MRC Biostatistics Unit, University of Cambridge, working on statistical methods for quantitative proteomics and Bayesian modelling of mass spectrometry data. **Alexandre Godmer** is a clinical microbiologist at Hôpital Saint-Antoine and Sorbonne Université, working on mass spectrometry-based identification and antimicrobial resistance characterisation.

**Thibaut Douché** is a research engineer on the Proteomics Platform at Institut Pasteur, specialising in mass spectrometry acquisition and data processing for quantitative proteomics.

**Mariette Matondo** heads the Proteomics Platform and Mass Spectrometry for Biology Unit at Institut Pasteur, developing quantitative proteomics approaches for infectious disease and host– pathogen biology.

**Quentin Giai Gianetto** is a statistician in the Bioinformatics and Biostatistics HUB and the Proteomics Platform at Institut Pasteur, developing statistical methods and software for mass spectrometry-based proteomics.

## Author contributions

**Marie Chion**: investigation, methodology, formal analysis, writing — review and editing.

**Alexandre Godmer**: investigation, writing — review and editing.

**Thibaut Douché**: investigation, formal analysis.

**Mariette Matondo**: resources, funding acquisition, supervision, writing — review and editing.

**Quentin Giai Gianetto**: conceptualization, methodology, software, formal analysis, supervision, writing — original draft, writing — review and editing.

## Funding

This work was supported by the Agence Nationale de la Recherche [grant number ANR-21-CE35-0007, PureMagRupture project].

## Conflict of interest

The authors declare no competing interests.

## Data availability

diagFDR version 0.2.1 [47], which computes every quantity reported here, is available from CRAN (https://cran.r-project.org/package=diagFDR) and GitHub (https://github.com/Jacky11/diagFDR, release v0.2.1). Raw files, DIA-NN reports (basic, semi-tryptic and entrapment searches, with their first-pass reports and logs) and mokapot outputs are on Zenodo [48] (https://doi.org/10.5281/zenodo.18621806); the DIA benchmark is ProteomeXchange PXD028735. The simulation code, the analysis scripts reproducing every number and table, and the audit pipeline are at https://github.com/Jacky11/diagFDR-paper; the audit scores and evidence are Supplementary Data S1. The Python port diagfdr 0.1.0 is on PyPI (https://pypi.org/project/diagfdr) and at https://github.com/Jacky11/diagfdr-py.

## Use of artificial intelligence

A large language model (Claude Opus 5.5, Anthropic) assisted the reporting audit of Section 2: it read the full text of each article and proposed, for every item, a score together with the sentence supporting it. Every proposed score was checked by one author against the article, and a script verified that each quoted sentence occurs verbatim in its source (Note S7). The scores, the supporting sentences and the script are public (Data S1 and the code repository). An LLM (Claude Opus 5.5, Anthropic) was also used to edit the wording and to check the consistency of the text and code. All content was verified and is the responsibility of the authors.

## Acknowledgements

The authors thank Thomas Burger (CNRS, CEA, Grenoble) for helpful discussions.

## Notes

https://cran.r-project.org/web/packages/diagFDR/index.html

https://doi.org/10.5281/zenodo.18621806

https://github.com/Jacky11/diagFDR/

https://github.com/Jacky11/diagFDR-paper

https://pypi.org/project/diagfdr

https://github.com/Jacky11/diagfdr-py

### Competing Interest Statement

The authors have declared no competing interest.

### Summary of Updates

Revised version submitted to a journal: new title; three-question framework (specification, consistency, independent evidence); Lib.Q.Value mechanism of the DIA-NN two-pass filter clarified; granularity paradox; cluster-bootstrap stability; Python port diagfdr added; diagFDR updated to 0.2.1.

