## Supplemental Data 1 for "diagFDR: Assessing False Discovery Rate Claims in Proteomics with Scope, Calibration and Stability Diagnostics"

**S2 Calibration and stability for p-value-based workflows** 7

**S3 What to do when a diagnostic fires** 7

**S4 Entrapment design** 8

**S10 Supplementary tables and figure** **25**

### S1 Derivation and proofs

This note contains the three analytical results of the paper. Note S1.1 derives Proposition 1 of the main text, the approximation $\mathrm{CV}(\mathrm{FDP}_\alpha) \simeq \sqrt{2}\, D_\alpha^{-1/2}$, and compares it with simulation. Notes S1.3 and S1.4 prove Propositions S1 and S2, which say that better separation of target and decoy scores can either increase or decrease the number of accepted decoys, depending on whether the analyst holds the sensitivity or the nominal level fixed.

#### S1.1 Derivation of Proposition 1

**Setting.** Consider one analysis. Let $T(t)$ and $D(t)$ be the numbers of targets and decoys with score at least $t$. The threshold $t_\alpha$ is the one the step-up rule selects, that is the most liberal $t$ with $D(t)/T(t) \le \alpha$. Write $T_\alpha = T(t_\alpha)$ and $D_\alpha = D(t_\alpha)$, and let $V_\alpha \le T_\alpha$ be the number of accepted targets that are in fact incorrect, so that $\mathrm{FDP}_\alpha = V_\alpha / T_\alpha$.

**Step 1: the sequence of labels.** Keep only the incorrect matches, that is the units whose winning candidate is not a correct identification, order them by decreasing score, and write $L_1, L_2, \dots$ for their target/decoy labels. Assumption (i) says that each $L_j$ is a decoy with probability $\frac{1}{2}$, whatever the score, so the sequence does not depend on the ordering; assumption (ii) says that the $L_j$ are independent. The labels are therefore independent Bernoulli$(\frac{1}{2})$ variables, which is the only probabilistic input of the derivation. Assumption (iii) keeps correct identifications out of the picture: they count in $T_\alpha$ but never in $D_\alpha$ or $V_\alpha$. Under competition this assumption fails whenever a decoy scores higher than the correct target of its own unit, because that decoy is then accepted without standing for any incorrect match. In the simulation of Note S6 at $\alpha = 1\%$ this happens for 2% of accepted decoys with Gaussian nulls, 25% with Gumbel nulls and 18% with Student-$t_4$ nulls (`proposition1_decoy_share.csv`).

**Step 2: a stopping-time picture.** Every accepted decoy is an incorrect match. Think of the accepted set as the incorrect matches that come before the $D_\alpha$-th decoy in the sequence $(L_j)$; the accepted incorrect targets are the target-labelled terms before that point. Counting decoys as successes (probability $\frac{1}{2}$) and incorrect targets as failures gives

$$\Pr(V_\alpha = v) \approx \binom{v + D_\alpha - 1}{v} \left(\tfrac{1}{2}\right)^{D_\alpha + v}, \qquad v = 0, 1, 2, \dots, \tag{S1}$$

the negative binomial distribution NegBin$(D_\alpha, \frac{1}{2})$ in its failures-before-the-$r$-th-success form, with mean $D_\alpha(1-p)/p = D_\alpha$ and variance $D_\alpha(1-p)/p^2 = 2D_\alpha$ at $p = \frac{1}{2}$. Across experiments $D_\alpha$ itself varies; treating it as the stopping index and taking iterated expectations gives $\mathbb{E}[V_\alpha] \approx \mathbb{E}[D_\alpha]$ and $\mathrm{Var}(V_\alpha) \approx 2\,\mathbb{E}[D_\alpha] + \mathrm{Var}(D_\alpha)$, which is Equation (2) of the main text. When $\mathrm{Var}(D_\alpha) \ll \mathbb{E}[D_\alpha]$ (the first half of assumption (iv)) the variance reduces to $2D_\alpha$.

Two details make this an approximation rather than an exact result. First, under the step-up rule the entry just below $t_\alpha$ is a decoy (a target there would lower the ratio), so the accepted set ends just before the $(D_\alpha + 1)$-th decoy. This adds about one to the mean and two to the variance, the same "+1" that appears in finite-sample analyses of competition-based procedures [1], and is negligible when $D_\alpha$ is in the hundreds. Second, $D_\alpha$ is not a fixed index: the rule that selects it also looks at $T(t)$, so the observed $D_\alpha$ carries some information about $V_\alpha$. Among experiments that happen to share the same $D_\alpha$, the distribution of $V_\alpha$ is therefore narrower than NegBin$(D_\alpha, \frac{1}{2})$. The representation describes how $V_\alpha$ varies across experiments with a given decoy support, not its distribution conditional on the observed $D_\alpha$.

**Step 3: from $V_\alpha$ to the FDP.** A first-order expansion of the ratio $V_\alpha/T_\alpha$ gives

$$\mathrm{CV}(\mathrm{FDP}_\alpha)^2 \;\approx\; \mathrm{CV}(V_\alpha)^2 + \mathrm{CV}(T_\alpha)^2 - 2c_{VT}\,\mathrm{CV}(V_\alpha)\mathrm{CV}(T_\alpha), \qquad c_{VT} = \mathrm{corr}(V_\alpha, T_\alpha). \tag{S2}$$

The second half of assumption (iv), $\mathrm{CV}(T_\alpha) \ll \mathrm{CV}(V_\alpha)$, makes the last two terms negligible, so that

$$\mathrm{CV}(\mathrm{FDP}_\alpha) \;\simeq\; \mathrm{CV}(V_\alpha) \approx \frac{\sqrt{2D_\alpha}}{D_\alpha} = \sqrt{2}\,D_\alpha^{-1/2}, \tag{S3}$$

which is Equation (3) of the main text.

**Comparison with simulation.** The benchmark is a plug-in quantity: the variability expected across experiments that have the decoy support of the one observed, with $D_\alpha$ replaced by its observed value. Under correct specification (Note S6, Gaussian nulls, $\alpha = 1\%$), $\mathbb{E}[D_\alpha] = 659$, $\mathrm{Var}(V_\alpha) = 1295$ against $2\,\mathbb{E}[D_\alpha] = 1318$, and the ratio $\kappa = \mathrm{CV}(\mathrm{FDP})/\overline{D_\alpha^{-1/2}}$ is 1.41; from $\alpha = 5 \times 10^{-4}$ to 0.05 it stays between 1.41 and 1.48. A separate run of the same scenario (`06_proposition1_check.R`, 1000 replicates) gives $\mathrm{CV}(\mathrm{FDP}) = 0.054$ against $\sqrt{2}\,\overline{D_\alpha^{-1/2}} = 0.055$ at $\alpha = 1\%$, 0.133 against 0.127 at 0.002, and 0.284 against 0.266 at $5 \times 10^{-4}$ (where $\mathbb{E}[D_\alpha] = 28$). In the same run, among lists with nearly the same $D_\alpha$, the variance of $V_\alpha$ is only 63–86% of $2D_\alpha$ and its mean rises by about 4 to 12 per additional decoy instead of one: this is the conditional narrowing described above. The proposition should therefore not be read as the distribution of $V_\alpha$ given the reported $D_\alpha$; finite-sample upper prediction bounds on the FDP [1] serve that purpose.

**Heavy-tailed nulls: when assumptions (iii) and (iv) fail.** The same script repeats the check with Gumbel and Student-$t_4$ nulls. At $\alpha = 1\%$ both assumptions fail at once. With Gumbel nulls, 25% of accepted decoys come from units that carry a correct target, so $\mathbb{E}[V_\alpha] = 279$ is below $\mathbb{E}[D_\alpha] = 369$, and $D_\alpha$ varies much more than with Gaussian nulls: $\mathrm{Var}(V_\alpha) = 3531$, close to $2\,\mathbb{E}[D_\alpha] + \mathrm{Var}(D_\alpha) = 3633$ but far from $2\,\mathbb{E}[D_\alpha] = 738$. With Student-$t_4$ nulls (18% of decoys from units with a correct target) the three values are 1369, 1264 and 1210. The ratio $\kappa$ is 1.73 and 1.62 in this run (1.81 and 1.67 in the reference run of Note S6, `reference_summary.csv`), against 1.41 with Gaussian nulls; at $\alpha = 5\%$ it returns to 1.42–1.46 in all three families. When correct and incorrect scores overlap in their tails, the benchmark is therefore a lower reference.

**What assumption (iv) asks for in numbers.** Write $T_\alpha = C_\alpha + V_\alpha$ with $C_\alpha$ the accepted correct identifications. Since $V_\alpha/T_\alpha \approx \alpha \ll 1$, $T_\alpha$ is driven by $C_\alpha$, and since $D_\alpha \simeq \alpha T_\alpha$ at the selected threshold, the condition $\mathrm{CV}(T_\alpha) \ll \sqrt{2}\,D_\alpha^{-1/2}$ becomes

$$\mathrm{sd}(T_\alpha) \;\ll\; \frac{\sqrt{2D_\alpha}}{\alpha}. \tag{S4}$$

At $\alpha = 1\%$ with $D_\alpha = 832$, the value of the DIA-NN global list (main-text Section 4.2), this allows $\mathrm{sd}(T_\alpha)$ up to about $4 \times 10^3$ for $T_\alpha \approx 8 \times 10^4$, that is any $\mathrm{CV}(T_\alpha)$ well below 4.9%. Well-separated score distributions give exactly that: the accepted correct identifications sit far above the boundary and barely move between replicates. A small $D_\alpha$ relaxes the condition rather than breaking it, because $\mathrm{CV}(V_\alpha)$ grows as $D_\alpha$ falls; a granular list is uninformative because the benchmark itself is large, not because (iv) fails.

**Dependence, and what the benchmark is not.** Assumption (ii) is the one that fails in real data: shared peptides, match-between-runs and protein grouping all correlate labels within blocks of units. Scenario S5 of Note S6 raises $\kappa$ from $\sqrt{2} = 1.414$ to 1.47, 1.80 and 2.51 at within-block correlations $\rho = 0.2$, 0.5 and 0.8, so Equation (3) of the main text is an independence benchmark; we have not shown that every dependence structure inflates it. The benchmark is also a coefficient of variation, not a tail probability: $\Pr(\mathrm{FDP}_\alpha > \alpha)$ depends on the shape of the distribution of $V_\alpha$, on the realised target count and on the dependence structure. Under correct specification, the simulated FDP exceeded nominal in 32% of replicates at $\alpha = 1\%$ while its mean stayed at 0.98%.

##### S1.2 Discrimination and decoy support: the model

Take a location model in which a single parameter governs discrimination. Decoy scores follow $S_D \sim \mathcal{N}(0,1)$ and correct target scores $S_T^+ \sim \mathcal{N}(\delta, 1)$ with $\delta > 0$; under equal chance, incorrect target matches are distributed like $S_D$. Write $\phi$ and $\Phi$ for the standard normal density and distribution function and $\bar{\Phi} = 1 - \Phi$. The area under the ROC curve is

$$\mathrm{AUROC}(\delta) = \Phi\left(\delta/\sqrt{2}\right), \tag{S5}$$

strictly increasing in $\delta$.

Fix a target sensitivity $S^* = \Pr(S_T^+ \geq t_\alpha)$. Inverting gives $t_\alpha = \delta + z^*$ with $z^* = \Phi^{-1}(1 - S^*)$, so with $n_D$ independent decoy scores

$$\mathbb{E}[D_\alpha] = n_D\, \bar{\Phi}(\delta + z^*). \tag{S6}$$

##### S1.3 Proposition S1 (fixed sensitivity)

*Proposition S1.* In the Gaussian location model, with the sensitivity $S^*$ held fixed, AUROC increases strictly with $\delta$ while $\mathbb{E}[D_\alpha]$ decreases strictly; the decrease accelerates in proportional terms, each further gain of $\varepsilon$ in $\delta$ removing a larger fraction of the remaining expected decoys than the last (the absolute decrement $n_D \phi(\delta + z^*)$ itself shrinks); and $\mathbb{E}[D_\alpha] \to 0$ as $\delta \to \infty$.

*Proof.* Differentiating,

$$\frac{\partial\, \mathrm{AUROC}}{\partial \delta} = \frac{1}{\sqrt{2}} \phi\left(\frac{\delta}{\sqrt{2}}\right) > 0, \qquad \frac{\partial\, \mathbb{E}[D_\alpha]}{\partial \delta} = -n_D\, \phi(\delta + z^*) < 0, \tag{S7}$$

so the two quantities move in opposite directions for every $\delta > 0$: the curve $\delta \mapsto (\mathrm{AUROC}, \mathbb{E}[D_\alpha])$ is a strictly decreasing frontier. By the Mills ratio bound $\bar{\Phi}(x) \leq \phi(x)/x$, with $\bar{\Phi}(x) \sim \phi(x)/x$ as $x \to \infty$, Equation (S6) decays faster than exponentially in $\delta$, $\mathbb{E}[D_\alpha] \propto \exp\{-(\delta + z^*)^2/2\}$ (a Gaussian property, see Note S1.5), while the marginal AUROC gain $\phi(\delta/\sqrt{2})/\sqrt{2}$ falls to zero. As $\delta \to \infty$, AUROC $\to 1$ while $\mathbb{E}[D_\alpha]^{-1/2} \to \infty$, and with it the FDP-CV benchmark. This statement concerns the expected support; in a given sample with $D_\alpha = 0$ the decoy-based claim is simply unsupported.

The acceleration is the second clause. For a fixed increment $\varepsilon > 0$ the ratio $r(\delta) = \bar{\Phi}(\delta + z^* + \varepsilon)/\bar{\Phi}(\delta + z^*)$ satisfies $\mathrm{d}\log r/\mathrm{d}\delta = h(\delta + z^*) - h(\delta + z^* + \varepsilon) < 0$, because the normal hazard rate $h(x) = \phi(x)/\bar{\Phi}(x)$ is strictly increasing: $h'(x) = \phi(x)\{h(x) - x\}/\bar{\Phi}(x) > 0$ since the Mills bound gives $h(x) > x$. Each further gain in discrimination therefore removes a larger share of the remaining support than the last while returning less AUROC; the statement is about the survival ratio, not about the absolute decrement.

##### S1.4 Proposition S2 (fixed nominal level)

*Proposition S2.* If q-values are obtained by target–decoy counting on the reported list, $\alpha$ is held fixed, and the step-up threshold at level $\alpha$ is attained inside that list (the candidate set contains the crossing of the decoy ratio with $\alpha$, so the list is neither accepted whole nor truncated before the crossing by an export ceiling or a sparse q-value grid, and the accepted boundary is followed by the decoy that first violates the threshold), then $D_\alpha = \alpha T_\alpha + O(1)$, that is $D_\alpha/T_\alpha = \alpha + O(T_\alpha^{-1})$. The relation need not hold when the q-values were assigned on another universe than the reported list, when the list was filtered or aggregated after selection, or when no crossing occurs. In the independent location-shift model on a fixed finite set of units, where only the correct-target scores move with $\delta$, null target and decoy scores are unchanged, assumption (iii) of Proposition 1 holds (accepted decoys displace incorrect matches only) and the q-values are recomputed on the same reported list, $D_\alpha$ and $T_\alpha$ are non-decreasing in $\delta$ for every realisation, hence $\mathbb{E}[D_\alpha]$ is non-decreasing in $\delta$, up to the ceiling $\alpha m_1$.

*Proof.* In practice analysts fix $\alpha$, not the sensitivity. If the q-values come from target–decoy counting on the reported list, the operating threshold is the most liberal score $\tau_\alpha = \min\{t : D(t)/T(t) \le \alpha\}$ (with $D(t)$ and $T(t)$ the decoys and targets scoring at least $t$), so at that threshold

$$D_\alpha = \alpha\, T_\alpha + O(1), \tag{S8}$$

the $O(1)$ coming only from the discreteness of the step-up rule. Decoy support is therefore set by the number of accepted targets. The monotonicity in $\delta$ is pointwise under a coupling: generate every score once and let the location shift act on the correct targets only, so that for a given realisation of the noise a larger $\delta$ raises each correct-target score and leaves every null score where it was. Under assumption (iii) of Proposition 1 (accepted decoys displace incorrect matches only), $D(t)$ then does not depend on $\delta$ while $T(t)$ is non-decreasing in $\delta$ for every $t$, so $D(t)/T(t)$ is non-increasing in $\delta$, the set $\{t : D(t)/T(t) \le \alpha\}$ can only grow, $\tau_\alpha$ is non-increasing and $D_\alpha = D(\tau_\alpha)$ and $T_\alpha$ are non-decreasing in $\delta$ for that realisation. Taking expectations gives the statement; the result is exact for finite lists, not asymptotic. Without (iii) a decoy can beat the correct target of its own unit; raising $\delta$ turns some of these decoy wins into target wins, so $D(t)$ can decrease with $\delta$; the coupling argument then no longer applies and the monotonicity has to be assessed under the specific competition mechanism or empirically (such decoys are 18% of the accepted decoys with Student-$t_4$ nulls, Note S1.1, and under 1% with Gaussian nulls; Figure S1 shows the expected behaviour in both cases). The decoy-support index $(\alpha T_\alpha)^{-1/2}$ is accordingly non-increasing, up to a ceiling: with $m_1$ the number of units carrying a correct match, $T_\alpha \uparrow m_1$ and $\mathbb{E}[D_\alpha] \uparrow \alpha m_1$. The statement is for the independent location-shift model with decoys left unchanged; it does not cover rescoring that also moves the decoy scores.

Two situations free $D_\alpha$ from $\alpha T_\alpha$ even at fixed $\alpha$: when the confidence value is not a decoy ratio computed on the reported list (because it comes from a model, or was counted on another set of candidates, as when mokapot q-values computed over all candidate PSMs are applied to spectrum-level winners; main-text Section 4.3), and when $\alpha$ is so stringent that $T_\alpha$ is far from $m_1$ and falling.

#### S1.5 How general are these results

The three clauses of the two propositions need the Gaussian assumption to different degrees.

*The sign in Proposition S1 is general.* Let $F_0$ be any continuous distribution with density $f_0 > 0$ on the relevant range, with decoy scores $S_D \sim F_0$ and correct targets $S_T^+ \sim F_0(\cdot - \delta)$. Fixing the sensitivity $S^*$ fixes $t_\alpha - \delta = z^* = F_0^{-1}(1 - S^*)$ whatever $F_0$ is, so $\mathbb{E}[D_\alpha] = n_D\{1 - F_0(\delta + z^*)\}$ and

$$\frac{\partial\, \mathbb{E}[D_\alpha]}{\partial \delta} = -\, n_D\, f_0(\delta + z^*) < 0. \tag{S9}$$

Discrimination increases with $\delta$ in any location family, because $S_T^+$ then stochastically dominates $S_D$ and AUROC $= \Pr(S_T^+ > S_D)$ grows. The frontier is therefore strictly decreasing for every continuous location family, with no assumption on unimodality or tails.

*Proposition S2 is general only in part.* The relation $D_\alpha = \alpha T_\alpha + O(1)$ needs no distributional assumption: it is a property of the step-up rule, under the conditions listed in the statement (the crossing lies inside the reported list and the q-values were counted on it). It fails, by construction, for a list whose q-values were computed on another universe, for a list cut by an export ceiling before the crossing, and for a list accepted whole, which is why the main text distinguishes the decoy ratio recomputed on the reported list from a q-value column inherited from elsewhere. The monotonicity does need one. In the independent location-shift model, increasing $\delta$ raises the number of accepted correct targets at a fixed target/decoy ratio, so expected decoy support is non-decreasing; extending this to an arbitrary workflow would require conditions on the score ordering, the dependence structure and the threshold rule. The Gaussian assumption only buys the closed form.

*The rate and the acceleration are not general.* The decay $\exp\{-(\delta + z^*)^2/2\}$ is specific to the Gaussian tail, and the acceleration clause requires the hazard rate $h = f_0/(1 - F_0)$ to be increasing, which holds for the Gaussian and the Gumbel but not in the tail of a Student-$t$. Under heavy tails the decay becomes polynomial and the effect is weaker but keeps its sign. The model also holds the decoy distribution fixed while the targets move, whereas rescoring re-scores the decoys too; neither proposition addresses this.

The practical consequence is simple. As scoring improves, the $\alpha$ in use is pushed further into the tail, where few or no decoys remain to support the estimate, and decoy-based FDR becomes numerically fragile even though targets and decoys are well separated. Stringent claims should therefore be read together with $T_\alpha$, $D_\alpha$ and a support indicator, and, when the claim rests on a small sub-list, with the smallest $\alpha^*$ at which $D_{\alpha^*} \geq K$ (say $K = 50$).

### S2 Calibration and stability for p-value-based workflows

When the confidence value is a p-value, calibration has a precise meaning under the Benjamini–Hochberg procedure [2, 3]: for true nulls the p-values must be super-uniform, $\Pr(p_i \leq u \mid H_{0,i}) \leq u$. This is checked with histograms, quantile–quantile plots or calibration plots; `dfdr_plot_p_calibration()` draws both.

Most proteomics workflows output scores, which must first be turned into p-values. Couté *et al.* [4] proposed a transformation that makes BH-style procedures and their diagnostics available; `score_to_pvalue()` returns p-values when the assumptions are reasonably met and *pseudo*-p-values otherwise (the option `decoy_ecdf` uses the empirical distribution of the decoy scores as the null). When the p-values are approximately calibrated, $\pi_0$ can be estimated [5, 3] and read as the proportion of incorrect identifications, the null hypothesis being "this identification is incorrect"; the estimated FDR at threshold $t$ is then $\widehat{\mathrm{FDR}}(t) = \hat{\pi}_0 mt/R(t)$. A useful consistency check is to estimate $\hat{\pi}_0$ separately on the decoys, where it should be close to 100% because every decoy is incorrect, and on the targets.

Stability is less direct under BH than under target–decoy competition, because the result depends on the effective number of tests $m$ and on any pre-filtering. Two practical checks are implemented in `dfdr_bh_diagnostics()` and `dfdr_bh_elasticity()`: rerun BH after reasonable changes of the pre-filtering and report the change in the number of discoveries and the Jaccard overlap of the discovery sets; and count the p-values in a small neighbourhood of the BH cutoff, since a sparse neighbourhood means a list that is sensitive to small perturbations.

### S3 What to do when a diagnostic fires

**PEPs disagree with the decoy reference.** In order of preference: remove the selection bias, since the disagreement usually comes from a pre-filtering that enriches the export in high-confidence identifications, and rerunning with relaxed filters is the only remedy that addresses the cause; apply an empirical correction, which is appropriate only when the bias is systematic across the confidence range and the ranking remains meaningful; or use the PEPs only to rank identifications, without reading their values as probabilities, and take list-level control from the q-values.

**Equal chance fails under paired competition.** Check that the target and decoy databases are comparable in size and complexity. For predicted libraries, shuffle-and-predict decoys have been reported to give better target/decoy symmetry than template-based perturbation [6]. When the decoy fraction $\widehat{\pi}_D$ departs from 0.5 by a stable, measurable factor, a correction may be possible under a validated mislabelling model, but no correction repairs decoys that do not represent incorrect targets. Add external validation by spiking sequences known to be absent. Stratify the diagnostics by charge state, modification status, peptide length or intensity to locate where equal chance fails. When the failure is of the kind characterised by Pomogaev *et al.* [7], their MS1 precursor-envelope filter is one available remedy.

**Scope mismatch.** When the decoy ratio recomputed on the reported list departs from $\alpha$, the column that was thresholded was defined on another list. Either threshold the column defined on the list being reported (for an experiment-wide DIA-NN precursor list, `Global.Q.Value` rather than the minimum of `Q.Value` across runs), or keep the list and report the level it actually holds, which is the threshold of the matching column that returns the same list (2.2% instead of 1% in main-text Section 4.2). When the q-values were computed on a larger set of candidates than the one reported, as for mokapot's candidate-level q-values applied to spectrum-level winners, recount them on the reported list (`mokapot_competed_universe(q_mode = "recount")`).

**Sparse decoy support.** Consider relaxing the threshold, or switching from run-wise to global control in multi-run experiments. Sparse support should not be passed on silently to higher levels: if control is applied at precursor level with little support, protein-level inference needs explicit modelling. Report the situation, for example "at FDR = 0.01%, $D_\alpha = 3$, decoy-support index 0.58 (FDP-CV benchmark 0.82); the accepted list should be regarded as unstable".

#### S4 Entrapment design

An entrapment proteome must be known to be absent from the sample, large enough to give a measurable $E_\alpha$ at the operating cutoff, and distant enough phylogenetically to avoid ambiguous mapping through homology. Entrapment matches must not be added to $D_\alpha$, because that would change the estimator. We use a conservative *pure entrapment* rule: an accepted identification counts towards $E_\alpha$ only if every accession of its protein group belongs to the entrapment proteome. Counting instead every identification with at least one entrapment accession gives the same $E_\alpha$ on every list analysed here (177, 342 and 760 at $\alpha = 1\%$ for the global, $\min_r$ and run×precursor lists).

Because an entrapment estimate from a foreign proteome can be biased in either direction [8], we read it comparatively. $\widehat{\mathrm{FDP}}_{\mathrm{entrap}}(\alpha) \approx \alpha$ is consistent with valid control but does not prove it, since an optimistic decoy estimate and a pessimistic entrapment estimate can cancel; whereas $\widehat{\mathrm{FDP}}_{\mathrm{entrap}}(\alpha) \gg \alpha$ is strong evidence of anti-conservative control, provided the entrapment is valid. Wen *et al.* also describe a *paired* design, in which each target peptide is paired with a shuffled entrapment version so that the two compete and the size ratio is one by construction; it requires a modified library or database and was not evaluated here, which the main text lists as a limitation. When $E_\alpha$ is small, Wilson intervals show whether sampling variability dominates. These intervals treat accepted units as independent and ignore the clustering induced by shared peptides, runs or protein groups, so they are too narrow wherever such clustering exists; this applies to every interval in this Note, including the discordant/core comparison and the protein-level estimate.

##### S4.1 Composition of the database

The entrapment proteome is the reviewed *Arabidopsis thaliana* proteome downloaded on 6 February 2026: 16,418 accessions appended to the ProteoBench mixed-species target database (106,054 accessions including contaminants). *In silico* tryptic digestion under the cleavage rule of the search (after K and R, one missed cleavage, Note S8; peptide length 6–30, `dfdr_entrapment_size_ratio()`) gives 2,206,114 target and 962,791 entrapment peptides, of which 10,037 are shared. The entrapment-to-target size ratio is therefore $r = 0.155$ at protein level, 0.436 at peptide level including the shared peptides, and 0.432 excluding them. With the length range of the DIA-NN search (8–25) the peptide-level ratio is 0.437; the choice changes the corrected estimates by about one percent relative and none of the conclusions.

One practical point deserves a record because it is exactly the kind of silent failure this paper is about. `Protein.Group` in the DIA-NN export holds bare UniProt accessions with no species tag, so entrapment identifications must be recognised by membership in the set of entrapment accessions. A string prefix such as `ARATH`, which the FASTA headers might suggest, matches nothing and returns $\widehat{\text{FDP}}_{\text{entrap}} = 0$ without any error. The analysis script, and `diann_entrapment_ids()` in the package, therefore stop if no accepted row carries an entrapment accession.

##### S4.2 Which $r$, and which estimator

All corrected precursor-level estimates use $r = 0.432$, the peptide-level ratio with shared peptides removed, because competition takes place between precursors and not between proteins; the protein-group analysis of Note S4.3 also reports the value obtained with the protein-level ratio $r = 0.155$.

Throughout, $T_\alpha$ counts every accepted non-decoy match, entrapment matches included, so the accepted list has $T_\alpha$ members of which $E_\alpha$ are known to be false. Wen *et al.* [8] define three estimators on these counts. The raw proportion $E_\alpha/T_\alpha$ is a lower bound on the FDP of the *combined* target-plus-entrapment list. Their "combined" estimate, the one the main text adopts (its Equation (4)),

$$\widehat{\text{FDP}}^{\text{corr}}_{\text{entrap}}(\alpha) = \frac{E_\alpha(1 + 1/r)}{T_\alpha}, \tag{S10}$$

adds the expected number $E_\alpha/r$ of false matches that landed on real target sequences; they show that it overestimates the combined-list FDP on average, so it is read as an upper bound. Their "sample" estimate, $(E_\alpha/r)/(T_\alpha - E_\alpha)$, targets the FDP among the original targets only, the list an ordinary experiment reports, but is neither a lower nor an upper bound and typically underestimates. On the counts of Note S4.3 the combined and sample estimates give 0.719% and 0.503% for the global list and 1.29% and 0.90% for the $\min_r$(`Q.Value`) list; at protein level 2.20% and 1.54% (peptide-level $r$) or 4.94% and 4.31% (protein-level $r$). All verdicts in the paper use the raw value as the lower bound and Equation (S10) as the upper bound; `dfdr_entrapment()` returns all three (version 0.2.1; version 0.2.0 divided Equation (S10) by $T_\alpha + E_\alpha$, counting the entrapment matches twice, which moved the values in their third digit). All three are monotone in $E_\alpha/T_\alpha$ at fixed $r$, so any comparison between lists is the same under each; only the absolute values differ.

#### S4.3 Results

At $\alpha = 1\%$ the pure-entrapment rule gives $E_\alpha = 177$ among $T_\alpha = 81{,}643$ accepted targets on the `Global.Q.Value` list ($\widehat{\text{FDP}}_{\text{entrap}} = 0.217\%$, 95% Wilson interval $[0.187, 0.251]\%$; corrected 0.719%) and $E_\alpha = 342$ among $T_\alpha = 88{,}064$ on the $\min_r$(`Q.Value`) list (0.388%, $[0.349, 0.432]\%$; corrected 1.29%). Taking the minimum run-wise q-value therefore nearly doubles the proportion of entrapment matches. Following Wen *et al.* [8], we use the raw proportion as a lower-bound-like estimate (evidence of anti-conservative control if it exceeds $\alpha$) and the corrected one as an upper-bound-like estimate (consistent with control if it stays below $\alpha$), both under the assumptions of Wen *et al.*; homology and differential detectability can bias either in either direction, so neither is a guaranteed bound. Read this way, the global list is consistent with 1% control (0.719%), the $\min_r$ list is not (1.29%), and neither raw value indicates a failure.

**Clustered errors.** The Wilson intervals treat accepted precursors as independent, whereas entrapment matches cluster within protein groups (the 165 entrapment precursors of the discordant set belong to 4,350 added groups, the 18 at $\alpha = 10^{-4}$ to 14 groups). A bootstrap that resamples protein groups rather than precursors (2,000 draws; `05f_entrapment_cluster_bootstrap.R`; each accepted precursor is assigned to the `Protein.Group` string of its row, so every precursor belongs to exactly one cluster; in each draw the clusters of the set are sampled with replacement, all precursors of a drawn cluster are kept together, the entrapment proportion is recomputed on the resampled precursors, and for the ratio the two sets are resampled independently and the ratio is formed within each draw; intervals are the 2.5 and 97.5 percentiles) gives 0.217% $[0.180, 0.255]$ for the shared precursors and 2.57% $[2.17, 2.97]$ for the added ones at $\alpha = 1\%$, a ratio of 11.9 $[9.3, 14.9]$, and $[2.7, 9.0] \times 10^{-4}$ for the raw estimate at $\alpha = 10^{-4}$, which still excludes $10^{-4}$. The clustering therefore widens the intervals little on these data; it would matter more for a sample with fewer, larger protein groups, which is why the intervals are reported as descriptive. Protein groups are one clustering unit among several: peptides shared between groups and the six runs induce dependence that this bootstrap does not capture.

**The main report, the matrices and the number of runs.** DIA-NN's main report is filtered at `--qvalue` on the run-wise `Q.Value` and on the library q-value `Lib.Q.Value`, not on `Global.Q.Value`: in the deposited exports (`--qvalue 0.5`) the ceilings of `Q.Value` and `Lib.Q.Value` are both 0.5 and that of `Global.Q.Value` is 1 (Note S8), and the log reads "Output will be filtered at 0.5 FDR". In these two-pass searches `Lib.Q.Value` is the experiment-wide q-value of the precursor in the first pass (equal to the first-pass `Global.Q.Value` for all 165,538 target precursors present in both passes), and the second-pass library holds only the precursors below the first-pass ceiling, so the main report carries an implicit experiment-wide filter at `--qvalue`. What a user obtains by counting the unique precursors of the main report across runs at `Q.Value` $\leq 0.01$ therefore depends on the export: at the default `--qvalue 0.01` the library constraint is active and the list is the conservative `Q.Value` $\wedge$ `Lib.Q.Value` construction of Note S7.2 ($T = 85{,}197$, $D = 712$, 0.84% on the basic export); in a single-pass search against a predicted library, where `Lib.Q.Value` is zero, or in an export written at a liberal ceiling, as recommended to keep the decoys, the constraint is not binding at 1% and the same count gives the $\min_r$(`Q.Value`) list of main-text Table 3 (89,718 targets, 2,001 decoys). The effect grows with the number of runs, because each run gives an incorrect precursor another chance to pass: restricting the basic search to its first $k$ runs gives decoy ratios of 1.00%, 1.42%, 1.71%, 1.90%, 2.08% and 2.23% for $k = 1, \ldots, 6$ ($T_{0.01} = 69{,}968$, 79,972, 82,252, 86,499, 88,945, 89,718), so cohorts of tens or hundreds of runs will see a larger inflation than the one reported here. The matrices are different: DIA-NN's graphical interface and its `--matrix-qvalue` option filter each row on *both* `Q.Value` and `Global.Q.Value`. Script `09_double_filter.R` adds this construction to the scope audit: rows passing both filters at $\alpha$, collapsed to one entry per precursor.

On all three searches it returns exactly the `Global.Q.Value` list ($T$ = 83,149, $D$ = 832 on the basic search; $T$ = 81,643, $D$ = 816, $E$ = 177 on the entrapment search; Jaccard index 1 with the global list), because a precursor whose experiment-wide q-value passes $\alpha$ always has at least one run that passes it too. The matrices are therefore safe. The $\min_r$ list of the main text is what filtering on the run-wise column *alone* produces, without the library or experiment-wide constraint; of the ten DIA-NN studies that state a scope, two filter on `Q.Value` and `Lib.Q.Value` (Note S7.2).

**Scope-discordant set.** For two accepted lists $A_1$ and $A_2$ on a common identifier, entrapment matches are counted separately on the shared core $A_1 \cap A_2$, on the identifications added by $A_2$ ($A_2 \setminus A_1$) and on those it removes ($A_1 \setminus A_2$). Main-text Equation (5) pools the last two, which is appropriate only when one list contains the other; otherwise the added and removed sets should be reported separately, since a single rate can average an error-rich addition with an error-poor removal (`dfdr_entrapment_discordant()` returns all four). Here the global list is contained in the $\min_r$ list at every threshold of the grid (no precursor is removed), so $\widehat{\mathrm{FDP}}_{\mathrm{disc}}$ is the rate on the added set and $\widehat{\mathrm{FDP}}_{\mathrm{core}}$ the rate on the shared core. The added set is also the margin of a looser list: `Global.Q.Value` and $\min_r$(`Q.Value`) rank the target precursors almost identically (Spearman correlation 0.99), and at $\alpha$ = 1% and 5% the $\min_r$ list coincides exactly with the `Global.Q.Value` list at 2.19% and 10.1% on the entrapment search (2.23% and 10.6% on the basic search, 2.08% and 9.8% on the semi-tryptic one; at $\alpha = 10^{-3}$ the $\min_r$ list and the global list of the same size share 99.1–99.6% of their union, 98.6% at $5 \times 10^{-3}$ on the entrapment search). A relaxation of the global list to the same size therefore accepts the same precursors, and $\widehat{\mathrm{FDP}}_{\mathrm{disc}}$ measures the error of the margin that the mislabelled threshold adds (script `05d_discordant_size_matched.R`). At $\alpha$ = 1% the core holds 81,643 precursors with 177 entrapment matches (0.217%) and the added set 6,421 with 165 (2.57%, Wilson interval $[2.21, 2.99]$%), a ratio of 11.9. The ratio is 4.2, 3.9, 10.4, 11.9, 11.5 and 4.2 at $\alpha$ = 0.001, 0.002, 0.005, 0.01, 0.02 and 0.05, while the Jaccard index of the two lists stays between 0.91 and 0.94. Below $\alpha = 10^{-3}$ the added set holds too few entrapment matches (2 and 6 at $10^{-4}$ and $5 \times 10^{-4}$) for the ratio to mean much. These values come from `05b_entrapment_discordant.R`.

At protein-group level the same search gives $E_\alpha = 58$ among $T_\alpha$ = 8,751 on `Global.PG.Q.Value`: 0.663% raw ($[0.513, 0.856]$%), three times the precursor-level figure on the same data at the same nominal level, and 2.20% or 4.94% corrected, depending on whether the peptide-level or the protein-level $r$ is used. That range overlaps the 1.81–2.85% that Gu *et al.* [9] measured on an *in vitro*-expressed human proteome of known composition, and falls inside the broader 0.76–8.41% that Konno *et al.* [10] obtained from an entrapment search across three engines. The comparison is only descriptive, because instruments, engines, databases, entrapment designs and estimands all differ; its width is itself the reason to report $r$.

Across the threshold grid the comparison runs both ways. Over most of the grid the raw entrapment estimate sits *below* the decoy ratio, by a factor of five at $\alpha$ = 1%, and the upper-bound-like estimate stays below nominal at 1% (0.719%), consistent with conservative control at precursor level, the direction Gu *et al.* also report; a raw estimate alone could not show this. The order reverses below $\alpha \approx 10^{-3}$: at $\alpha = 10^{-4}$ the decoy ratio reads $1.2 \times 10^{-4}$ against a raw entrapment estimate of $5.4 \times 10^{-4}$ ($E_\alpha = 18$ among 33,484; Wilson interval $[3.4, 8.5] \times 10^{-4}$). Read as a lower-bound-like estimate, the raw value exceeds $\alpha$ and its interval excludes it, which by the rule above is evidence of anti-conservative control at $10^{-4}$, subject to the validity of the entrapment. This happens exactly where $D_\alpha$ becomes small ($D_\alpha = 4$ on the entrapment search), the regime that the stability diagnostics flag.

#### S5 Sensitivity to default parameters

Two defaults are used in the main text: the window width $w = 0.2$ that defines $D_{\alpha,\mathrm{win}}$, and the minimum bin occupancy $n_{\min} = 200$ together with the number of PEP bins that decide which bins enter the IPE. Neither has a theoretical justification, so both were recomputed over a grid on the two DIA-NN searches at $\alpha = 1\%$ (Table S6).

**Window width.** $D_{\alpha,\mathrm{win}}$ grows almost exactly linearly with $w$: on the basic global list 46, 93, 183 and 460 decoys for $w \in \{0.05, 0.1, 0.2, 0.5\}$, that is $920w$ to within 2%. The linearity shows that the decoy q-values are locally uniform near the cutoff, so no choice in this range falls into a gap, and the qualitative conclusion is the same at every setting on every list. Because the quantity scales with $w$, a reported $D_{\alpha,\mathrm{win}}$ means nothing without its window, which is why the checklist asks for both.

**PEP binning.** Over bin counts $\{5, 10, 20\}$ and $n_{\min} \in \{50, 100, 200, 500\}$ the IPE of the basic global list ranges from 0.0736 to 0.0755, a relative spread of 2.6%, and the flag is *caution* at all twelve settings; the semi-tryptic global list behaves the same (0.0687–0.0709); the run×precursor list gives $5.9$–$6.7\times10^{-5}$ and is *ok* throughout. The three orders of magnitude between the two scopes reported in the main text are therefore a property of the data, not of the binning. The grid does show where the estimator degrades: with 20 bins and $n_{\min} = 500$ only 16 of 20 bins survive on the basic global list, which is where most of the spread comes from. The default used throughout (bin width 0.05 restricted to PEP $\le 0.5$, $n_{\min} = 200$, nine usable bins) sits in the flat interior, and the number of bins actually used should be reported with any IPE.

#### S6 Ground-truth simulation

##### S6.1 Generative model

Each of $n$ units carries a target score $X_i$ and a decoy score $Y_i$. A fraction $\pi_0$ of the units are null: their two scores are independent draws from the same distribution $F_0$. The other units carry a correct match, whose target score is drawn from $F_0$ shifted by $\delta$. Three null families are used, Gaussian, Gumbel and Student-$t_4$, each standardised to mean 0 and variance 1 so that $\delta$ has the same meaning in all three. Unless stated otherwise the two scores of a unit compete: the unit enters the list with score $W_i = \max(X_i, Y_i)$ and is labelled target if $X_i > Y_i$ and decoy otherwise. For a null unit the two labels are then equally likely whatever the score, which is the equal-chance assumption. Q-values are computed by target–decoy counting: entries are sorted by decreasing score, the cumulative counts $T(t)$ and $D(t)$ are formed, $\widehat{\mathrm{FDR}}(t) = D(t)/\max(T(t), 1)$, and $q$ is the running minimum from the bottom of the list. Because the null status of every unit is recorded, the realised FDP of any accepted list is known exactly.

The model is implemented as `dfdr_simulate_tdc()` in `diagFDR` (since version 0.2.0), with `dfdr_realised_fdp()` for the realised FDP and `dfdr_check_cv_benchmark()` for the ratio $\kappa$ of Note S1.1; the script `simulations/07_package_reproduction.R` reproduces the S0–S3 and S5 rows of Table S4 from the package alone.

##### S6.2 Parameters and their relation to the data

We ran $B = 1000$ replicates of $n = 2 \times 10^5$ units at $\alpha \in \{5 \times 10^{-4}, 0.002, 0.01, 0.05\}$. We first tried to derive $\pi_0$ and $\delta$ from the DIA-NN export, but both derivations turned out to be fixed by the export settings rather than by the data, so we report them as design values:

- $\pi_0 = 0.667$. Under 1:1 competition with equal chance, the number of incorrect targets is estimated by the decoy count, so $\pi_0 \approx 2D_{\text{tot}}/(T_{\text{tot}} + D_{\text{tot}})$ over the full run×precursor export, here the `--qvalue 0.5` report. But at the bottom of an export capped at $q_{\max}$ the decoy ratio is $D_{\text{tot}}/T_{\text{tot}} \approx q_{\max}$, so the expression returns $2q_{\max}/(1 + q_{\max}) = 0.667$ for $q_{\max} = 0.5$ whatever the data (and about 0.02 for a report filtered at 1%). We therefore treat $\pi_0$ as a design choice that, together with $\delta$, gives accepted counts of the order of those in the DIA-NN data; Note S6.5 repeats the reference scenario over five $(\pi_0, \delta)$ pairs.
- $\delta = 4.65$. The export carries no `CScore` column, so $\delta$ had to be inferred from the accepted fraction at $\alpha = 1\%$ by solving $T_\alpha/\{T_{\text{tot}}(1 - \pi_0)\} = 1 - \Phi(t - \delta)$ with $t = z_{0.99}$ under the Gaussian model. On the run×precursor export ($T_{\text{tot}} = 680{,}203$, $D_{\text{tot}} = 340{,}262$) the left-hand side is $375{,}129/226{,}577 = 1.656$, above one because $\pi_0$ is set by the ceiling; the derivation clips it to 0.99, which gives $\delta = 2z_{0.99} = 4.65$ whatever the data. The value is therefore a design choice too: 99% of correct targets exceed the 1% upper quantile of the null. It gives $\mathbb{E}[D_{0.01}] = 659$ under S0, of the order of the 832 decoys of the DIA-NN global list, and should not be quoted as a measured property of the search engine.

#### S6.3 Scenarios and alerts

In S1 and S2 the decoy model is wrong. In S3, S4 and S5 the decoys are correct and something else changes: how the list is assembled (S3, S4) or the dependence between units (S5).

**S0, correct specification.** The model above, unchanged. It gives the rate at which each alert fires when nothing is wrong.

**S1, decoys too easy to beat.** Decoy scores are shifted by $-0.35$, so targets win null competitions more often than they should.

**S2, decoys under-dispersed.** Decoy scores are multiplied by 0.85, which keeps their centre and empties their upper tail.

**S3, enriched library without competition.** Targets and decoys are listed separately, $n$ of each, with no pairing, and the null fraction among targets is $\pi_0^{\text{lib}} = 0.45$ instead of $\pi_0$. This is the situation of a second search pass whose library has been reduced to the identifications of a first pass: fewer of the remaining targets are incorrect, while one decoy is still generated per target. Null targets and decoys have the same distribution, so nothing is wrong with the decoys. Second-pass scores are drawn afresh, so the scenario isolates the change in library composition from any reuse of first-pass evidence.

**S4, cross-run aggregation.** $k = 6$ independent runs of S0 on the same units are searched separately, and each unit keeps the smallest of its six run-level q-values. The unit is labelled a decoy if a decoy won in any run (S4a, decoy-if-any) or according to the run that supplied the minimum (S4b, winning row). S4a and S4b are the same simulated runs labelled by the two rules.

**S5, dependence between units.** Units are grouped in blocks of ten. Within block $g$, $X_i = \sqrt{\rho}\, b_g + \sqrt{1-\rho}\, e_i$ and $Y_i = \sqrt{\rho}\, c_g + \sqrt{1-\rho}\, f_i$ with all terms independent standard Gaussian, plus $\delta$ on the target of non-null units; $\rho \in \{0.2, 0.5, 0.8\}$. Because the block effects $b_g$ and $c_g$ are independent, equal chance holds exactly for every null unit: S5 changes the dependence between units and nothing else. Gaussian nulls only.

Four alerts are evaluated on each replicate, with the thresholds of main-text Table 4: $D_\alpha < 10$; decoy-support index $D_\alpha^{-1/2} > 0.30$; IPE $> 0.05$; and $|\widehat{\pi}_D - 0.5| > 0.15$, where $\widehat{\pi}_D$ is the decoy fraction among entries with $q \in [0.2, 0.5]$, at least 100 entries being required in that band. The first two thresholds are nearly equivalent ($D_\alpha^{-1/2} > 0.30$ means $D_\alpha \leq 11$) and are called the support alerts. The IPE needs the true PEP, which has a closed form only for competed Gaussian lists, so it is evaluated under S0, S1 and S2 with Gaussian nulls and marked "n/e" elsewhere.

A replicate is *bad* when its realised FDP exceeds $1.5\alpha$. The *sensitivity* of an alert is the proportion of bad replicates in which it fires; it is undefined ("—") where no replicate is bad. The rates in Table S4 are operating characteristics of one alert in one scenario: they depend on the null family, the threshold and the diagnostic, and are not error rates of the suite as a whole.

##### S6.4 What the simulation shows

Table S4 gives every cell at $\alpha = 1\%$.

*Nothing fires under correct specification, unless the level cannot be reached.* At $\alpha = 1\%$ no alert fires in any S0 replicate of any family. At $\alpha \leq 0.002$ the Gumbel and Student-$t_4$ lists collapse: the mean accepted decoy count is below 0.2, with about 150 accepted targets (Gumbel) or 2 (Student-$t_4$), so the nominal level is out of reach and the realised FDP (0.5% and 38%) is not a failure of control. The two support alerts fire in every such replicate, which is what they are for: they refuse to certify a degenerate list.

*The internal alerts do not see a wrong decoy model.* In the Gaussian S1 and S2 cells every one of the 1000 replicates is bad (realised FDP 2.59% and 2.83% for a nominal 1%) and the sensitivity of all four alerts is 0.00. The decoy ratio stays at 1.00% by construction, $D_\alpha$ stays near 680, the IPE stays at 0.009 and 0.010, and $\widehat{\pi}_D$ moves only from 0.5 to 0.428 and 0.531, inside the 0.15 tolerance. With heavier tails the same perturbations inflate the FDP less (1.32% and 1.85% with Gumbel nulls, 1.26% and 1.57% with Student-$t_4$) and are equally invisible.

*Cross-run aggregation: the alert fires on the harmless variant.* Under S4a the realised FDP is 0.18%, the equal-chance fraction is 0.98 and its alert fires in every replicate; the recomputed decoy ratio reads 11.5%, sixty times the realised error. Under S4b the realised FDP is 5.36%, the equal-chance fraction is 0.5 and no alert fires, although every replicate is bad. S4b is caught by the scope diagnostic: the decoy ratio recomputed on the aggregated list is 5.35% against a nominal 1%. The pattern is the same with Gumbel (0.08% against 2.45%) and Student-$t_4$ nulls (0.14% against 4.22%).

*Dependence leaves the mean alone and widens the spread.* Under S5 the mean realised FDP is 0.98% at every $\rho$ and no alert fires, but $\kappa = \mathrm{CV}(\mathrm{FDP})/\overline{D_\alpha^{-1/2}}$ rises from 1.41 under independence to 1.47, 1.80 and 2.51 at $\rho = 0.2$, 0.5 and 0.8 ($\alpha = 1\%$), and the share of replicates whose FDP exceeds 1% from 32% to 33%, 36% and 41%.

Two further quantities come from the same runs. First, under S0 with Gaussian nulls $\kappa$ is 1.46, 1.48, 1.41 and 1.41 at $\alpha = 5 \times 10^{-4}$, 0.002, 0.01 and 0.05: constant over two orders of magnitude in $\alpha$ and within Monte Carlo error of the $\sqrt{2} = 1.414$ that Proposition 1 predicts (main-text Equation (3)). Over the same cells the coefficient of variation of the *reported* decoy ratio falls from 0.010 to 0.00008, two to three orders of magnitude smaller, because the ratio is pinned to $\alpha$ by the step-up rule. Second, comparing the true PEP with the two candidate comparators over PEP $\leq 0.5$ gives IPE $= 0.0033$ for the pooled denominator $D_b/(T_b + D_b)$ against 0.0016 for $D_b/T_b$, a factor of 2.1 even under correct specification; on the real DIA-NN lists the same comparison spans a factor of 2.1 to 2.8 on the global lists and two orders of magnitude on the run×precursor lists. An IPE quoted without naming its comparator is therefore not comparable between papers.

##### S6.5 How stable is the variability factor

The values above come from one configuration: Gaussian nulls, $\pi_0 = 0.667$, $\delta = 4.65$. We therefore repeated S0 over five $(\pi_0, \delta)$ pairs around the reference values and all three null families, and recomputed $\kappa$ in each cell (Table S8; $B = 500$ replicates per cell, so each entry carries a Monte Carlo error of roughly 3%). A cell is evaluable when its mean accepted decoy count is at least 10.

Over the 38 evaluable cells of 60, $\kappa$ ranges from 1.35 to 1.74 with a median of 1.48. With Gaussian nulls it stays between 1.35 and 1.55; the Gumbel and Student-$t_4$ cells reach 1.74 and 1.66, because under heavier tails some accepted decoys displace correct targets and $D_\alpha$ varies more between replicates, so assumptions (iii) and (iv) of Proposition 1 hold less well (Note S1.1). Two conclusions follow. First, $D_\alpha^{-1/2}$ is a simple support index, and under the independent-label model the corresponding benchmark for $\mathrm{CV}(V_\alpha)$ is $\sqrt{2}D_\alpha^{-1/2}$; in every evaluable cell the realised variability exceeded $D_\alpha^{-1/2}$, although this does not establish a universal lower bound under arbitrary dependence. Second, where the margin between a reported FDP and a decision boundary is narrow enough for the difference between 1.4 and 1.7 to matter, the multiplier should be re-derived locally. Where the accepted decoy count is too small for the ratio to be estimated at all (the Gumbel and Student-$t_4$ cells at stringent $\alpha$), the list is in the granular regime itself and no multiplier helps.

The same five pairs show that the headline inflation figures do not depend on the choice of $(\pi_0, \delta)$: with Gaussian nulls at $\alpha = 1\%$ ($B = 200$), the mean realised FDP ranged from 2.44% to 2.70% under S1 and from 2.52% to 3.15% under S2, with no alert in any replicate.

##### S6.6 Does the result depend on the size of the misspecification

The two magnitudes of Note S6.3, a $-0.35$ shift and a $0.85\times$ scaling, were chosen by us. To see whether those cells are typical we swept the magnitude: shifts $s \in \{-0.10, -0.20, -0.35, -0.50, -0.70\}$ and scalings $c \in \{0.95, 0.90, 0.85, 0.80, 0.70\}$, under all three null families, with $B = 200$ replicates of $n = 2 \times 10^5$ at $\alpha = 1\%$ (Table S9).

Three things stand out. First, the realised FDP rises monotonically with the magnitude in both mechanisms and all three families, from about nominal at the smallest perturbations to $7.0\times$ nominal at $c = 0.70$ with Gaussian nulls ($4.0\times$ with Gumbel, $3.1\times$ with Student-$t_4$). The 2.6–2.8$\times$ of the reference cells is therefore a mid-range value on a continuum.

Second, no alert fires in 16 of the 18 cells whose mean realised FDP exceeds $1.5\alpha$ (28 of all 30 cells, the other 12 having little or nothing to detect). The support alerts cannot fire, because the accepted decoy count stays in the hundreds whatever the decoy model: decoy *abundance* carries no information about decoy *validity*. The equal-chance fraction leaves its tolerance in two cells only, the Gumbel shifts of $-0.50$ and $-0.70$ ($\widehat{\pi}_D = 0.35$ and 0.29, realised FDP $1.6\times$ and $2.2\times$ nominal). With Gaussian nulls it moves from 0.5 to 0.381 across the whole shift sequence and to 0.576 across the scaling sequence, both inside the tolerance, while the realised FDP reaches $6.0\times$ and $7.0\times$ nominal. The reason is clearest in the scaling arm: a narrowed decoy distribution keeps its centre, so the decoy share in the null-dominated band $q \in [0.2, 0.5]$ stays near one half, while the upper tail, the only region that matters at $\alpha = 1\%$, is depleted. The statistic looks at the wrong part of the distribution.

The count of undetected cells depends on the tolerance, which was set at 0.15 so that the check does not fire on the real first-pass export ($\widehat{\pi}_D = 0.55$, Note S8). Re-reading Table S9 with other tolerances: at 0.10, 14 of the 18 bad cells stay undetected and the largest undetected inflation is still $7.0\times$; at 0.05, eight stay undetected (all scalings of 0.80 or milder, and the Gaussian shift of $-0.20$), the largest undetected inflation falls to $3.9\times$, and four cells whose FDP is within $1.5\alpha$ raise an alert; at 0.03, two stay undetected ($2.0\times$) and six cells within $1.5\alpha$ fire. A tighter tolerance therefore buys sensitivity to the shift mechanism at the cost of alerts on valid lists, and buys little against the scaling mechanism, whose signature in the band is small whatever the tolerance; the main-text statement that the FDP reaches seven times nominal without an alert holds at the suite's tolerance of 0.15 and at 0.10, and becomes 3.9 times at 0.05.

Third, these mechanisms are simplifications. A global shift and a global scaling are the two cleanest ways to break equal chance, but real decoy misspecification is structured: reversed or shuffled decoys need not reproduce modification-site or missed-cleavage statistics, match-between-runs leaks identifications across runs, and shared peptides correlate at protein inference. The grid shows how the diagnostics behave under a controlled, monotone family of violations; it does not show that a real violation of comparable severity would be detected.

##### S6.7 An enriched library moves the equal-chance fraction with valid decoys

Scenario S3 addresses an observation of the main text: in real two-pass DIA-NN exports the decoy fraction among low-confidence precursors is 0.66–0.71, against 0.55 after a single pass. Under paired competition a null unit yields a decoy with probability 1/2, which is where the reference value 0.5 comes from. When targets and decoys are listed separately instead, a low-confidence entry is a decoy or a null target in the proportions in which the two are present, that is $1 : \pi_0^{\text{lib}}$, so the expected decoy fraction is $1/(1 + \pi_0^{\text{lib}})$ and exceeds 0.5 as soon as some library targets are correct.

Table S10 varies $\pi_0^{\text{lib}}$ with Gaussian nulls ($B = 200$, $\alpha = 1\%$). The equal-chance fraction follows $1/(1 + \pi_0^{\text{lib}})$ closely, from 0.53 at $\pi_0^{\text{lib}} = 0.9$ to 0.77 at 0.3, and its alert fires in every replicate once $\pi_0^{\text{lib}} \leq 0.5$. In none of these cells is the list anti-conservative: the decoy ratio reads 1.00% while the realised FDP is $\pi_0^{\text{lib}}$ times smaller. At the reference value $\pi_0^{\text{lib}} = 0.45$ the fraction is 0.69 for all three null families (Table S4).

A library whose targets are about 40–50% incorrect therefore suffices to produce the excess seen in the two-pass exports, and one with about 80% the single-pass value, with no failure of the null in either case. This shows that selection can account for the observation; it does not show that this is the mechanism at work in DIA-NN, whose second pass we have not modelled. It is the reason the equal-chance verdict is reported as *not evaluable* on two-pass exports. The same scenario predicts that the decoy ratio over-states the realised error, which is the direction of the entrapment comparison of Note S4.3.

##### S6.8 The sign flip between discrimination and support

Figure S1 plots the mean accepted decoy count against AUROC for $\delta \in [2, 5]$ (AUROC 0.921 $\to$ 0.9998; Gaussian nulls, 200 replicates per point) under the two operating rules. At a fixed nominal level $\alpha = 1\%$, $D_\alpha$ rises from 44 to 667 and $D_\alpha^{-1/2}$ falls from 0.150 to 0.039. At a fixed sensitivity $S^* = 0.6$ the same improvement drives $D_\alpha$ from 6763 to 0.2, passing through 98 at $\delta = 3.5$ and 5 at $\delta = 4.25$. The curves cross near $\delta \approx 3.1$. This is Propositions S1 and S2 in one picture: the sign of the relationship is set by the analyst's operating rule.

#### S7 The audit of reporting practice

##### S7.1 What the audit is, and what it can conclude

The audit covers 63 full-text articles from the 2024–2026 DIA proteomics literature. All are original research articles reporting bottom-up DIA data of their own, searched in the target–decoy framework, with the identification analysis described in the main text. Candidates were retrieved from PubMed (peer-reviewed research articles in English published 2024–2026; preprints, protocols, book chapters, corrections and reviews excluded), and an article entered the sample once its full text confirmed that it met these criteria. The sample is one of convenience. We make no claim to representativeness and give no confidence intervals for the proportions: establishing representativeness would require a sampling frame for a population with no defined boundary. Table S5 gives the composition, so a reader can judge the scope of the sample directly.

The audit motivates the study rather than being its subject; its purpose is to show that the quantities the diagnostics need are not in the published record. Two kinds of conclusion are drawn from it, and only one would need a representative sample. The proportions describe these 63 studies and nothing more, with three caveats. External validation was performed in seven studies, six of them methods or benchmark papers, so that figure reflects a sample skewed towards methodological work, where reporting tends to be more complete; the same applies to the proportion mentioning no identification error rate at all. The scores were produced in a single pass (a language model proposed each score with its supporting sentence, one author checked every score against the article, and a script verified every quote; Note S7.5); because each score is published next to the verbatim sentence that supports it, any reader can re-examine any judgement, and the two items on which the argument rests are presence-or-absence checks that need no interpretation. Finally, the audit measures what articles state: a study may have made a sound choice without documenting it.

The two conclusions the main text uses are null results, and a uniform absence is a different kind of claim from a proportion. Across 63 studies in 34 journals (40 applied, 15 methodological or technical, 8 benchmarks) using 21 distinct software combinations, none reported the accepted decoy count $D_\alpha$ and none stated a target/decoy label-propagation rule. One study scored *Partial* on the second point: it named the precursors entering the protein-group q-value but not how the label was assigned to the group, a choice that matters (Note S9).

Three features make this double absence insensitive to the caveats above. It is mechanical: whether a decoy count or a propagation rule appears is a presence-or-absence check. It is predictable: neither quantity appears in the default output of the major DIA engines (DIA-NN writes decoys only with `--report-decoys`), and neither is required by current reporting guidelines, so an article that omits them simply follows convention. And it is uniform across a heterogeneous sample: had either omission been confined to one journal or one software ecosystem, 34 journals and 21 software combinations would have exposed it. The audit therefore establishes a uniform absence in this sample, consistent with a reporting convention, without measuring its prevalence.

The ambiguity itself is not new: ten years ago The *et al.* [11] observed that protein-level FDR statements in shotgun proteomics were consistently imprecise, and set out what a report would need. What was missing was a measure of the gap and a way to close it from exports that laboratories can already produce, which is what the audit and the package supply.

##### S7.2 Scope-reporting practice among DIA-NN users

The scope mismatch of main-text Section 4.2 only matters if experiment-wide lists are in fact built from run-wise q-values. The scoring sheet answers this for the 36 studies that use DIA-NN: 26 state no scope at all, so a reader cannot tell which column was thresholded; of the ten that do (*Yes* or *Partial*), two (articles A29 and A50 of Data S1) filter the main report on the run-wise `Q.Value` and on `Lib.Q.Value`, the q-value of the precursor in the spectral library (constant across runs: one precursor in 290,461 has two distinct values in the basic export), without the experiment-wide `Global.Q.Value`; the others name a global column or the `--matrix-qvalue` double filter, which returns the global list (Note S4.3). The A29/A50 construction is not the $\min_r$ list: on the basic search `Q.Value` $\leq 0.01$ alone gives 89,718 targets and 2,001 decoys (2.23%), adding `Lib.Q.Value` $\leq 0.01$ gives 85,197 targets and 712 decoys (0.84%), because the library filter removes most of the low-confidence decoys that the run-wise filter admits, and is in fact the construction DIA-NN's default two-pass main report applies implicitly (Note S4.3). On the entrapment search the same construction gives $T = 83{,}949$, $E = 159$, a raw entrapment FDP of 0.189% (global list: 0.217%; $\min_r$: 0.388%) and a corrected value of 0.63%, so on these data it is conservative; its decoy ratio is nevertheless not a target–decoy estimate of anything, since the library filter is a selection of the kind Scenario S3 models (Note S6.7), and neither article gives the reader the means to know this. Across all 63 studies, eight state a scope and six name the column they thresholded.

#### S7.3 Criteria

Each article was scored from its main text only: an item given only in the supplementary material does not count, because it is the main text that carries the claim a reader meets. The scores refer to any identification analysis the article describes, including the search that builds a spectral library. Ten criteria were scored *Yes* or *No*, five of them also admitting *Partial* (1, 5, 6, 7 and 9): (1) the controlled unit is named with a numeric threshold; (2)–(4) a count of identifications is reported, under the three nested rules below; (5) the scope of the threshold is stated, run-wise or global; (6) the match-between-runs setting is stated; (7) the confidence column that was thresholded is named; (8) the number of accepted decoys $D$ is reported; (9) the target/decoy label-propagation rule is stated; (10) external validation, by entrapment or on samples of known composition, is performed. Descriptors were recorded beside them: whether an identification-level threshold is stated at all and its values at precursor or peptide and at protein level; whether a threshold for differential abundance is stated; whether two incompatible thresholds are given for the same list; whether a posterior error probability is used; whether the decoy generation strategy is named; whether the search is described as run with default settings; and the software and its version.

*Partial* was used when an article gave part of the information but not enough to reconstruct the claim: naming the controlled entity without a numeric threshold, or only through a column name; distinguishing two confidence values without saying which is run-wise and which global; discussing match-between-runs without giving its state in the analysis; giving a command-line flag or a filter setting without the column name. Listing thresholds at PSM, peptide and protein level is not a statement of scope. Partial scores are reported separately and never pooled with *Yes*. The full codebook is in Supplementary Data S1.

#### S7.4 Three nested rules for counting identifications

The counting criterion is the one where the scoring rule changes the figure the most, so three defensible rules are reported:

1. **Reported anywhere.** A count of identifications appears somewhere in the main text *and* a numeric identification-level threshold is stated somewhere; the two need not be linked.
2. **Reported together.** The count and the threshold appear in the same sentence, in two consecutive sentences, or in a table or caption introduced by the sentence stating the threshold, so the reader can see which filter produced which count.
3. **Of the controlled unit.** Rule 2 is met and the entity counted is one for which the article states a threshold (a precursor-level threshold accompanied by a count of precursors).

A count is a number of identified, detected or quantified PSMs, precursors, peptides, proteins or protein groups, given in the text, a table or a caption. Counts of differentially abundant entities do not qualify, since admitting them would let articles that state no identification threshold satisfy Rule 1 trivially; nor do numbers that can only be read from a plot. Ten articles report no count in this sense.

The drop from 84%, the proportion reporting a count irrespective of any threshold, through 73%, 19% and 16% under the three rules is the finding, and it is why the result cannot be reduced to one proportion. The information the reader needs is almost always present; what is missing is the link between its parts.

##### S7.5 How the scores were obtained and how to check them

The text of each article was extracted with `pdftotext`, in reading order and in physical layout, and sentences matching *false discovery*, *FDR*, *q-value*, *PEP*, *decoy*, *entrapment*, *match-between-runs*, *target–decoy* and the literal names of export columns were listed as candidates. That step locates sentences and assigns no score. A large language model (Claude Opus 5.5, Anthropic) then read the full text of each article and proposed a score for every item together with the sentence supporting it, and one author checked every proposed score against the article. Both the model and the checking author saw the full text, including author and journal names; no blinding was applied.

Each score other than *No* is published beside the sentence that supports it, quoted verbatim, in Supplementary Data S1, so any judgement can be re-examined against its source without re-reading the article. A script applies five checks to the scoring sheet: the DOI recorded for an article occurs in the text of that article; every quote occurs in the text of its article, compared after removal of spacing, punctuation, ligatures and line-end hyphenation; every score other than *No* carries a quote and every *No* carries none; the scores are logically consistent, the three count rules being nested; and no DOI occurs twice. All five pass on the published sheet. The same script computes every number of this Note and Tables S3 and S5 from the sheet. The two items on which the main text relies (criteria 8 and 9, both scored *No* for every article) are presence-or-absence checks: a reader who disagrees with any judgement can re-derive them from the quoted sentences, and a single counter-example would be visible in the sheet.

#### S8 Search and processing parameters

##### S8.1 DIA-NN searches

All three DIA-NN searches used version 2.3.1 (Academia build, compiled 5 December 2025) on the six ProteoBench LFQ Orbitrap AIF runs (`Condition_A_Sample_Alpha_01-03` and `Condition_B_Sample_Alpha_01-03`), 12 threads on an Intel Xeon Gold 6346 at 3.10 GHz with 127 GB RAM.

| Setting | Value |
|---|---|
| Sequence database | ProteoBench `MixedSpecies_HYE` FASTA |
| Spectral library | predicted *in silico* (`.predicted.speclib`), generated from the FASTA |
| Enzyme / missed cleavages | `--cut K*,R*`, `--missed-cleavages 1` |
| N-terminal Met excision | enabled (`--met-excision`) |
| Fixed modification | carbamidomethyl (C) (`--unimod4`) |
| Variable modifications | none |
| Peptide length | 8–25; precursor $m/z$ 300–1800; charge 2–4 |
| Fragment $m/z$ | 200–1800 |
| Mass accuracy | optimised automatically (14 ppm MS2; 11 ppm MS1 recommended on run 1) |
| Match between runs | enabled (`--reanalyse`, with `--rt-profiling`) |
| Decoy export | enabled (`--report-decoys`) |
| q-value export ceiling | `--qvalue 0.5` |

The three searches differ only as follows. The *basic* search uses the parameters above unchanged (wall clock 54 min; 1,020,465 report rows, 71 columns, 6 runs). The *semi-tryptic* search adds `--semi` (500 min, the expected cost of the larger search space). The *entrapment* search adds the reviewed *A. thaliana* proteome as a second `--fasta` and regenerates the predicted library over the combined database with `--gen-spec-lib` (74 min). The complete command lines are in the DIA-NN logs deposited with the data.

Two of these settings matter for the diagnostics. First, `--qvalue 0.5` is what makes the low-confidence diagnostics possible: the equal-chance band $q \in [0.2, 0.5]$ and the windowed support $D_{\alpha,\mathrm{win}}$ both require q-values above the reporting threshold, and a report filtered at 1% supports neither. The actual ceilings in the export are 0.5 for `Q.Value` and `Lib.Q.Value`, and 1 for `Global.Q.Value`, `PG.Q.Value` and `Global.PG.Q.Value`; because they differ by column, the ceiling must be read per column and not assumed; they also show that the main report is filtered at `--qvalue` on the run-wise `Q.Value` and on `Lib.Q.Value`, not on `Global.Q.Value` (the log reads "Output will be filtered at 0.5 FDR"), whereas the matrices apply the double filter of Note S4.3. In these two-pass exports `Lib.Q.Value` is the first-pass `Global.Q.Value` of the precursor (identical for all 165,538 target precursors present in both passes) and the second-pass library holds only the precursors below the first-pass ceiling, so the main report carries an implicit experiment-wide filter at `--qvalue` (Note S4.3). Second, `--reanalyse` makes these two-pass exports, which is why the equal-chance check is reported as *not evaluable*. DIA-NN also states in its log that protein inference is performed only for precursors with a global or run-wise q-value $\leq 0.05$, a selection step between the precursor and the protein level and one more reason why the two levels are not interchangeable.

#### S8.2 MaxQuant, MS$^2$Rescore and mokapot

The rescoring case study is a single 7-minute HeLa digest run, `20161213_NGHF_DBJ_SA_Exp3A_HeLa_1ug_7min_15000_02`, from the MS$^2$Rescore example set [12].

| Setting | Value |
|---|---|
| *MaxQuant / Andromeda* | |
| Version | 1.6.2.3 |
| Enzyme / missed cleavages | Trypsin/P, up to 2 |
| Fixed modification | carbamidomethyl (C) |
| Variable modifications | oxidation (M), acetyl (protein N-term), Gln→pyro-Glu, deamidation (NQ) |
| Precursor tolerance | 20 ppm first search, 4.5 ppm main search |
| Peptide length / mass | $\geq 7$ residues; $\leq 7500$ Da |
| Decoy construction | `revert` |
| Match between runs | disabled (single run) |
| PSM and protein FDR | set to 1, i.e. *unfiltered*: `msms.txt` retains all ranks and all decoys |
| *MS²Rescore v3.2.0* | |
| Feature generators | `basic` (10 features), MS²PIP (71 features), DeepLC (6 features); 87 in total |
| MS²PIP | model `HCD2021`, fragment tolerance 0.02 Da |
| DeepLC | no retraining, 20 epochs, calibration set fraction 0.15 |
| PSM ranks | input $\leq 10$, output $\leq 1$ |
| Spectrum key | `spectrum_id_pattern = ".*scan=(\d+)$"`, so `spectrum_id` is the MaxQuant scan number |
| Rescoring engine | mokapot, as bundled with MS²Rescore v3.2.0 |

MS²Rescore removed 4240 PSMs of rank > 10 and two with invalid amino acids, retaining 31,314 PSMs of which 35.96% are decoys; after rank-1 filtering 10,568 spectra remain. The exported mokapot q-values are target–decoy counts, $(D+1)/T$ with a running minimum, computed over all 31,314 candidate PSMs (up to ten per spectrum): recomputing them on that universe from the mokapot score reproduces every exported value within $6 \times 10^{-5}$, and thresholding them at 1% on it accepts 7987 targets and 78 decoys (0.98%). Applied to the 10,568 spectrum-level winners, the same q-values accept 7070 targets and 43 decoys (0.61%), whereas counting on the winners themselves accepts 7164 targets and 70 decoys (0.98%), the number MS²Rescore reports in its log. `mokapot_competed_universe()` in `diagFDR` offers both (argument `q_mode`). Two points matter for the diagnostics. The export carries no pre-rescoring score column, so the list before rescoring is rebuilt from `msms.txt` as described in Note S8.3. And because MaxQuant was run with its FDR thresholds set to 1, `msms.txt` is unfiltered, which is what makes the before/after comparison possible; it is the MaxQuant equivalent of DIA-NN's `--qvalue 0.5`.

This case study has no entrapment arm: a single HeLa digest offers no proteome known to be absent, so the external check is available only for the DIA-NN case study. This limitation comes from the dataset, not from the method.

#### S8.3 Rebuilding the list before rescoring

The mokapot export identifies a PSM by `run` and `spectrum_id`, whereas `msms.txt` identifies it by an internal integer `id`. A Jaccard index computed between accepted lists keyed on the one and on the other measures the key mismatch and not any change in identifications; it returns 0.05–0.10 whatever the data, so the before/after overlap curve must be computed on a key common to both exports.

The mapping is in fact exact. MS²Rescore was run with `spectrum_id_pattern = ".*scan=(\d+)$"`, so `spectrum_id` is the MaxQuant `Scan number` and `run` is the MaxQuant `Raw file`. Competing `msms.txt` to one winner per (`Raw file`, `Scan number`) yields exactly 10,568 spectra — the same number $MS^2$Rescore reports after rank-1 filtering — and the two competed lists share 100% of their identifiers. The overlap curve of main-text Figure 5E is computed on that key by `06b_msrescore_overlap.R`, which also recomputes and checks every accepted count quoted in main-text Section 4.3.

The before/after counts of main-text Section 4.3 are given on this competed list, so that both sides index the same 10,568 spectra: before rescoring $T = 5610$ and $D = 55$ at $\alpha = 1\%$, $T = 2138$ and $D = 0$ at $5 \times 10^{-4}$; after rescoring 7164 and 70 (counting on the mokapot score) or 7070 and 43 (mokapot q-values) at 1%, and 6013 and 5958 targets at $5 \times 10^{-4}$. The curve of main-text Figure 5D before rescoring is drawn on the uncompeted PSM-level list that `read_dfdr_maxquant_msms()` builds ($T = 5481$, $D = 53$ at 1%; 2139 and 0 at $5 \times 10^{-4}$), about 2% smaller at 1%. A Jaccard index between lists of different granularity has no clear meaning, which is why Figure 5E uses the competed list only.

#### S8.4 Software versions

The diagnostics were first run under R 4.3.3 on Windows 10 x64 (build 19045), 20 logical cores, with `diagFDR` 0.1.1 (0.1.2 for the report of main-text Figure 2B), `arrow` 22.0.0.1, `dplyr` 1.1.4, `ggplot2` 4.0.3 and `cp4p` 0.3.6. Version 0.2.1 of the package, the one cited in the main text, moves the label-propagation rules, picked-protein competition, the entrapment estimators and the simulator from the analysis scripts into the package (`collapse_universe()`, `dfdr_label_rule_audit()`, `dfdr_picked_protein()`, `dfdr_entrapment()`, `dfdr_entrapment_discordant()`, `dfdr_entrapment_size_ratio()`, `dfdr_simulate_tdc()`). Every number of main-text Table 3 and Section 4.2 and every entrapment value was recomputed with version 0.2.1 from the deposited exports; the scripts `03c_label_rules_package.R` and `05e_entrapment_package.R` in the code repository do this and stop if any value differs. Reading the three ≈100 MB parquet files is the only costly step; the diagnostics themselves run in seconds on the derived lists. Version 0.1.1 had one defect relevant to this work: when several rows were collapsed to one unit, the DIA-NN builders kept the minimum rather than the maximum of the `score` column. No number reported here is affected, because the DIA-NN exports carry no score column and every diagnostic used q-values and PEPs; the defect was fixed in 0.1.2.

**Python port.** `diagfdr` 0.1.0 (Python $\geq$ 3.9; PyPI and `https://github.com/Jacky11/diagfdr-py`) reimplements the diagnostics of Table S1 with the same function and column names (readers for DIA-NN, mokapot, MaxQuant and Spectronaut; the four label rules and picked-protein competition; the entrapment estimators, the discordant-set rate and the size ratio $r$; the alerts and the HTML report; the simulator). Its test suite compares every function with oracle values written by `diagFDR` 0.2.1 on the same tables, and reproduces from the deposited exports every number of Table 3, Section 4.2 and Section 4.3. The mzIdentML reader, the p-value diagnostics of Note S2 and the plots are not ported.

#### S8.5 Audit software

The audit of Note S7 used `pdftotext` (poppler-utils) for text extraction and Python 3 for sentence retrieval, the five verification checks and the tables, all implemented in `audit_pipeline.py`. The script and the scoring sheet (`scoring.csv`) are in the code repository (Data availability, main text); Supplementary Data S1 is generated from that sheet.

#### S8.6 Use of artificial intelligence

Two uses of a large language model (Claude Opus 5.5, Anthropic) are declared, as required by the journal. First, in the audit of Note S7, the model read the full text of each article and proposed, for every criterion, a score with the sentence supporting it; every score was checked by one author against the article and every quoted sentence was verified by script to occur verbatim in its source (Section S7.5). Second, the model was used to edit the wording of the manuscript and to check the consistency of the text, the numbers and the code against the deposited exports. No text, figure, table or reference was generated from a prompt; the authors wrote the manuscript and verified all content, for which they take full responsibility.

### S9 Target/decoy label propagation: the four rules

Search engines export several rows per reported unit: one row per run and precursor, and many precursor rows per protein group. Building the list a paper reports therefore means collapsing rows to units, which raises a question the reporting vocabulary has no word for: *if the rows being collapsed do not all carry the same target/decoy label, which label does the collapsed unit get?* This note defines the four possible answers, which are compared in main-text Table 3, Panel B. None of the audited studies says which one it used.

#### S9.1 Definitions

Let $k$ index the reported units and $R_k = \{1, \ldots, n_k\}$ the exported rows that map to unit $k$, row $j$ carrying q-value $q_{kj}$ and label $\ell_{kj} \in \{\mathrm{T}, \mathrm{D}\}$. Write $n_k^{\mathrm{D}} = \#\{j : \ell_{kj} = \mathrm{D}\}$ and call $k$ *label-mixed* when $0 < n_k^{\mathrm{D}} < n_k$. In all four rules the retained q-value is the group minimum $q_k = \min_j q_{kj}$, attained at row $j^*(k)$; the rules differ only in the label $\ell_k$ they assign, and in one case in how many units the collapse produces.

**`any_decoy` (decoy-if-any).** $\ell_k = \mathrm{D}$ if $n_k^{\mathrm{D}} > 0$. One unit per identifier. This is what a direct `any(is_decoy)` aggregation produces.

**`winning_row` (winning row).** $\ell_k = \ell_{kj^*(k)}$: the unit takes the label of the row that supplied its q-value; if a target row and a decoy row share the minimum, the decoy label is taken (the conservative choice; no such tie occurs in the exports analysed here). One unit per identifier. The rule a reader most often assumes.

**`majority`.** $\ell_k = \mathrm{D}$ if $n_k^{\mathrm{D}}/n_k > 1/2$, ties resolving to T. One unit per identifier.

**`split` (split target/decoy).** No label is propagated: the target rows and the decoy rows of an identifier are two distinct hypotheses, $(k, \mathrm{T})$ at $\min\{q_{kj} : \ell_{kj} = \mathrm{T}\}$ and $(k, \mathrm{D})$ at $\min\{q_{kj} : \ell_{kj} = \mathrm{D}\}$. A label-mixed identifier therefore gives one target unit *and* one decoy unit. On the DIA-NN export analysed here this is the construction that reproduces the protein-group q-values (decoy ratio 1.001% at `Global.PG.Q.Value` $\leq$ 1%): the decoy protein group behaves as a separate competing entity.

#### S9.2 Worked example

One protein-group identifier with four exported precursor rows, at $\alpha = 1\%$:

| row | 1 | 2 | 3 | 4 |
|---|---|---|---|---|
| $q$ | 0.004 | 0.02 | 0.30 | 0.60 |
| label | D | T | T | T |

| rule | units, labels | counted |
|---|---|---|
| `any_decoy` | 1 at $q = 0.004$, D | one decoy |
| `winning_row` | 1 at $q = 0.004$, D | one decoy |
| `majority` | 1 at $q = 0.004$, T | one target |
| `split` | 2 at 0.004 (D), 0.02 (T) | one decoy |

Two things are visible. `any_decoy` and `winning_row` agree here but for different reasons, and they diverge whenever the minimum-q row is a target while some other row is a decoy, which is the common case. And `majority` gives a target label to a unit whose q-value came from a decoy row, keeping the decoy's evidence while discarding its label, which is why we do not recommend it even though its values sit between the others.

##### S9.3 Invisible at one level, decisive at the next

Whether the rules can differ at all depends on the identifier: if no identifier is label-mixed the label is *intrinsic* and all four rules coincide exactly.

A DIA-NN `Precursor.Id` encodes the stripped sequence, the modifications and the charge of a library entry, and a decoy entry is a different sequence, so the label is intrinsic (exactly one of 290,461 precursors carries both) and the choice of rule makes no difference. A `Protein.Group` carries no decoy marking at all, because DIA-NN writes the same accession string for the target and the decoy precursors, so a label has to be assigned: 10,261 of the 13,750 protein-group identifiers of the basic search carry both labels somewhere in the `--qvalue 0.5` export.

On the protein list filtered at `Global.PG.Q.Value` $\leq 0.01$, the export a laboratory reporting a 1% protein-level FDR would normally keep, the four rules give:

| rule | $T_{0.01}$ | $D_{0.01}$ | ratio | $D_{0.01}^{-1/2}$ |
|---|---|---|---|---|
| `any_decoy` | 8,965 | 90 | 1.004% | 0.105 |
| `winning_row` | 8,991 | 64 | 0.712% | 0.125 |
| `majority` | 8,987 | 68 | 0.757% | 0.121 |
| `split` | 8,991 | 90 | 1.001% | 0.105 |

Of the 9,055 groups accepted at this threshold, 8,991 take their minimum q-value from a target row and 64 from a decoy row; 26 keep both a target and a decoy row. Every difference in the table comes from those 26 groups. `winning_row` counts them as targets, because their best evidence is a target; `any_decoy` counts them as decoys; `split` counts each of them twice, once on each side; `majority` divides them according to how many of their surviving rows are decoys. The spread in the reported ratio is a factor of 1.41, and the spread in the accepted decoy count is 64 to 90, a 41% change in the quantity that fixes both the reported error rate and, through the decoy-support index, its stability.

On the *unfiltered* export the rules no longer agree even approximately (`dfdr_label_rule_audit()` on the full `--qvalue 0.5` report, $\alpha = 1\%$):

| rule | units | $T_{0.01}$ | $D_{0.01}$ | ratio |
|---|---|---|---|---|
| `split` | 24,011 | 8,991 | 90 | 1.001% |
| `winning_row` | 13,750 | 8,991 | 64 | 0.712% |
| `any_decoy` | 13,750 | 963 | 8,092 | 840% |
| `majority` | 13,750 | 8,819 | 236 | 2.68% |

`split` and `winning_row` do not change, because the minimum-q row of a group is the same whatever the ceiling. `any_decoy` collapses: almost every group has at least one low-confidence decoy row somewhere in the export, so 8,092 of the 9,055 groups accepted at 1% become decoys and the list is destroyed. `majority` moves from 68 to 236 decoys. This is the practical reason never to apply `any(is_decoy)` to an export that has not been filtered first, and to state the ceiling of the export together with the rule. On the entrapment search, filtered at `Global.PG.Q.Value` $\leq 1\%$, the four rules give $T = 8{,}751$, 8,750,

8,733 and 8,750 targets with 88, 71, 88 and 71 decoys (ratios 1.006%, 0.811%, 1.008% and 0.811%) and 58, 58, 57 and 58 entrapment matches (raw entrapment FDP 0.65–0.66%): the rule changes the reported number, not the list.

On a filtered export the effect is modest beside the scope effects of main-text Table 3: it is an unstated degree of freedom of the same size as the difference between a 0.7% and a 1.0% claim. It is not a rounding artefact, because the groups it acts on are exactly those at the decision boundary, and a reader cannot detect it, because no paper states the rule. Two further remarks. `any_decoy` and `split` give the same $D_\alpha$ on the filtered export, differing only in whether the 26 mixed groups also count as targets, so on a filtered export the naive implementation happens to land close to the right answer, which is why a wrong rule goes unnoticed. And `majority` is the one rule whose value depends on how many low-confidence rows survived the filter, so it is a property of the export threshold as much as of the evidence.

##### S9.4 Recommendation

Report the rule, in the words of the checklist item of Table S2. Where the exporting tool treats decoy entities as separate competing hypotheses, as DIA-NN's protein-group q-values do on this export and as target–decoy competition at precursor level does in general, `split` is the rule consistent with how the q-values were computed, and it is what we use throughout. Where a genuine collapse to one unit per identifier is required, `winning_row` is the default we recommend, because the retained q-value and the retained label then come from the same row; the rules define different sets of hypotheses (`split` tests a target and a decoy hypothesis per identifier, the other three one per identifier), so when the software's construction is unknown the sensitivity to the four rules should be reported rather than one rule asserted. `any_decoy` should not be used on label-mixed identifiers, and never on an unfiltered export; `majority` is hard to interpret, because it can keep the q-value of a decoy row while giving the unit a target label and its value depends on how many low-confidence rows survived the export filter, so it should not be used unless that meaning is justified. All four are implemented in `collapse_universe()` of `diagFDR` (since 0.2.0), `dfdr_label_rule_audit()` returns the tables above in one call, and `dfdr_picked_protein()` gives the picked-protein comparator of main-text Section 4.2 (its Panel B row: $T = 9{,}146$, $D = 91$, ratio 0.995%). Picked-protein competition changes the list, not only the ratio: on the entrapment search it accepts 8,825 targets against 8,751 under `split`, and the 75 groups it adds contain 10 entrapment matches (13%; one group is dropped, with none), against 58 among the 8,750 shared groups (0.66%). The groups that picked competition recovers are the margin of a looser list, so `split`, which reproduces DIA-NN's construction, and picked competition, a different procedure, should not be presented as interchangeable. Because a filtered export can hide an inappropriate rule behind a reasonable-looking number, the check is worth running on the most liberal decoy-inclusive export available.

#### S10 Supplementary tables and figure

The tables are collected here so that their numbering does not depend on where they are discussed.

Table S1 maps the diagnostics to the package functions. Every diagnostic needs the exported decoys and a stated universe; the equal-chance bands and $D_{\alpha,\mathrm{win}}$ need q-values exported above the threshold (e.g. DIA-NN `--report-decoys --qvalue 0.5`); the IPE and $\sum$PEP need PEPs; the equal-chance check needs a single-pass export (`two_pass = TRUE` marks it not evaluable); external validation needs entrapment or ground-truth labels.

Table S1: Mapping between the diagnostics proposed in this work and the functions implemented in the `diagFDR` R package. Every number derived from the deposited exports was recomputed with v0.2.1 (Note S8); versions 0.1.1 and 0.1.2 were used during development and to draw the figures.

| Theme | What is checked or reported | Function(s) |
|---|---|---|
| **Input standardisation** | | |
| Universe definition | Standardise a target/decoy table (`id`, `is_decoy`, `q`; optional `pep`, `run`, `score`) and store the metadata that makes a claim auditable: `unit`, `scope`, `q_source`, `q_max_export`. | `as_dfdr_tbl()` |
| Tool adapters | Build competed universes, or reconstruct target–decoy competition q-values, from common tool outputs at PSM or precursor level. | `read_diann_parquet()`, `read_spectronaut()`, `read_mokapot_psms()`, `mokapot_competed_universe()`, `read_dfdr_maxquant_msms()`, `read_dfdr_mzid()` |
| DIA-NN scope-aware universes | The run×precursor universe (run-wise control), the deduplicated global precursor universe (global control), and the $\min_r$ aggregate as an explicit anti-pattern comparator. | `diann_runxprecursor()`, `diann_global_precursor()`, `diann_global_minrunq()`, `diann_global_proteingroup()` |
| Label propagation | Collapse exported rows to units under an explicit rule (`split`, `winning_row`, `any_decoy`, `majority`; Note S9); test whether the label is intrinsic; audit $T_\alpha$, $D_\alpha$ and the ratio under each rule; picked-protein competition as a procedural comparator. | `collapse_universe()`, `label_is_intrinsic()`, `dfdr_label_rule_audit()`, `dfdr_picked_protein()` |
| **Scope** | | |
| Scope disagreement | Jaccard overlap of accepted target sets across thresholds, pairwise or as a matrix at fixed $\alpha$. | `dfdr_scope_disagreement()`, `dfdr_plot_scope_disagreement_matrix()`, `dfdr_accepted()` |
| **External validation (A3)** | | |
| Entrapment FDP | Raw $E_\alpha/T_\alpha$ with Wilson interval, size-corrected estimate (main-text Equation 4) with the combined-list and target-only conventions (Note S4.2); $r$ at protein and peptide level from the FASTA files; attribution by accession set under the `pure` or `any` rule. | `dfdr_entrapment()`, `dfdr_entrapment_size_ratio()`, `read_fasta_accessions()`, `diann_entrapment_ids()`, `dfdr_is_entrapment_group()` |
| Scope-discordant set | Entrapment rate on the shared core, the added, the removed and the discordant identifications of two lists (main-text Equation 5). | `dfdr_entrapment_discordant()` |
| **Ground-truth simulation** | | |
| Competed lists with known truth | Generative model of Note S6 (three null families, decoy shift and scale, separate listing, block dependence), realised FDP, and the Monte Carlo ratio $\kappa$ against the benchmark $\sqrt{2}$. | `dfdr_simulate_tdc()`, `dfdr_realised_fdp()`, `dfdr_check_cv_benchmark()`, dataset `sim_tdc_example` |
| **Calibration** | | |
| Decoy-reference PEP concordance (IPE) | Bin by predicted PEP and compare the mean predicted PEP among targets with the competition-style comparator $\widehat{e}_b = \min\{1, (D_b + c)/T_b\}$, with $c$ the optional pseudo-count `add_decoy` (0 by default, giving main-text Equation (1)); summarise as the IPE (decoy-reference PEP concordance error). | `dfdr_ipe()`, `dfdr_pep_reliability_tdc()`, `dfdr_pep_reliability()` and plots |
| PEP sanity | Decoy PEP distribution (fraction of decoys with PEP $\leq 0.01/0.05/0.1/0.2$) and target-vs-decoy PEP densities; decoys should be stochastically larger. | `dfdr_pep_decoy_sanity()`, `dfdr_plot_pep_density_by_decoy()` |
| Absolute error burden | $\sum_{i \in A(\alpha)} \mathrm{PEP}_i$ and mean PEP among accepted targets across $\alpha$. | `dfdr_sumpep()` |
| Equal chance under paired competition | Decoy fraction by q-value band, with a pooled binomial test that it is near 0.5 in a mismatch-dominated region such as $q \in [0.2, 0.5]$. | `dfdr_equal_chance_qbands()`, `dfdr_plot_equal_chance()` |
| p-value workflows | Score-to-p-value conversion, super-uniformity plots, and Storey $\pi_0(\lambda)$ estimated separately from targets and decoys (Section S2). | `score_to_pvalue()`, `dfdr_plot_p_calibration()`, `dfdr_pi0_storey()` |

*Continued on next page*

Table S1 – continued

| Theme | What is checked or reported | Function(s) |
|---|---|---|
| **Stability** | | |
| Headline tail support | At $\alpha$: $T_\alpha$, $D_\alpha$, $D_\alpha/T_\alpha$, $D_\alpha^{-1/2}$ and the FDP-CV benchmark $\sqrt{2}D_\alpha^{-1/2}$, with $\pm k$ decoy-sensitivity displays. | `dfdr_headline()` |
| Support curves | $D_\alpha$ and the decoy-support index across a grid of $\alpha$, to expose granular regimes at stringent thresholds. | `dfdr_curve_stability()`, `dfdr_plot_dalpha()`, `dfdr_plot_cv()` |
| Local boundary support | $D_{\alpha,\mathrm{win}} = \#\{\text{decoys} : q \in (\alpha, (1+w)\alpha]\}$, with truncation handling when q-values are exported to a ceiling. | `dfdr_local_tail()`, `dfdr_plot_dwin()` |
| Threshold elasticity | $S_\alpha(\varepsilon) = J(A(\alpha), A((1+\varepsilon)\alpha))$, and the run-by-run Jaccard matrix of accepted targets. | `dfdr_elasticity()`, `dfdr_run_jaccard()` |
| BH workflows | BH threshold diagnostics and BH elasticity, for p-value-based pipelines. | `dfdr_bh_diagnostics()`, `dfdr_bh_elasticity()` |
| **Reporting** | | |
| Suite and report | Run the standard suite over a named list of `dfdr_tbl` objects and export tables, plots, manifest and an HTML report in which evaluable and non-evaluable alerts are distinguished and each alert is tagged by its evidential basis (derived check, simulation-informed alert, heuristic alert, descriptive; entrapment is external evidence; main-text Table 2). | `dfdr_run_all()`, `dfdr_write_report()`, `dfdr_render_report()`, `flag_headline()`, `dfdr_alert_basis()` |

Table S2: Comprehensive checklist for auditable reporting of "FDR = $\alpha$%". The abbreviated version is main-text Table 1.

| Theme | Item to report | Risk if missing | Example wording |
|---|---|---|---|
| **Object & scope** | Controlled object (PSM / precursor / peptide / protein / peak group) | "1% FDR" is not comparable across objects | *"FDR controlled at precursor level (DIA-NN row level)."* |
| | Reported object, with an explicit statement if it differs from the controlled object | Control at PSM level but proteins reported as if covered | *"Counts reported at precursor level; protein inference presented separately."* |
| | Scope: run-wise, global, or experiment-wide | Different q-values answer different questions | *"Selection on `Global.Q.Value` $\leq 0.01$ (global control)."* |
| | Whether match-between-runs was enabled | Transfer changes the effective universe after scoring; stated by 14 of the 63 audited studies | *"MBR enabled (`--reanalyse`); transfer across all six runs."* |
| | Target/decoy label-propagation rule when rows collapse to one unit | Four rules span a factor of 1.4 in the reported ratio and 41% in $D_\alpha$ at protein level on the same data (Section S9) | *"Target and decoy entries of a protein group are treated as distinct hypotheses; decoy-if-any is not used."* |
| **Method** | Method type: target–decoy competition, BH on p-values, mixture model | Assumptions differ (equal chance, p-value calibration) | *"Target–decoy competition; monotone q-values from $D/T$."* |
| | Exact formula ($D/T$, $(D+1)/T$, $2D/T$) and the level at which q-values are computed | Conventions change thresholds and comparisons; otherwise "1%" is ambiguous | "$\widehat{\mathrm{FDR}} = (D_\alpha + 1)/T_\alpha$*; computed globally after precursor deduplication."* |
| **Decoys** | Decoy strategy, ratio, library-based versus predicted | Equal chance and null representativeness may be violated | *"DIA-NN decoys (1:1), exported with `--report-decoys`."* |
| | Whether the report including decoys is available | Without decoys, calibration and stability cannot be verified | *"Full report with decoys on Zenodo [DOI]."* |
| | Export ceiling per exported field | Needed for the low-confidence diagnostics; ceilings differ between fields of one export | *"`--qvalue 0.5`; realised ceilings `Q.Value` = 0.5, `Lib.Q.Value` = 0.5, `Global.Q.Value` = 1."* |
| **Thresholds** | Primary $\alpha$ and any secondary values | Needed to diagnose cutoff sensitivity | *"Main results at 1%; diagnostics at 0.5, 1, 2, 5, 10%."* |
| | $T_\alpha$ *and* $D_\alpha$ at the headline threshold, in the same clause as the threshold | Without $D_\alpha$ no granularity diagnostic is possible (for q-values counted on the reported list, $D_\alpha \approx \alpha T_\alpha$ gives an approximation, which breaks when the q-values come from another universe or a model); reported by none of 63 audited studies | *"At 1%: $T_\alpha$ = 83,149, $D_\alpha$ = 832 (global)."* |

*Continued on next page*

Table S2 – continued

| Theme | Item to report | Risk if missing | Example wording |
|---|---|---|---|
| **Stability** | $D_\alpha$ across $\alpha$, and the decoy-support index $D_\alpha^{-1/2}$ or the FDP-CV benchmark $\sqrt{2}D_\alpha^{-1/2}$, *named* | Detects granular regimes; realised FDP variability is about $\sqrt{2}$ times the index | *"At 1%: $D_\alpha^{-1/2} = 0.035$, FDP-CV benchmark 0.049."* |
| | Local support $D_{\alpha,\mathrm{win}}$, *with the window stated* | A large $D_\alpha$ can hide a locally sparse boundary, and the value scales with the window | *"$D_{0.01,\mathrm{win}} = 183$ on $q \in (\alpha, 1.2\alpha]$."* |
| | Threshold elasticity, and inter-run overlap where relevant | Direct measure of list robustness | *"$S_{0.01}(0.2) = 0.98$; within-condition run overlap $> 0.85$."* |
| **Calibration** | IPE with the comparator named, and the number of bins used | The two candidate denominators differ by a factor of $\approx 2$ even under correct specification | *"IPE $= 0.075$ using $\widehat{e}_b = D_b/T_b$, 9 bins with PEP $\leq 0.5$, $n_b \geq 200$."* |
| | Equal-chance test under paired competition, with effect size, *and* whether the export is single- or two-pass | A two-pass export conditions the low-confidence region, so the test is not evaluable there | *"$\widehat{\pi}_D = 0.68$ on $q \in [0.2, 0.5]$; two-pass export, verdict* not evaluable.*"* |
| | $\sum$ PEP among accepted targets | Absolute error burden | *"At 1%: $\sum$ PEP $\approx 320$."* |
| **External** | $E_\alpha$ and $\widehat{\mathrm{FDP}}_{\mathrm{entrap}}$ with a descriptive Wilson interval (ignores clustering) | The principal check that does not rely on the same target/decoy labels | *"At 1%: $E = 177$, $\widehat{\mathrm{FDP}} = 0.217\%$ $[0.187, 0.251]\%$."* |
| | Counting rule, database size, and which $r$ any correction used | The corrected value depends on whether $r$ is measured at peptide or protein level: $r$ differs 2.8-fold here and the corrected value 2.2-fold | *"Pure entrapment; 16,418 accessions; corrected with peptide-level $r = 0.432$."* |
| **Reproducibility** | Software version, key parameters, and availability of configuration files | Calibration differs between releases; 7 of the 63 audited studies gave no version and 25 described the search only as run with default settings | *"DIA-NN v2.3.1; `--cut K*,R*`; scripts and reports on Zenodo [DOI]."* |
| **Limitations** | What the diagnostics do not guarantee | Avoids over-selling "1%" | *"Internal consistency is not ground truth; entrapment is sequence-specific. Post-processors can fail FDR control under spectrum multiplicity [13]."* |

Table S3: Reporting practice in the audited sample ($n = 63$). Percentages count *Yes* only; *Partial* scores are listed separately in Supplementary Data S1 and are never pooled with *Yes*, so the controlled-unit row would read 81% instead of 73% if partial disclosures were admitted. Rows 2–4 are the three nested count rules of Section S7.4; the collapse across them is the substantive result. Per-article scores with the verbatim supporting sentence are Supplementary Data S1.

| **Reporting item** | $n$/**63** | % |
|---|---|---|
| *The ten criteria* | | |
| 1. Controlled unit named (PSM, precursor, peptide, protein group) | 46 | 73 |
| 2. Identification count and threshold both reported, anywhere | 46 | 73 |
| 3. . . . reported together | 12 | 19 |
| 4. . . . and the count is of the controlled unit | 10 | 16 |
| 5. Scope stated (run-wise or global) | 8 | 13 |
| 6. Match-between-runs setting stated | 14 | 22 |
| 7. Thresholded confidence column named (e.g. `Global.Q.Value`) | 6 | 10 |
| 8. Accepted decoy count $D$ reported | 0 | 0 |
| 9. Target/decoy label-propagation rule stated | 0 | 0 |
| 10. External validation (entrapment or known composition) performed | 7 | 11 |
| *Descriptors* | | |
| Identification count reported, with or without a threshold | 53 | 84 |
| Numeric identification-level threshold stated | 53 | 84 |
| No identification-level threshold stated at all | 7 | 11 |
| of which a differential-abundance threshold is stated | 2 | 3 |
| Identification threshold stated, but none at protein level | 11 | 17 |
| Search software version given | 56 | 89 |
| Search described as default settings | 25 | 40 |
| Decoy generation strategy named | 13 | 21 |
| Posterior error probability used or reported | 1 | 2 |

Table S4: Ground-truth simulation at $\alpha = 1\%$: $B = 1000$ replicates of $n = 2 \times 10^5$ units, $\pi_0 = 0.667$, $\delta = 4.65$. Scenarios and alerts are defined in Note S6.3. FDP is the mean realised false discovery proportion, $D/T$ the mean reported decoy ratio, $\overline{D_\alpha}$ the mean accepted decoy count, $\widehat{\pi}_D$ the mean decoy fraction among entries with $q \in [0.2, 0.5]$, and "% bad" the share of replicates with FDP above $1.5\alpha$. The next four columns give the share of replicates in which each alert fires ("n/e": IPE not evaluated, the true PEP having no closed form), and the last the share of bad replicates in which at least one alert fires ("—" where no replicate is bad). Under S1, S2 and S4b the FDP exceeds nominal and no alert fires (the recomputed decoy ratio of main-text Table 4, step 2, is not part of the four alerts tabulated here; it would fire under S4a and S4b, whose ratios exceed $1.5\alpha$, and under no other scenario); under S3 and S4a the equal-chance alert fires although the FDP is below nominal.

| scenario | null family | FDP (%) | $D/T$ (%) | $\overline{D_\alpha}$ | $\widehat{\pi}_D$ | % bad | share of replicates flagged $D$ | $D^{-1/2}$ | IPE | eq. | sens. any |
|---|---|---|---|---|---|---|---|---|---|---|---|
| S0 | Gaussian | 0.98 | 1.00 | 659 | 0.500 | 0 | 0.00 | 0.00 | 0.00 | 0.00 | — |
| S0 | Gumbel | 0.75 | 1.00 | 368 | 0.500 | 0 | 0.00 | 0.00 | n/e | 0.00 | — |
| S0 | Student-$t_4$ | 0.82 | 1.00 | 605 | 0.500 | 0 | 0.00 | 0.00 | n/e | 0.00 | — |
| S1 | Gaussian | 2.59 | 1.00 | 678 | 0.428 | 100 | 0.00 | 0.00 | 0.00 | 0.00 | 0.00 |
| S1 | Gumbel | 1.32 | 1.00 | 609 | 0.390 | 1 | 0.00 | 0.00 | n/e | 0.00 | 0.00 |
| S1 | Student-$t_4$ | 1.26 | 1.00 | 643 | 0.410 | 0 | 0.00 | 0.00 | n/e | 0.00 | — |
| S2 | Gaussian | 2.83 | 1.00 | 680 | 0.531 | 100 | 0.00 | 0.00 | 0.00 | 0.00 | 0.00 |
| S2 | Gumbel | 1.85 | 1.00 | 659 | 0.535 | 100 | 0.00 | 0.00 | n/e | 0.00 | 0.00 |
| S2 | Student-$t_4$ | 1.57 | 1.00 | 653 | 0.526 | 82 | 0.00 | 0.00 | n/e | 0.00 | 0.00 |
| S3 | Gaussian | 0.45 | 1.00 | 1085 | 0.689 | 0 | 0.00 | 0.00 | n/e | 1.00 | — |
| S3 | Gumbel | 0.45 | 1.00 | 922 | 0.690 | 0 | 0.00 | 0.00 | n/e | 1.00 | — |
| S3 | Student-$t_4$ | 0.45 | 1.00 | 1042 | 0.688 | 0 | 0.00 | 0.00 | n/e | 1.00 | — |
| S4a | Gaussian | 0.18 | 11.46 | 7626 | 0.984 | 0 | 0.00 | 0.00 | n/e | 1.00 | — |
| S4a | Gumbel | 0.08 | 6.52 | 4246 | 0.984 | 0 | 0.00 | 0.00 | n/e | 1.00 | — |
| S4a | Student-$t_4$ | 0.14 | 11.15 | 7267 | 0.984 | 0 | 0.00 | 0.00 | n/e | 1.00 | — |
| S4b | Gaussian | 5.36 | 5.35 | 3768 | 0.500 | 100 | 0.00 | 0.00 | n/e | 0.00 | 0.00 |
| S4b | Gumbel | 2.45 | 2.65 | 1793 | 0.500 | 100 | 0.00 | 0.00 | n/e | 0.00 | 0.00 |
| S4b | Student-$t_4$ | 4.22 | 4.37 | 3036 | 0.500 | 100 | 0.00 | 0.00 | n/e | 0.00 | 0.00 |
| S5 ($\rho$ =0.2) | Gaussian | 0.98 | 1.00 | 659 | 0.500 | 0 | 0.00 | 0.00 | n/e | 0.00 | — |
| S5 ($\rho$ =0.5) | Gaussian | 0.98 | 1.00 | 659 | 0.500 | 0 | 0.00 | 0.00 | n/e | 0.00 | — |
| S5 ($\rho$ =0.8) | Gaussian | 0.98 | 1.00 | 659 | 0.500 | 0 | 0.00 | 0.00 | n/e | 0.00 | — |

Table S5: Composition of the audited sample ($n = 63$). This is a heterogeneous, non-representative set of studies from the 2024–2026 DIA proteomics literature (Section S7.1). Journals contributing a single study are pooled in the last row of the first block; software counts are mentions, so they sum to more than 63 because 24 studies use more than one software. The spread of this table is what makes the two zero-valued items of Table S3 informative.

| **Journal** | $n$ | **Article type** | $n$ |
|---|---|---|---|
| Journal of Proteome Research | 14 | Applied study | 40 |
| Molecular & Cellular Proteomics | 6 | Method or tool | 15 |
| Nature Communications | 5 | Benchmark | 8 |
| Analytical Chemistry | 4 | | |
| Proteomics | 4 | **Year** | |
| Journal of the American Society for Mass Spectrometry | 2 | 2024 | 9 |
| 28 further journals, one article each | 28 | 2025 | 24 |
| | | 2026 | 30 |
| | | **Acquisition** | |
| | | DIA only | 40 |
| | | DIA with a DDA, TMT or targeted arm | 23 |
| **Software (mentions)** | | **Totals** | |
| DIA-NN | 36 | Journals | 34 |
| Spectronaut | 29 | Distinct software combinations | 21 |
| FragPipe / MSFragger | 11 | Articles using more than one software | 24 |
| Proteome Discoverer | 5 | | |
| PaSER | 3 | | |
| MaxQuant | 3 | | |
| Mascot | 2 | | |
| PEAKS | 2 | | |
| 9 others, one mention each | 9 | | |

Table S6: Sensitivity of the two defaults, at $\alpha = 1\%$ on both DIA-NN searches. Upper block: the local window width $w$, with $D_{\alpha,\mathrm{win}}$ scaling essentially linearly, so no setting in this range lands in a gap and the verdict is *supported* throughout. "Supported" means the qualitative diagnostic conclusion was unchanged over the window widths evaluated; it does not define a universal cut-off. Lower block: the IPE over the $3 \times 4$ grid of bin counts $\{5, 10, 20\}$ and minimum occupancies $n_{\mathrm{min}} \in \{50, 100, 200, 500\}$, summarised as the range across the twelve settings with the alert each returned; the full grid is in `out/tables/S5_ipe_grid.csv`. The alert is constant within every universe, so neither default drives a conclusion.

| universe | value | | | alert |
|---|---|---|---|---|
| $D_{\alpha,\mathrm{win}}$ *by window width* $w = 0.05/0.1/0.2/0.5$ | | | | |
| basic global | 46 / 93 / 183 / 460 | | | supported |
| basic run × precursor | 218 / 430 / 863 / 2189 | | | supported |
| semi global | 44 / 87 / 175 / 452 | | | supported |
| | min | max | bins used | |
| *IPE across the* $3 \times 4$ *binning grid* | | | | |
| basic global | 0.0736 | 0.0755 | 5–19 | caution (12/12) |
| basic run × precursor | 0.000059 | 0.000067 | 5–20 | ok (12/12) |
| semi global | 0.0687 | 0.0709 | 4–17 | caution (12/12) |

Table S7: Symbols used in the main text, in order of first appearance, to be read alongside Section 3. Table S1 gives the corresponding `diagFDR` functions.

| **Symbol** | **Meaning** |
|---|---|
| $\alpha$ | Nominal FDR threshold at which a list is reported. |
| $A(\alpha)$ | The accepted set: exported rows retained at threshold $\alpha$. |
| $T_\alpha$, $D_\alpha$ | Accepted target and decoy counts in $A(\alpha)$. |
| $D_\alpha/T_\alpha$ | Decoy ratio recomputed on the reported list: a consistency audit of the exported column (A2), not an estimate of the FDP. |
| $J(X,Y)$ | Jaccard overlap $\lvert X\cap Y\rvert/\lvert X\cup Y\rvert$ of two accepted sets. |
| $J_{\mathrm{run/global}}(\alpha)$ | Overlap between the run-wise and global accepted lists at $\alpha$; reported descriptively. |
| $\mathrm{PEP}_i$ | Posterior error probability of unit $i$: $\Pr(\text{incorrect} \mid \text{target}, \text{score})$. |
| $T_b$, $D_b$ | Target and decoy counts falling in PEP bin $b$. |
| $\widehat{e}_b$ | Competition-style comparator for bin $b$, $\min\{1, D_b/T_b\}$; the estimand the mean predicted PEP in that bin should match. |
| $\overline{\mathrm{PEP}}_{b,T}$ | Mean predicted PEP among targets in bin $b$. |
| IPE | Decoy-reference PEP concordance error: weighted mean absolute gap between $\overline{\mathrm{PEP}}_{b,T}$ and $\widehat{e}_b$ over usable bins. |
| $n_{\min}$ | Minimum bin occupancy for a bin to enter the IPE (default 200). |
| $\widehat{\pi}_{D,j}$ | Decoy fraction within q-value band $j$; should approach 0.5 in mismatch-dominated bands if equal chance holds under paired competition. |
| $\mathbb{E}[V(\alpha)]$ | Expected number of false targets accepted, estimated by $\sum_{i\in A(\alpha)} \mathrm{PEP}_i$. |
| $D_\alpha^{-1/2}$ | Decoy-support index, the quantity the support alert thresholds (column `support_index` in `diagFDR`). The FDP-CV benchmark of Proposition 1 is $\sqrt{2}$ times larger; in every evaluable cell of Table S8 the realised FDP variability exceeded $D_\alpha^{-1/2}$ (Section S6.5). |
| $D_{\alpha,\mathrm{win}}$ | Local boundary support: decoys with $q \in (\alpha, (1+w)\alpha]$. Meaningless without $w$ stated. |
| $w$ | Relative width of that window (default 0.2). |
| $S_\alpha(\varepsilon)$ | Threshold elasticity $J(A(\alpha), A((1+\varepsilon)\alpha))$: list robustness to moving the cutoff. |
| $\delta$ | Discrimination parameter: location shift of correct targets. |
| $S^*$, $z^*$ | Target sensitivity held fixed in Proposition S1, and the corresponding standard-normal threshold $\Phi^{-1}(1-S^*)$. |
| $n_D$ | Total number of decoy scores in the universe. |
| $\pi_0$ | Proportion of null (incorrect) units among those tested. |
| $\pi_0^{\mathrm{lib}}$ | Proportion of null targets in the library of simulation scenario S3 (Section S6.7). |
| $\kappa$ | Ratio of the coefficient of variation of the realised FDP across replicates to the mean of $D_\alpha^{-1/2}$; Proposition 1 predicts $\sqrt{2}$ (Section S6.5). |
| $E_\alpha$ | Accepted identifications mapping exclusively to the entrapment proteome. |
| $\widehat{\mathrm{FDP}}_{\mathrm{entrap}}(\alpha)$ | Raw entrapment estimate $E_\alpha/T_\alpha$. |
| $E_\alpha^{-1/2}$ | Entrapment-support index, the analogue of $D_\alpha^{-1/2}$ for the external labels (main-text Figure 3D). |
| $r$ | Entrapment-to-target database size ratio, measured where competition occurs. |
| $\widehat{\mathrm{FDP}}^{\mathrm{corr}}_{\mathrm{entrap}}(\alpha)$ | Size-ratio-corrected entrapment estimate, $E_\alpha(1+1/r)/T_\alpha$. |
| $\widehat{\mathrm{FDP}}_{\mathrm{disc}}$, $\widehat{\mathrm{FDP}}_{\mathrm{core}}$ | Raw entrapment estimate restricted to the identifications on which two accepted lists disagree (symmetric difference $A_1 \triangle A_2$, identifications accepted by one list only) or that they share ($A_1 \cap A_2$); main-text Equation (5). |

Table S8: Stability of the FDP-variability factor $\kappa = \mathrm{CV}(\mathrm{FDP})/\overline{D_\alpha^{-1/2}}$ under correct specification (S0), over five $(\pi_0, \delta)$ pairs and three null families ($B = 500$ replicates of $n = 2 \times 10^5$ per cell; Monte Carlo error $\approx 3\%$ per entry). "—" marks cells whose mean accepted decoy count is below 10, which is the granular regime itself. Over the 38 evaluable cells $\kappa$ lies between 1.35 and 1.74, median 1.48, against the $\sqrt{2} = 1.414$ of Proposition 1. The reference cell ($\pi_0 = 0.667$, $\delta = 4.65$) is an independent run of 500 replicates and differs from the 1000-replicate values of Note S6 by Monte Carlo error.

| null family | $\pi_0$ | $\delta$ | $\alpha = 5 \times 10^{-4}$ | 0.002 | 0.01 | 0.05 |
|---|---|---|---|---|---|---|
| Gaussian | 0.550 | 5.50 | 1.45 | 1.37 | 1.44 | 1.35 |
| Gaussian | 0.600 | 4.00 | 1.49 | 1.51 | 1.45 | 1.38 |
| Gaussian | 0.667 | 3.50 | 1.52 | 1.55 | 1.48 | 1.47 |
| Gaussian | 0.667 | 4.65 | 1.37 | 1.37 | 1.48 | 1.50 |
| Gaussian | 0.750 | 5.00 | 1.48 | 1.46 | 1.38 | 1.39 |
| Gumbel | 0.550 | 5.50 | — | 1.65 | 1.54 | 1.47 |
| Gumbel | 0.600 | 4.00 | — | — | — | 1.59 |
| Gumbel | 0.667 | 3.50 | — | — | — | 1.71 |
| Gumbel | 0.667 | 4.65 | — | — | 1.74 | 1.45 |
| Gumbel | 0.750 | 5.00 | — | — | 1.59 | 1.42 |
| Student-$t_4$ | 0.550 | 5.50 | — | — | 1.43 | 1.39 |
| Student-$t_4$ | 0.600 | 4.00 | — | — | 1.41 | 1.54 |
| Student-$t_4$ | 0.667 | 3.50 | — | — | — | 1.56 |
| Student-$t_4$ | 0.667 | 4.65 | — | — | 1.66 | 1.51 |
| Student-$t_4$ | 0.750 | 5.00 | — | — | 1.53 | 1.41 |

Table S9: Realised FDP and alert sensitivity across a grid of decoy-misspecification magnitudes, at $\alpha = 1\%$ ($B = 200$ replicates of $n = 2 \times 10^5$ per cell). $\overline{D_\alpha}$ is the mean accepted decoy count and $\widehat{\pi}_D$ the mean equal-chance fraction. A replicate is *bad* when its realised FDP exceeds $1.5\alpha$; the last column is the share of bad replicates in which at least one alert fires ("—" where no replicate is bad). The only cells in which an alert fires are the Gumbel shifts of −0.50 and −0.70, through the equal-chance alert. Monte Carlo error on each FDP is about 0.4% relative.

| **S1: decoy scores shifted by $s$** | | | | | | | |
|---|---|---|---|---|---|---|---|
| Null family | $s$ | realised FDP | × nominal | $\overline{D_\alpha}$ | $\widehat{\pi}_D$ | % bad | sens. any flag |
| Gaussian | −0.10 | 1.31% | 1.31 | 664 | 0.478 | 0 | — |
| Gaussian | −0.20 | 1.73% | 1.73 | 669 | 0.456 | 100 | 0.00 |
| Gaussian | −0.35 | 2.58% | 2.58 | 678 | 0.427 | 100 | 0.00 |
| Gaussian | −0.50 | 3.78% | 3.78 | 688 | 0.403 | 100 | 0.00 |
| Gaussian | −0.70 | 6.04% | 6.04 | 707 | 0.381 | 100 | 0.00 |
| Gumbel | −0.10 | 0.89% | 0.89 | 471 | 0.468 | 0 | — |
| Gumbel | −0.20 | 1.04% | 1.04 | 541 | 0.436 | 0 | — |
| Gumbel | −0.35 | 1.31% | 1.31 | 608 | 0.390 | 0 | 0.00 |
| Gumbel | −0.50 | 1.65% | 1.65 | 646 | 0.345 | 96 | 0.99 |
| Gumbel | −0.70 | 2.21% | 2.21 | 671 | 0.290 | 100 | 1.00 |
| Student-$t_4$ | −0.10 | 0.92% | 0.92 | 619 | 0.470 | 0 | — |
| Student-$t_4$ | −0.20 | 1.04% | 1.04 | 630 | 0.443 | 0 | — |
| Student-$t_4$ | −0.35 | 1.25% | 1.25 | 643 | 0.410 | 0 | — |
| Student-$t_4$ | −0.50 | 1.52% | 1.52 | 652 | 0.396 | 60 | 0.00 |
| Student-$t_4$ | −0.70 | 1.97% | 1.97 | 662 | 0.443 | 100 | 0.00 |
| **S2: decoy scores multiplied by $c$** | | | | | | | |
| Null family | $c$ | realised FDP | × nominal | $\overline{D_\alpha}$ | $\widehat{\pi}_D$ | % bad | sens. any flag |
| Gaussian | 0.95 | 1.41% | 1.41 | 666 | 0.509 | 8 | 0.00 |
| Gaussian | 0.90 | 2.03% | 2.03 | 673 | 0.519 | 100 | 0.00 |
| Gaussian | 0.85 | 2.84% | 2.84 | 680 | 0.531 | 100 | 0.00 |
| Gaussian | 0.80 | 3.88% | 3.88 | 689 | 0.544 | 100 | 0.00 |
| Gaussian | 0.70 | 6.98% | 6.98 | 714 | 0.576 | 100 | 0.00 |
| Gumbel | 0.95 | 1.05% | 1.05 | 543 | 0.511 | 0 | — |
| Gumbel | 0.90 | 1.41% | 1.41 | 622 | 0.522 | 9 | 0.00 |
| Gumbel | 0.85 | 1.85% | 1.85 | 658 | 0.535 | 100 | 0.00 |
| Gumbel | 0.80 | 2.40% | 2.40 | 676 | 0.548 | 100 | 0.00 |
| Gumbel | 0.70 | 3.98% | 3.98 | 693 | 0.579 | 100 | 0.00 |
| Student-$t_4$ | 0.95 | 1.01% | 1.01 | 628 | 0.507 | 0 | — |
| Student-$t_4$ | 0.90 | 1.27% | 1.27 | 643 | 0.516 | 0 | — |
| Student-$t_4$ | 0.85 | 1.57% | 1.57 | 653 | 0.526 | 78 | 0.00 |
| Student-$t_4$ | 0.80 | 1.94% | 1.94 | 662 | 0.538 | 100 | 0.00 |
| Student-$t_4$ | 0.70 | 3.09% | 3.09 | 676 | 0.566 | 100 | 0.00 |

Table S10: Scenario S3 over a range of library compositions: targets and decoys listed separately, null fraction $\pi_0^{\text{lib}}$ among targets, Gaussian nulls, $\alpha = 1\%$ ($B = 200$). The decoys are valid in every row. The equal-chance fraction $\widehat{\pi}_D$ follows $1/(1 + \pi_0^{\text{lib}})$, and the decoy ratio over-states the realised FDP by $1/\pi_0^{\text{lib}}$. The last column is the share of replicates in which the equal-chance alert fires.

| $\pi_0^{\text{lib}}$ | realised FDP | decoy ratio | ratio / FDP | $\widehat{\pi}_D$ | eq.-chance flag |
|---|---|---|---|---|---|
| 0.900 | 0.90% | 1.00% | 1.10 | 0.525 | 0.00 |
| 0.800 | 0.81% | 1.00% | 1.24 | 0.555 | 0.00 |
| 0.667 | 0.67% | 1.00% | 1.50 | 0.599 | 0.00 |
| 0.500 | 0.50% | 1.00% | 2.00 | 0.666 | 1.00 |
| 0.450 | 0.45% | 1.00% | 2.21 | 0.690 | 1.00 |
| 0.400 | 0.40% | 1.00% | 2.48 | 0.714 | 1.00 |
| 0.300 | 0.30% | 1.00% | 3.33 | 0.769 | 1.00 |

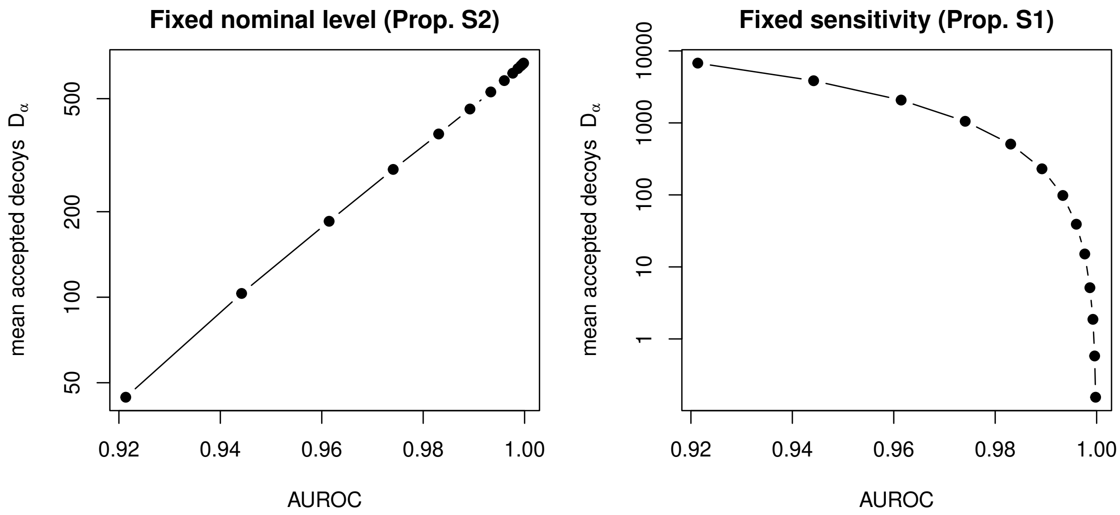

Figure S1: The discrimination–support sign flip. Mean accepted decoy count $D_\alpha$ against AUROC as the separation $\delta$ increases from 2.0 to 5.0 (AUROC 0.921 to 0.9998), under two operating rules; each point is the mean of 200 replicates. At a fixed nominal level $\alpha = 1\%$ (Proposition S2) improving discrimination *increases* decoy support, from $D_\alpha = 44$ to 667, because $D_\alpha$ is pinned to $\alpha T_\alpha$ and $T_\alpha$ grows. At a fixed target sensitivity $S^* = 0.6$ (Proposition S1) the same improvement destroys it, from 6763 to 0.2, because the operating threshold is pushed into the tail. The curves cross near $\delta \approx 3.1$. Gaussian nulls, $n = 2 \times 10^5$, $\pi_0 = 0.667$; specification in Note S6.

#### References

[1] Arya Ebadi, Dong Luo, Jack Freestone, William Stafford Noble, and Uri Keich. Bounding the FDP in competition-based control of the FDR, 2023. arXiv:2302.11837; R package `bandsfdp` on CRAN.

[8] Bo Wen, Jack Freestone, Michael Riffle, Michael J MacCoss, William S Noble, and Uri Keich. Assessment of false discovery rate control in tandem mass spectrometry analysis using entrapment. *Nature Methods*, 2025.

[9] Kongxin Gu, Masanaga Kenko, Koji Ogawa, Naoki Goshima, Takeshi Masuda, Shingo Ito, and Sumio Ohtsuki. Evaluation of the false discovery rate in library-free search by DIA-NN using in vitro human proteome. *Journal of Proteome Research*, 24(8):3874–3883, 2025.

[10] Ryo Konno, Masaki Ishikawa, Daisuke Nakajima, Kaori Inukai, Osamu Ohara, and Yusuke Kawashima. Thin-diaPASEF: diaPASEF for maximizing proteome coverage in single-shot proteomics. *DNA Research*, 32(4):dsaf019, 2025.

[11] Matthew The, Ayesha Tasnim, and Lukas Käll. How to talk about protein-level false discovery rates in shotgun proteomics. *Proteomics*, 16(18):2461–2469, 2016.

[12] CompOmics. Example input files for $MS^2$Rescore. Zenodo, January 2024.

[13] Jack Freestone, William Stafford Noble, and Uri Keich. Reinvestigating the correctness of decoy-based false discovery rate control in proteomics tandem mass spectrometry. *Journal of proteome research*, 23(6):1907–1914, 2024.
